# Long-term synaptic depression triggers local biogenesis of autophagic vesicles in dendrites and requires autophagic degradation

**DOI:** 10.1101/2020.03.12.983965

**Authors:** Emmanouela Kallergi, Akrivi-Dimitra Daskalaki, Evangelia Ioannou, Angeliki Kolaxi, Maria Plataki, Per Haberkant, Frank Stein, Mikhail M Savitski, Kyriaki Sidiropoulou, Yannis Dalezios, Vassiliki Nikoletopoulou

**Affiliations:** Department of Fundamental Neurosciences, University of Lausanne, Lausanne, 1005, Switzerland; Medical School, University of Crete, Greece; Proteomic Core Facility (PCF), European Molecular Biology Laboratory (EMBL), Heidelberg, Germany; School of biological sciences, University of Crete, Greece; Genome Biology Unit, European Molecular Biology Laboratory (EMBL)

## Abstract

In neurons, biogenesis of autophagic vesicles (AVs) is spatially confined to the axon tip under baseline conditions. However, it remains unknown whether their biogenesis can be induced in other neuronal compartments following synaptic activity in order to serve local functions. Here, we show that both major types of long-term synaptic depression (LTD), a form of plasticity expressed by the shrinkage and elimination of dendritic spines, trigger the rapid and local biogenesis of AVs in post-synaptic dendrites. In return, autophagy is indispensable for LTD, as either genetic ablation of *atg5* in pyramidal neurons or acute pharmacological inhibition of AV biogenesis totally prevents LTD induction. Using quantitative proteomic profiling of purified AVs, we reveal that upon LTD the autophagic cargo is significantly enriched for synaptic proteins, as well as modulators of the actin cytoskeleton and autism-implicated proteins. In line with these findings, a mild autophagy deficit is sufficient to impair behavioral flexibility, a cognitive function that requires efficient LTD. Therefore, local synthesis and assembly of the autophagic machinery in dendrites ensure the elimination of synaptic structures via degradation of their components, facilitating plasticity and associated behaviors.

**Graphical Abstract:** 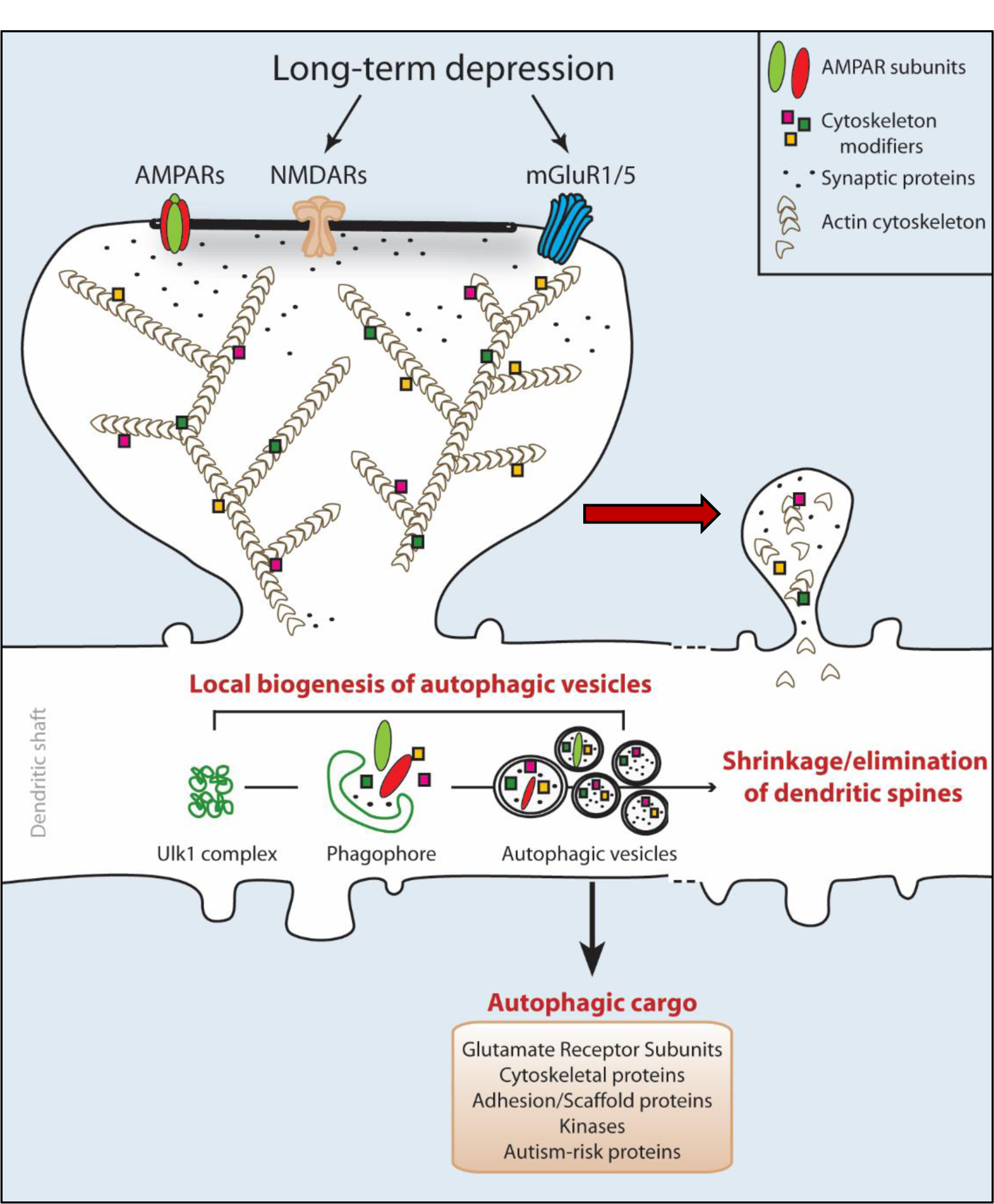

**In brief:** Kallergi, Daskalaki and colleagues demonstrate that autophagy is cell autonomously required in pyramidal excitatory neurons for the induction of long-term synaptic depression (LTD). They uncover the novel and local biogenesis of autophagic vesicles (AVs) in dendrites upon LTD, by which post-synaptic components are rapidly accessible on-site for autophagic degradation. Using quantitative proteomics on purified AVs, they reveal that upon LTD the autophagic cargo is enriched in synaptic, cytoskeletal and autism-implicated proteins.

**Highlights:**

- Autophagy is required cell-autonomously in pyramidal neurons for LTD.
- NMDAR- and mGluR-mediated LTD trigger the local biogenesis of autophagic vesicles in dendrites.
- Autophagic vesicles sequester primarily synaptic and cytoskeletal cargo upon LTD.
- Mild impairment in autophagy leads to deficits in cognitive flexibility.

## Introduction

Macroautophagy (hereafter referred to as autophagy) is a highly conserved mechanism that sequesters cellular constituents in autophagic vesicles (AVs) and delivers them to the lysosome for degradation. Biogenesis of AVs is a multistep process that entails the nucleation of a cup-shaped isolation membrane (also known as the phagophore), which sequesters cargo as it elongates and eventually closes to form the complete, double-membrane bound AV, as previously reviewed (Lamb et al., 2013; Shibutani and Yoshimori, 2014).

Though initially considered as a process in bulk, it is increasingly appreciated that autophagy can also serve specific functions, such as the regulation of food intake in hypothalamic neurons (Aveleira et al., 2015; Kaushik et al., 2011; Oh et al., 2016; Xiao et al., 2017). In line with this notion, the contribution of autophagy to higher brain functions and to the underlying mechanisms of synaptic plasticity is beginning to be addressed (Glatigny et al., 2019; Liang, 2019; Lieberman et al., 2020; Nikoletopoulou and Tavernarakis, 2018). For example, autophagy was recently shown to participate in memory erasure by synaptic destabilization (Shehata et al., 2018). Moreover, our previous work indicated that neuronal autophagy is negatively regulated by brain-derived neurotrophic factor (BDNF), a key regulator of synaptic plasticity, and that suppression of autophagy can rescue the synaptic defects associated with BDNF deficiency (Nikoletopoulou et al., 2017).

In contrast to most cell types, where AV biogenesis occurs indiscriminately throughout the cytoplasm, previous work demonstrated that in diverse neuronal populations, including hippocampal pyramidal neurons, biogenesis of AVs is spatially confined to the axon tip under baseline conditions (Maday and Holzbaur, 2016; Maday et al., 2012). In neurons, nascent phagophores mature to complete AVs as they are retrogradely transported via dynein motors from the axon tip towards the soma, where they fuse with lysosomes to deliver their cargo for degradation (Fu and Holzbaur, 2014a, b; Fu et al., 2014; Hollenbeck, 1993; Katsumata et al., 2010).

While in the pre-synaptic side autophagy has been shown to regulate neurotransmitter release directly by degrading synaptic vesicles (Binotti et al., 2015; Hernandez et al., 2012; Luningschror et al., 2017; Soukup et al., 2016), as reviewed previously (Luningschror and Sendtner, 2018; Nikoletopoulou and Tavernarakis, 2018; Vijayan and Verstreken, 2017), its role in regulating post-synaptic processes is less understood. Recent findings suggests that autophagy can facilitate the degradation of AMPA (Shehata et al., 2012) and GABA receptors (Rowland et al., 2006). In addition, it was shown to be cell autonomously required in excitatory neurons for developmental spine pruning (Tang et al., 2014), a process suggested to be mediated by mechanisms similar to long-term depression (LTD) of synaptic strength (Piochon et al., 2016).

However, the requirement of autophagy in LTD, a major form of synaptic plasticity underlying key cognitive functions across a broad range of species (Brigman et al., 2010; Dalton et al., 2008; Nabavi et al., 2014), remains elusive. Moreover, as this form of plasticity primarily involves the degradation of post-synaptic components and structures, two key questions arise regarding a) whether dendritic proteins are degraded by autophagy and b) how these dendritic constituents become timely accessible to neuronal AVs, which normally route between the axon and the soma. In this study, we addressed these questions, by investigating the interplay and interdependence between LTD and autophagy in pyramidal neurons.

## Results

### NMDAR- and mGluR-LTD trigger the rapid appearance of autophagic structures in dendrites of cultured neurons

Under baseline conditions, AV biogenesis is confined to the axon tip of cultured neurons and AVs are retrogradely transported to the soma to fuse with lysosomes (Maday and Holzbaur, 2016; Maday et al., 2012). We sought to examine whether AV distribution and biogenesis are altered by LTD. To this end, the two major types of LTD that coexist in the brain, NMDAR- and mGluR-LTD, were chemically induced in cultured hippocampal neurons by a brief pulse of NMDA or DHPG, respectively (Figure 1A). LTD induction was tested by surface labeling for GluA2 (GRIA2) one hour after the pulse, confirming that both treatments significantly reduced the surface expression of this AMPAR subunit in dendrites (Figure S1A). Control and LTD neurons were immunostained with an antibody against LC3, a protein that appears punctate when associated with autophagic vesicles and phagophores, as well as with MAP2 to label dendrites. Both forms of LTD triggered a rapid and significant increase of LC3-positive puncta in MAP2-positive dendrites only fifteen minutes after the end of the pulse (Figure 1B). Notably, pre-treatment with ifenprodil, a selective inhibitor of NR2B receptors known to have an important role in LTD induction (Kutsuwada et al., 1996) or with the mGluR inhibitors MTEP and JNJ (Linden, 2012; O’Riordan et al., 2018), prevented the appearance of LC3-positive puncta in dendrites after NMDAR- and mGluR-LTD, respectively (Figure 1C). This demonstrates that the immediate increase in dendritic autophagic structures is specifically induced by synaptic activation of NMDA and mGluR receptors.

**Figure 1.**
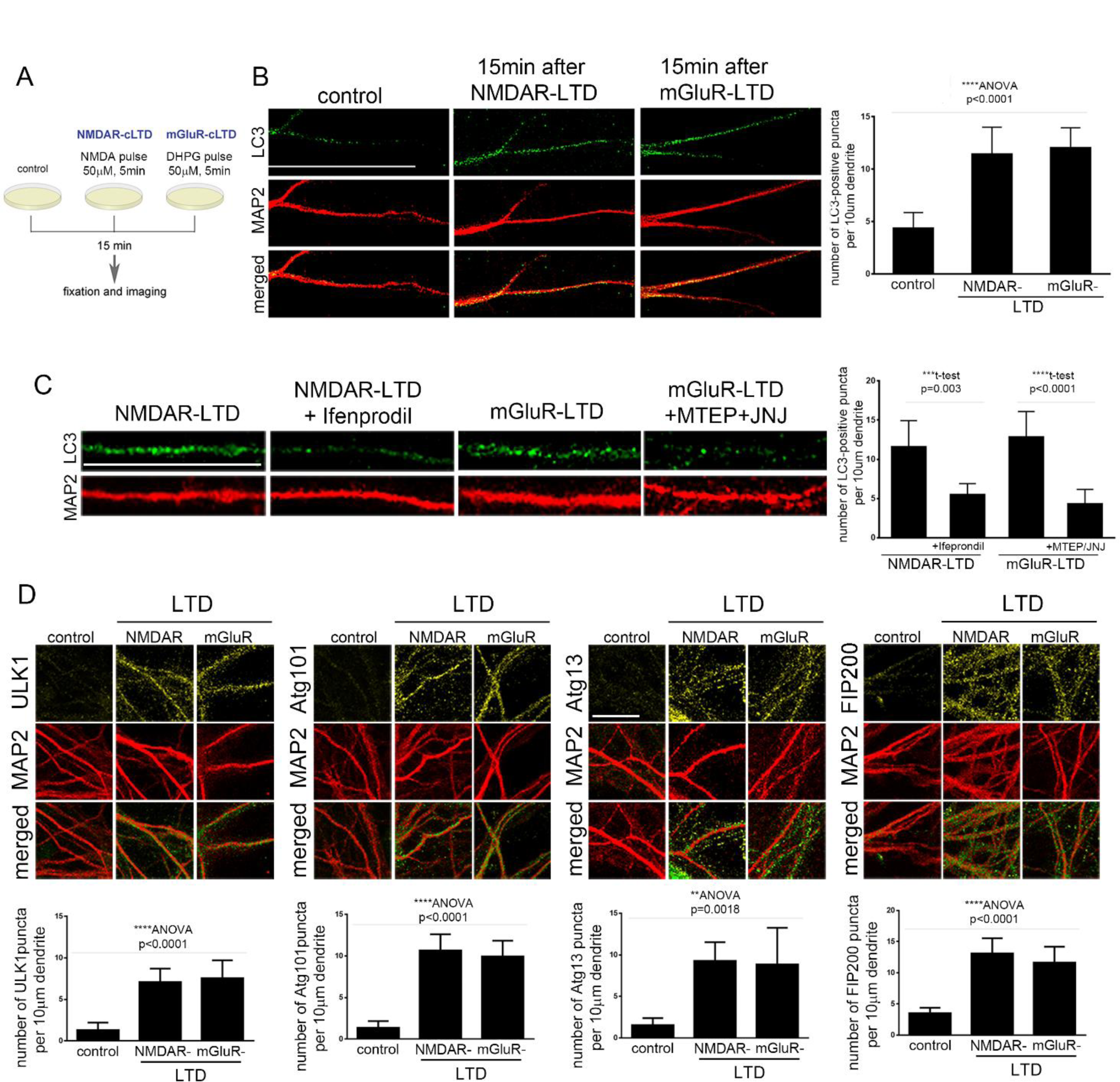
NMDAR- and mGluR-LTD trigger the rapid appearance of autophagic structures in dendrites of cultured neurons. (A) Schematic representation of chemical LTD protocols to induce NMDAR- and mGLUR-LTD in cultured neurons. (B) Representative confocal images of cultured neurons under control conditions or fifteen minutes after chemical NMDAR- or mGluR-LTD, immunostained with an antibody against LC3 to label autophagic structures and MAP2 to label dendrites. (Scale bar: 50μm). Graph showing the number of dendritic LC3-positive puncta, normalized to the dendritic length, in the indicated conditions. (N=3 independent experiments per condition). (C) Same as in (B), but neurons were either control, fifteen minutes after NMDAR-LTD and mGluR-LTD or with pre-treatment for one hour before the NMDA or DHPG pulse with Ifeprondil or MTEP and JNJ to pharmacologically inhibit NR2B and mGluR1/5 receptors, respectively. (Scale bar: 25μm). Graph showing the number of dendritic LC3-positive puncta, normalized to the dendritic length, in the indicated conditions. (N=3 independent experiments per condition). (D) Representative confocal images of neurons immuno-stained with antibodies against Atg13, ULK1, FIP200 and Atg101, along with MAP2 to label dendrites. Immuno-staining was performed in control neurons or fifteen minutes after chemical induction of NMDAR- or mGluR-LTD. Graphs show the number of puncta positive for each ULK1-complex component, normalized for MAP2, in every condition as indicated. (Scale bar: 20μm). Bars represent mean values +/- SEM. Statistical analyses were performed using student’s t-test or ANOVA, as indicated.

To determine whether the surge of dendritic AVs upon LTD represents an increase in autophagic flux, neurons were treated with Bafilomycin A1 during and for one hour after the LTD-inducing pulses. Western blot analysis for LC3 indicated increased autophagic flux at one hour after the pulses, as determined by the difference in the accumulation of lipidated LC3 (LC3-II) (Figure S1B). This was further supported by a small but significant decrease in the protein levels of the autophagic substrate p62 one hundred and eighty minutes, but not fifteen minutes, after the LTD-inducing pulses (Figure S1C).

To test whether autophagic structures are locally produced in dendrites upon LTD, we analyzed markers associated with AV biogenesis rapidly after LTD. Biogenesis is a multistep process that involves the orchestrated action of several proteins (Mercer et al., 2018). Unlike LC3, which is recruited during phagophore formation and remains associated with the autophagic membrane throughout all stages of maturation and fusion, other proteins are involved only in the initial steps of biogenesis and are then no longer found in mature AVs. One of the most upstream positions in the biogenesis cascade is occupied by the ULK1 complex, which in mammalian cells is comprised of ULK1, Atg13, FIP200 and Atg101 (Mercer et al., 2018). In yeast, the ULK1 complex is assembled upon starvation, while in cultured mouse embryonic fibroblasts it was shown to be pre-formed (Hosokawa et al., 2009). We compared the distribution of the ULK1 complex components in dendrites of cultured neurons under control conditions and fifteen minutes after chemical LTD, by immunostaining with antibodies against each of the four components and for MAP2 to label dendrites. Under control conditions, all four components were present in dendrites at very low levels and with diffuse appearance. However, fifteen minutes after NMDAR- or mGluR-LTD, the levels of all four components increased sharply in dendrites and exhibited a punctate pattern (Figure 1D).

Notably, pre-treatment with the NR2B inhibitor ifenprodil and the mGluR inhibitors MTEP and JNJ for one hour before and during the pulse prevented the appearance of Atg13-positive puncta (Figure S1D), as well as of ULK1, Atg101 and FIP200 (data not shown) in dendrites after NMDAR- and mGluR-LTD, respectively. Therefore, similar to LC3, the immediate increase in ULK1 complex component-positive structures is specifically induced by synaptic activation of NMDA and mGluR receptors.

### LTD triggers dendritic phagophores and autophagic vesicles in the hippocampus

The interplay between LTD and dendritic autophagic vesicles was next tested in hippocampal slices. NMDAR- or mGluR-LTD was induced in wild-type hippocampal slices, prepared at postnatal day 22, by a brief pulse of NMDA or DHPG, respectively. Slices were fixed fifteen minutes after the pulse and were immunolabeled with antibodies against each ULK1 complex component and MAP2 to mark dendrites. As a positive control, slices were immunostained with an antibody against Arc, an early response protein whose levels are known to be rapidly increased by local translation in dendrites in response to LTD (Figure 2A). Focusing on the *stratum radiatum* of the CA1 area (Figure 2A), we found that similar to Arc, the levels of all four ULK1 complex components sharply increased in these dendrites rapidly after NMDAR- and mGluR-LTD (Figure 2B-E). Moreover, as in cultured neurons, all four exhibited a very punctate pattern. Quantification of the number of puncta of each ULK1-complex component in CA1 *stratum radiatum* dendrites revealed that they are all significantly increased fifteen minutes after NMDAR- or mGluR-LTD, compared to control conditions (Figure 2B-E).

**Figure 2.**
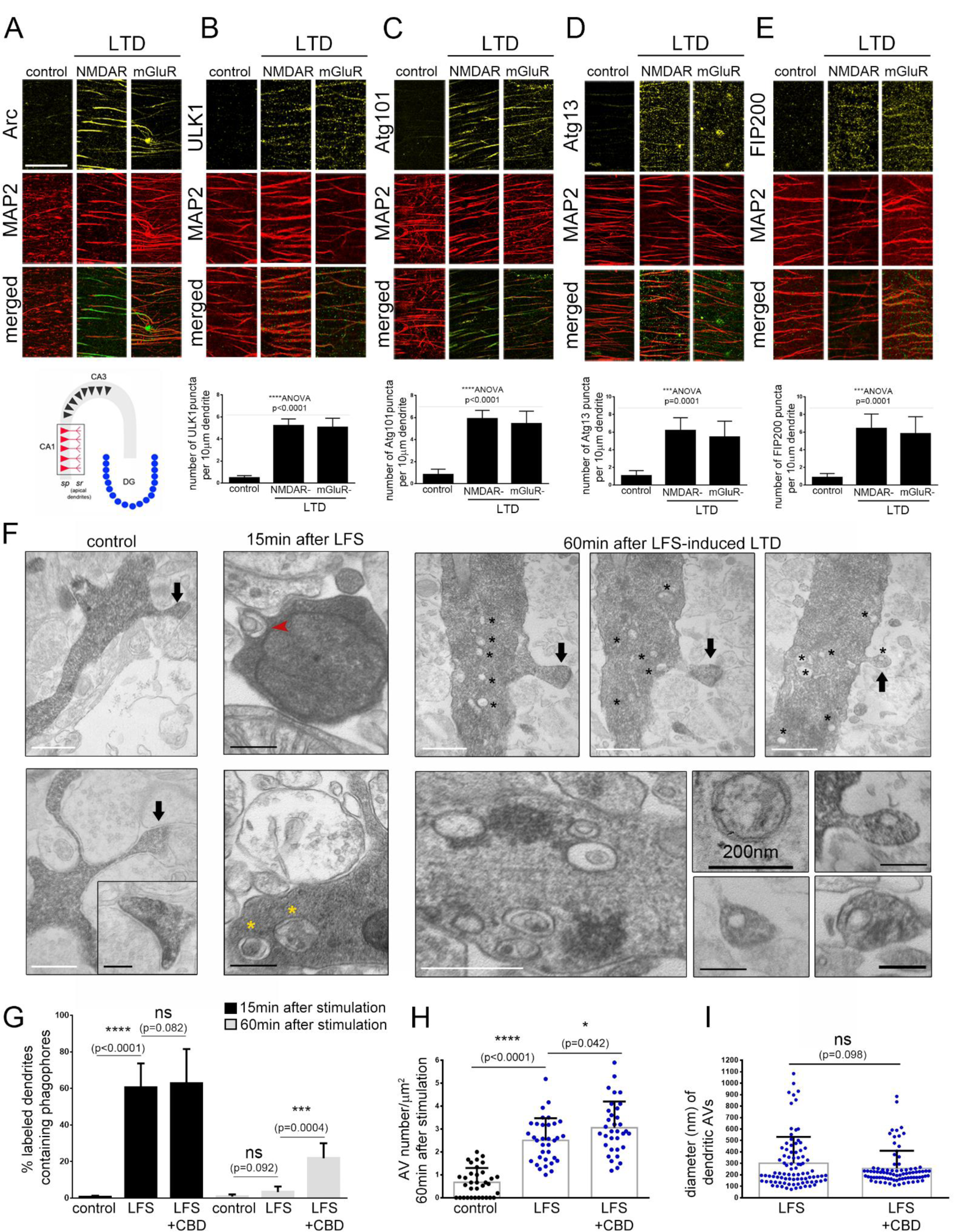
LTD triggers dendritic autophagy in hippocampal slices. (A-E) Representative confocal images of the CA1 *stratum radiatum* of wild-type P22-P28 hippocampal slices (as shown in the schematic), immuno-stained with antibodies against (A) Arc, (B) ULK1, (C) Atg101, (D) Atg13 and (E) FIP200 along with MAP2 to label dendrites. Immuno-staining was performed in control slices or fifteen minutes after chemical induction of NMDAR- or mGluR-LTD. Graphs show the number of puncta positive for each ULK1-complex component, normalized for MAP2, in every condition as indicated. (N=3 independent experiments per condition). Bars represent mean values +/- SEM. Statistical analyses were performed using ANOVA. (Scale bar: 50 μm). (F) Representative electron micrographs of neurobiotin-DAB labeled dendrites of pyramidal neurons in the CA1 *statum radiatum* in hippocampal slices under control conditions and fifteen minutes or one hour after LTD was induced by LFS. Black arrows indicate the labeled dendritic spines. Note the presence of multiple autophagic vesicles in the dendritic shaft near the neck of the spines. Bottom panels show characteristic double membrane-bound autophagic vesicles at higher magnification and the presence of vesicle-like structures inside dendritic spines. (Scale bars: 500nm (white bars), 200nm (black bars)). (G) Graph showing the percentage of labelled dendrites containing phagophores in the indicated conditions (N=48 dendrites per condition). Statistical analyses were performed using t-test. (H) Graph showing quantification of the density of autophagic vesicles in post-synaptic labeled dendrites under control conditions and one hour after LFS-induced LTD with or without application of Ciliobrevin D. (N=45 dendrites in control and 46 dendrites in LTD conditions). Statistical analyses were performed using Man Whitney U test. (I) Graph showing the size distribution of autophagic vesicles in post-synaptic labeled dendrites one hour after LFS-induced LTD with or without application of Ciliobrevin D. (N=89 autophagic vesicles). Statistical analyses were performed using Man Whitney U test.

In addition, LTD was induced by low frequency stimulation (LFS) in CA3 Shaffer collaterals in wild-type hippocampal slices and neurobiotin was concomitantly delivered by the recording electrode, allowing us to label post-synaptic dendrites in the recording area of the CA1 *stratum radiatum* (Figure S2). As control, neurobiotin was delivered under the same conditions but in the absence of LFS protocol. Neurobiotin labeling, which was achieved by 3,3′-Diaminobenzidine (DAB) method, was specific and restricted to the recording area of the CA1 (Figure S2A). Control or LTD slices were fixed fifteen minutes or one hour later (after LFS in the case of LTD) and processed for electron microscopy, as previously described (Kogo et al., 2004). AVs were visualized in labeled distal post-synaptic dendrites of pyramidal neurons in the *stratum radiatum*, as double membrane vesicles and phagophores as cup-shaped structures, and were traced in consecutive sections (Figure 2F). Consistent with previous reports (Maday and Holzbaur, 2016; Maday et al., 2012), labeled distal dendrites in control slices contained almost no phagophores and very few AVs, most often none or one vesicle per square µm surface (Figure S2B). No difference in the density of dendritic AVs was found between labeled and adjacent unlabeled dendrites in control slices (Figure S2B), indicating that labeling with neurobiotin does not by itself alter the AV content of dendrites. Moreover, neurobiotin labeling did not interfere with the ability of slices to undergo LTD upon LFS stimulation (Figure S2C).

First, we investigated the emergence of phagophores in labeled postsynaptic dendrites by electron microscopy. In line with our in vitro findings, fifteen minutes after LTD, we observed a significant increase in the percentage of labelled dendrites that contained cup-shaped phagophore structures (approximately 60%), which appeared U-shaped (indicated by red arrow) or more advanced towards closure (indicated by yellow asterisks) (Figure 2F, G). By contrast, one hour after stimulation this increase was not observed, and both LTD and control dendrites rarely contained phagophore structures. Bath application of Ciliobrevin D, a specific inhibitor of dynein-mediated transport, two hours prior, during and after the LFS stimulation did not prevent the emergence of phagophores in dendrites fifteen minutes after stimulation (Figure 2F), suggesting that phagophores are locally produced. However, Ciliobrevin treatment resulted in significantly more dendrites containing phagophores one hour after LTD stimulation (Figure 2G), suggesting that dynein may mediate their transport towards the soma for fusion with lysosomes.

Next, we investigated the emergence of complete, double-membrane AVs by electron microscopy. While AVs were not detected fifteen minutes after stimulation, a significant increase in their density, of approximately 2.5-fold compared to control, was observed in the dendritic shaft one hour after LTD, often within 1µm from the base of dendritic spines (Figure 2F, H). Bath application of Ciliobrevin D two hours prior, during and after the LFS stimulation did not prevent the increase in dendritic AV density one hour after LFS. Instead, as with phagophores at fifteen minutes after LFS, ciliobrevin D further potentiated the increase in dendritic AV density one hour after LFS, suggesting that AVs may be trafficked away from dendrites and towards the soma via dynein.

Most double membrane, complete autophagic vesicles were 200-300 nm in diameter, and only few vesicles had larger diameters ranging from 400-700 nm (Figure 2F, I). Often, a small vesicle which appeared incompletely sealed and approximately 100nm in diameter was found in spine heads, as shown in Figure 2F, with a morphology clearly distinct from that of the spine apparatus. Ciliobrevin D had no effect on the size of dendritic AVs (Figure 2I).

Taken together, these findings indicate that LTD triggers the biogenesis of dendritic autophagic vesicles both in cultured neurons and in acute hippocampal slices.

### Local translation of autophagy initiation components in dendrites upon LTD

The rapid appearance of phagophores shortly after LTD induction prompted us to investigate whether proteins involved in AV biogenesis may be locally translated on site. First, we performed bio-orthogonal labeling of newly synthesized proteins in hippocampal slices after LTD induction. NMDAR- or mGluR-LTD was induced in hippocampal slices by a brief pulse of NMDA or DHPG respectively and were exposed for twenty minutes to L-azidohomoalanine (AHA), a compound that is incorporated in newly synthesized proteins in the place of methionine (Tom Dieck et al., 2012). Slices were then fixed and subjected to a click-reaction with biotin-alkyne, thus biotinylating all AHA-containing newly synthesized proteins. NMDAR-LTD slices in the presence or absence of a click-reaction were immunostained with antibodies against biotin and Arc. Biotin staining was specific to click-reaction exposed slices and completely abolished in the absence of click reaction (Figure 3A). Moreover, biotin staining co-localized to a large extent with the signal of Arc in dendrites of the *stratum radiatum* in the CA1 area (Figure 3A). Immunostaining of NMDAR- and mGluR-LTD slices with antibodies against each ULK1-complex component and biotin demonstrated that Atg13, ULK1, FIP200 and Atg101 all co-localize with biotin in dendrites (Figure 3B). To determine whether these proteins are themselves biotinylated, and therefore locally translated, we performed immunoprecipitation of each ULK1 complex component in hippocampal slice lysates of control and NMDAR-LTD slices that were subjected to AHA-labeling and click reaction as previously described, followed by blotting with an antibody against biotin. In all cases, there was a marked increase in the biotin-positive signal at the expected size of each protein in the LTD condition compared to control, indicating that these components are translated rapidly after LTD induction (Figure 3C).

**Figure 3.**
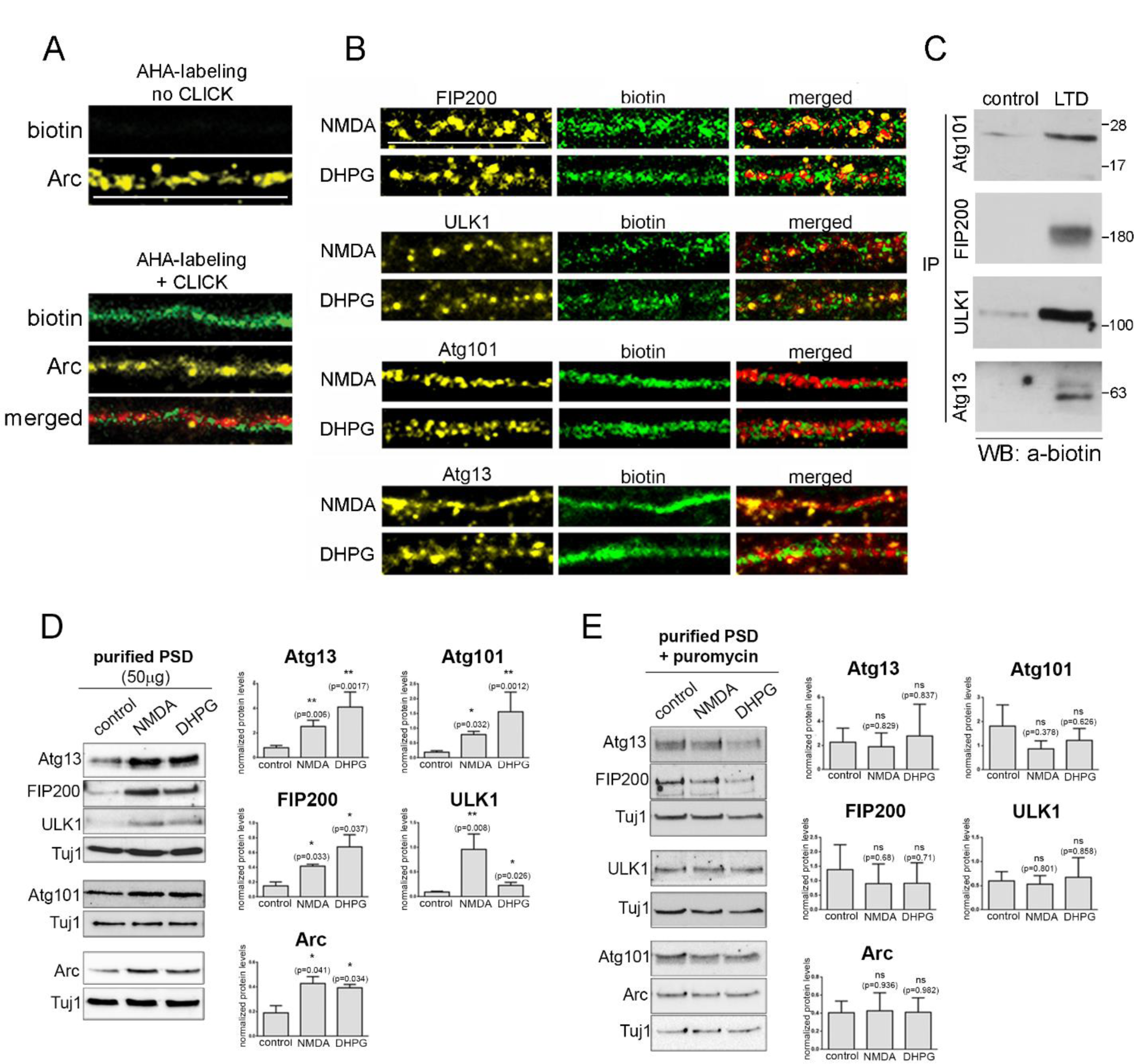
ULK1 complex components are locally translated in dendrites upon LTD. (A) Confocal images of CA1 *stratum radiatum* dendrites immunolabeled with antibodies against Arc and biotin. Immunolabeling was performed in NMDAR-LTD slices exposed to AHA and subjected or not to a click-reaction to biotinylate newly synthesized proteins. Note the specificity of the biotin immunoreactivity only in the click-reaction sample. (Scale bar: 20μm) (B) Representative confocal images of CA1 *stratum radiatum* dendrites immuno-stained with antibodies against each ULK1-complex component, biotin and MAP2. Immuno-staining was performed in control, NMDAR and mGluR-LTD slices, exposed to AHA and click reaction to biotinylate newly synthesized proteins. Note the co-localization of all ULK1-complex components with biotin. (Scale bar: 20μm) (C) Immunoprecipitation with antibodies against the indicated proteins, followed by western blot analysis with an antibody against biotin. The experiment was performed in lysates of hippocampal slices which were AHA-treated and subjected to a click-reaction under control or NMDAR-LTD conditions. (D) Western blot analyses for Atg13, FIP200, ULK1, Atg101, Arc and β-III tubulin (Tuj1) in synaptosomes, under control conditions and ten minutes after exposure to a ten-minute pulse of 50 μM NMDA or 50μM DHPG. Graphs showing the normalized protein levels of Atg13, FIP200, ULK1, Atg101and Arc in synaptosomes under control conditions and ten minutes after the NMDA or DHPG pulse. Bars represent mean values +/- SEM (N=3 independent experiments). (E) Western blot analyses for Atg13, FIP200, ULK1, Atg101, Arc and normalized for β-III tubulin (Tuj1) in synaptosomes ten minutes after exposure to a pulse of 50μM NMDA or DHPG in the presence of 1 μg/ml puromycin. Graphs showing the normalized protein levels of Atg13, FIP200, ULK1, Atg101 and Arc in the aforementioned conditions. Bars represent mean values +/- SEM (N=3 independent experiments).

We also validated these conclusions using a biochemical approach. We took advantage of the fact that biochemically purified synapses (synaptosomes), are devoid of other neuronal compartments but retain many functional properties of synapses, including their ability to respond to LTD-triggering stimuli (Bailey et al., 2003). Therefore, we treated purified PSDs with a low dose ten-minute NMDA- or DHPG-pulse and assessed the levels of ULK1-complex components fifteen minutes later by western blot analysis. Protein levels of Atg13, Atg101, ULK1 and FIP200 were significantly increased shortly after the NMDA and DHPG pulse, as did the positive control protein Arc (Figure 3D). Notably, treatment of purified synapses with puromycin, a translation inhibitor, during and fifteen minutes after the NMDA- and DHPG-pulse, completely prevented the upregulation of ULK1-complex components (Figure 3E), further indicating that these components are locally translated in the synapse.

### Inhibition of autophagic vesicle biogenesis abolishes NMDAR- and mGluR-LTD

Next, we sought to investigate whether LTD requires the formation of new autophagic vesicles. To this end, we tested SBI-0206965, a selective inhibitor of ULK1-mediated phosphorylation, which prevents the initial steps of autophagic vesicle biogenesis (Egan et al., 2015). Bath application of hippocampal slices in 500nM SBI-0206965 for fifteen or sixty minutes resulted in decreased phosphorylation of Atg13 at serine-318, a known target of ULK1 during autophagy initiation (Joo et al., 2011; Puente et al., 2016) (Figure 4A). Using acute hippocampal slices from wildtype animals, we tested whether inhibition of biogenesis impairs LTD. Bath application of SBI-0206965 for ten minutes before, during and ten minutes after LFS, totally prevented LFS-induced LTD (Figure 4B) and triggered instead potentiation.

**Figure 4.**
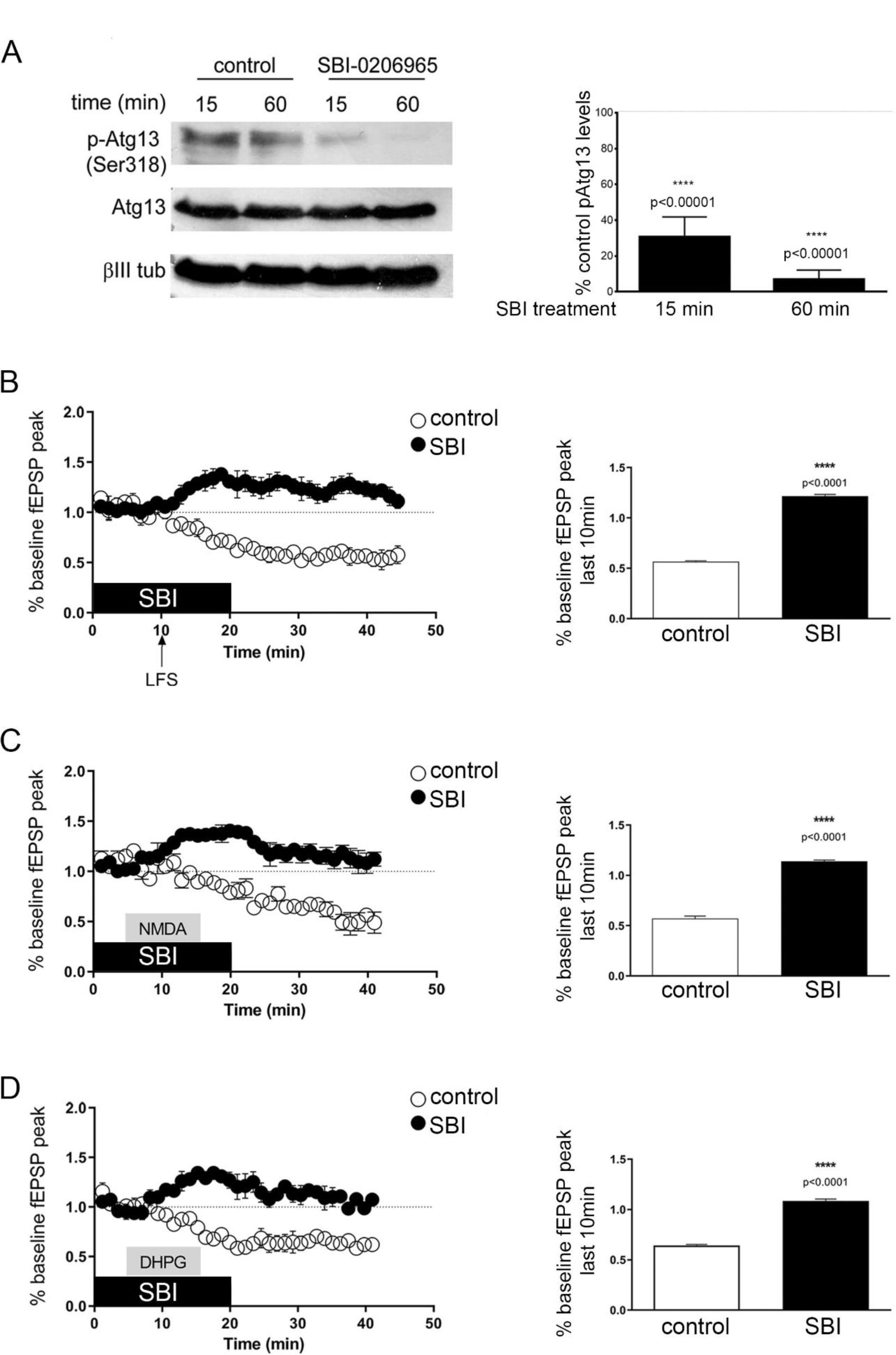
Pharmacological inhibition of autophagic vesicle biogenesis impairs LTD. (A) Quantification of Western blot analysis in lysates of wild-type P22 hippocampal slices treated by bath application with control vehicle or with 500nM SBI-0206965 for fifteen and sixty minutes. Blots show levels of pSer318-Atg13 and total Atg13 and normalized for β-III tubulin. (B-D) Time-plots of the normalized fEPSPs and bar graphs of average normalized fEPSP peaks during the last ten minutes of recording after (B) a fifteen -minute LFS stimulation, (C) a ten-minute NMDA (20µM) pulse or (D) a ten-minute DHPG (20µM) pulse in wild-type P22 hippocampal slices under control conditions (white circles and bars), or bath application with 500 nM SBI-0206965 (SBI) for the indicated duration (black circles and bars). N=6 animals per condition. In all cases, recordings are performed on 3 slices per animal. (Bars represent mean values +/- SEM. Statistical analyses was performed using student’s t-test).

NMDAR-LTD was induced in acute hippocampal slices by a pulse of NMDA, but was completely abolished by bath application of SBI-0206965 five minutes before, during and five minutes after the NMDA pulse (Figure 4C). Similarly, mGluR-LTD was induced by a DHPG pulse in control conditions, but was abolished by bath application of SBI-0206965 five minutes before, during and five minutes after the pulse (Figure 4D). Taken together, these findings indicate that both major forms of LTD require autophagic vesicle biogenesis during the early phase of LTD induction.

### Autophagy is cell autonomously required in excitatory neurons for NMDAR- and mGluR-LTD

To test whether autophagy is cell autonomously required in excitatory neurons for LTD, the *atg5* gene was conditionally ablated in excitatory neurons by crossing *atg5^f/f^* with *thy1-Cre^ERT2^* animals. Tamoxifen was administered to *thy1-Cre^ERT2^;atg5^f/f^* progeny for five consecutive days starting at postnatal day fifteen (P15). Atg5 protein levels were examined by Western blot in forebrain lysates of *atg5^f/f^* controls, *nestin-cKO*s, which lack atg5 in the entire neural lineage, and *thy1-cKO*s. As expected, atg5 was undetectable in *nestin-cKO*s, whereas in *thy1-cKO*s it was detectable but dramatically reduced compared to controls (Figure 5A).

**Figure 5.**
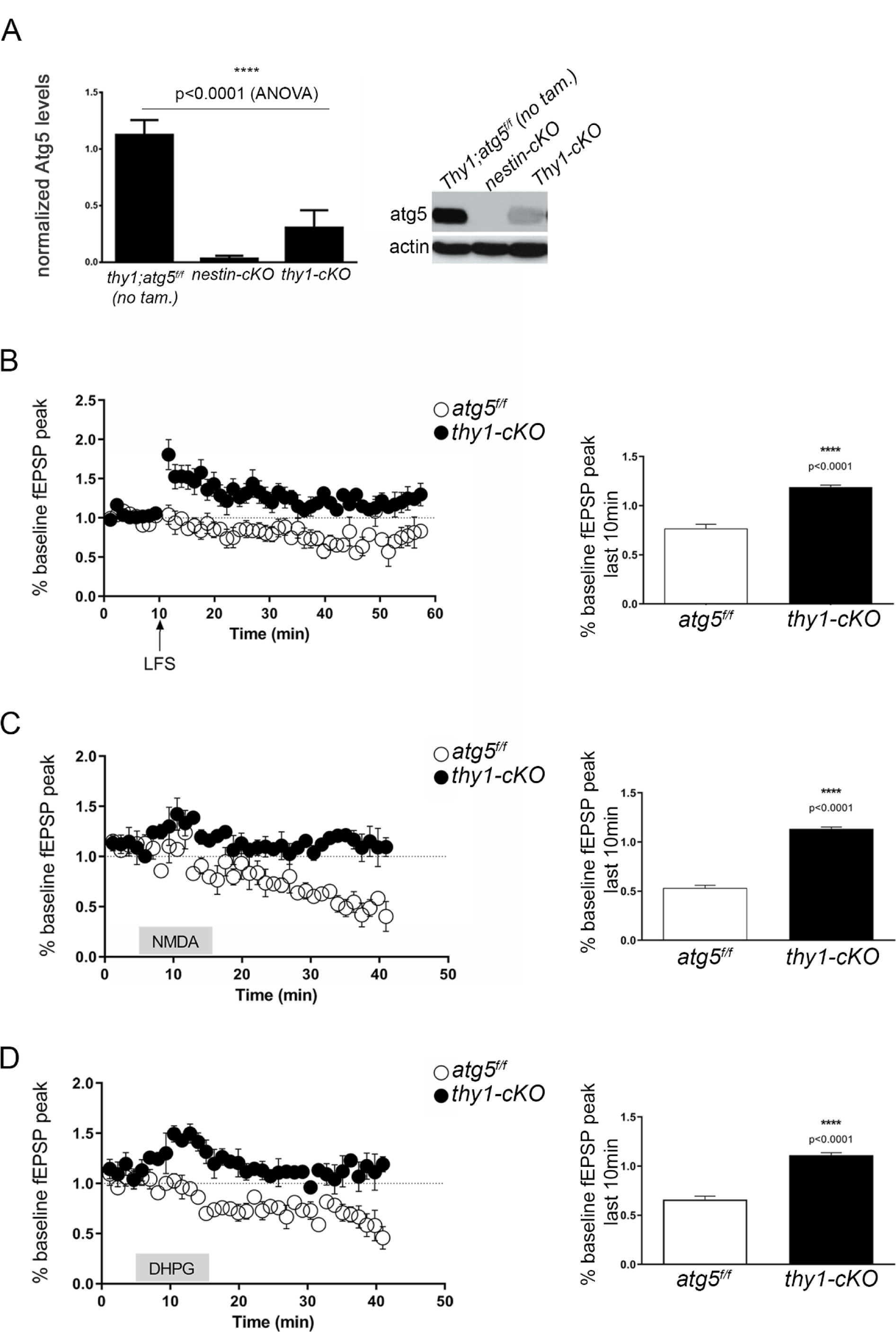
Autophagy is required cell autonomously in pyramidal neurons for LFS-, NMDAR- and DHPG-induced LTD. (A) Western blot analysis in P22 forebrain lysates of the indicated genotypes with an antibody against Atg5 and normalized against actin. Graph shows quantification of normalized Atg5 protein levels in corresponding genotypes. (N=3 animals per genotype). (B-D) Time-plots of the normalized fEPSPs and bar graphs of average normalized fEPSP peaks during the last ten minutes of recording after a fifteen-minute LFS stimulation (B) or a ten-minute NMDA (20µM) pulse (C) or ten-minute DHPG (20µM) pulse (D) in P22 hippocampal slices from control (*atg5^f/f^*, black circles and bars) and *thy1-cKO* (blue circles and bars) animals. (N=6 *atg5^f/f^* control and 5 *thy1-cKO* animals for LFS-induced LTD. N=6 *atg5^f/f^* control and 6 *thy1-cKO* animals for NMDA and DHPG-induced LTD. In all cases, recordings are performed on 3 slices per animal). (Bars represent mean values +/- SEM. Statistical analyses was performed using student’s t-test).

A cross-talk between autophagy and the ubiquitin-proteasome system (UPS) has been described in several cell types (Park and Cuervo, 2013), including cultured neurons (Du et al., 2009). Therefore, we first tested whether *thy1-cKO* animals exhibit altered proteasomal activity compared to control littermates. Comparing proteasome activity in cortical (Figure S3A) and hippocampal (Figure S3B) lysates of *thy1-cKO* and control animals at P40 indicated that there are no significant differences among these genotypes.

To investigate whether autophagy is cell-autonomously required in pyramidal neurons for LTD, we performed electrophysiology recordings from acute hippocampal slices prepared from P22 *thy1-cKO* and *atg5^f/f^* control animals. This age was purposefully chosen, as *atg5* is only ablated starting with tamoxifen at P15, thus excluding the accumulation of any defects from prolonged autophagy ablation.

We first compared the ability of low-frequency stimulation (LFS) (1 Hz for fifteen minutes) in Shaffer collaterals to elicit an LTD response in the CA1 *stratum radiatum*. LTD was successfully induced in control slices, as shown by the depression of the field excitatory postsynaptic potential (fEPSP), but completely failed in *thy1-cKO* slices (Figure 5B). Similarly, LTD was successfully induced in control slices by an NMDA (Figure 5C) or a DHPG pulse (Figure 5D), but completely failed in *thy1-cKO* slices (Figure 5C-D). Taken together, these results indicate that both major types of LTD depend on the autophagic machinery in pyramidal neurons.

### Proteomic profiling of the autophagic cargo upon LTD

Although autophagy is a major cellular mechanism for protein degradation, the identity and dynamic regulation of the autophagic cargo in neurons remains largely unknown. Pertinent to LTD, AMPAR subunits were shown to be substrates of autophagy (Shehata et al., 2012). However, in addition to the internalization and degradation of AMPARs, LTD entails the shrinkage and elimination of entire spines. Therefore, we postulated that autophagy might facilitate the degradation of many post-synaptic components. In order to understand how autophagy is indispensable for LTD, we sought to unravel the identity of the cargo sequestered in autophagic vesicles during this process.

To this end, hippocampal slices of P22-28 mice were exposed either to control conditions or to NMDAR-LTD and AVs were purified one hour later using a method that we recently established (Nikoletopoulou et al., 2017). Observation of the preparation by electron microscopy confirmed its high degree of purity, consisting of mature, closed AVs (Figure 6A). Western blot analysis of the purified AVs, and of other fractions obtained along the purification procedure, further indicated that the final AV preparation is enriched in LC3-II but negative for the phagophore marker Atg16L1, and for the ER marker GRP78Bip (Figure 6B).

**Figure 6.**
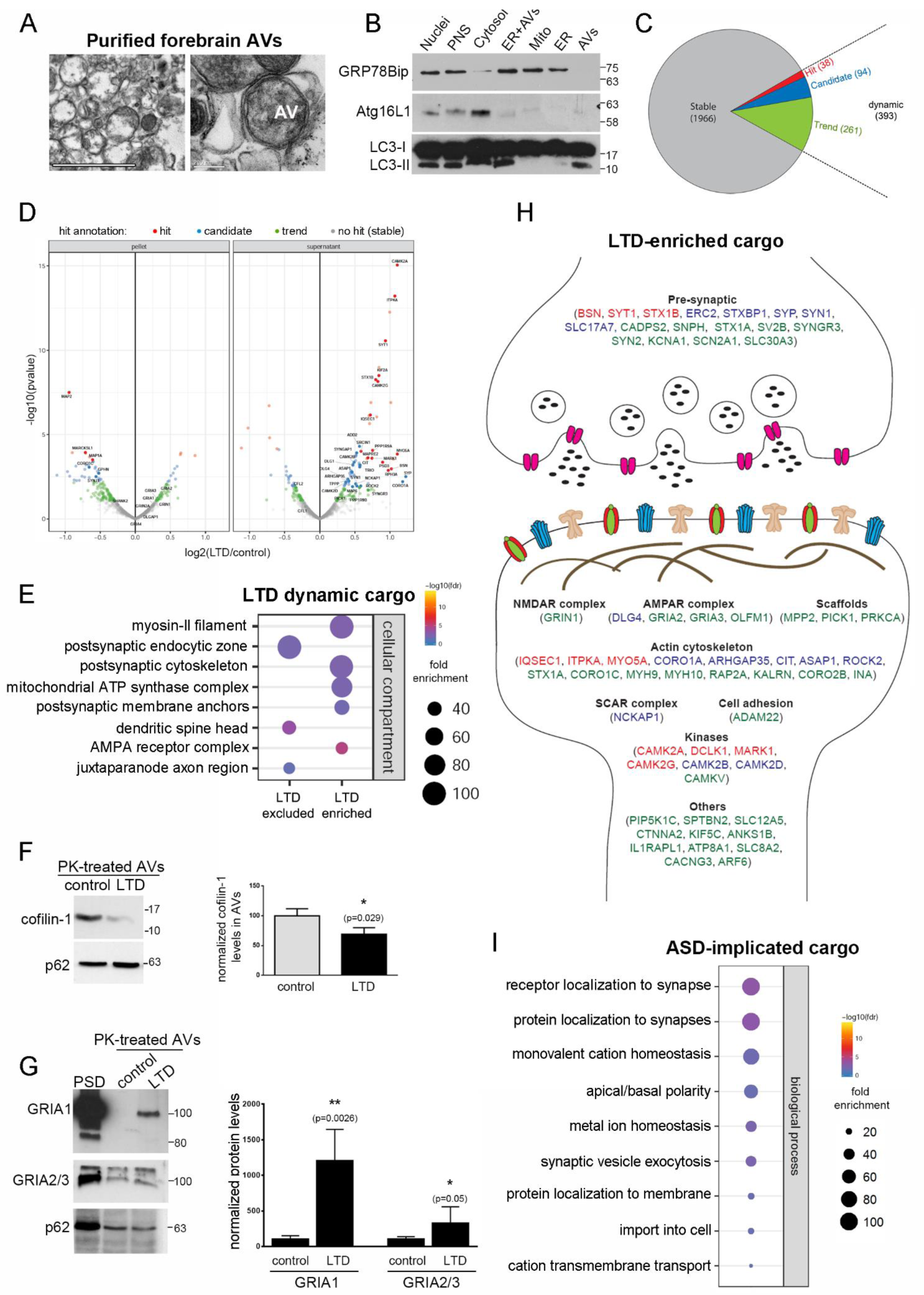
Proteomic profiling of the autophagic cargo during LTD. (A) Representative electron micrographs of the purified AV preparation, showing that it is comprised of intact, double membrane-bound vesicles. (Scale bar: 200 nm (right), 2 µm (left)). (B) Western blot analysis of different fractions along the purification procedure, with antibodies against the ER marker GRP78Bip, the phagophore marker Atg16L1 and LC3, which is present in all autophagic structures. Note that the final AV preparation is devoid of ER or phagophore markers, but positive for LC3-II. (C) Pie chart showing the proportion of the autophagic cargo that is stable between baseline and NMDAR-LTD conditions (grey), or dynamic in LTD (red, blue and green). Red represents hits with an value of as false discovery rate (FDR)-corrected p-values <0.05, blue represents candidates with a value of fdr<0.2 and green represents trends with a value of fdr<0.6. Numbers indicate the actual number of proteins in each category. (D) Volcano plot analysis of the proteins found in pellet and supernatant fractions of purified AVs. Color code is as in (C). (E) Graph showing the cell component analysis, as false discovery rate (FDR)-corrected p-values, of the dynamic cargo (total of 393 proteins) that is enriched (up) or less abundant (down) in AVs after LTD, compared to control. (F) Western blot analysis of proteinase K-treated control and LTD AVs with an antibody against cofilin-1 and normalized against p62. Graph shows the normalized levels of cofilin-1 in these conditions. (N=4 independent AV purifications). Bars represent mean values +/-SEM. Statistical analysis was performed using Student’s t-test. (G) Western blot analysis of purified post-synaptic densities and of proteinase K-treated control and LTD AVs with an antibody against GRIA1 and GRIA2/3, and normalized against p62. Graph shows the normalized levels of glutamate receptor subunits in proteinase K (PK)-treated control and LTD AVs. (N=4 independent AV purifications). Bars represent mean values +/-SEM. Statistical analysis was performed using Student’s t-test. (H) Graphical representation of proteins enriched in AVs upon LTD, with relation to the synapse. (I) Graph showing the biological process analysis, as false discovery rate (FDR)-corrected p-values, of the ASD-implicated proteins contained in the LTD enriched cargo (total of 80 proteins).

Control- and LTD-AVs were then subjected to carbonate extraction to separate the fraction containing the autophagic membranes from the one containing soluble material, as previously described (Nikoletopoulou et al., 2017). Both soluble and membrane fractions were subjected to TMT labeling and quantitative proteomic analyses. We found a total of 2359 proteins, of which the vast majority (1966 proteins, 83.3%) were equally abundant between control and LTD conditions and therefore constitute a stable or constitutive cargo of autophagy (Figure 6C). The remaining 393 proteins, summarized in Table S1, showed a dynamic behavior between control and LTD conditions and were annotated as hits (38 proteins), candidates (94 proteins) and trends (261 proteins), according to their fdr value (<0.05, <0.2 and <0.6 respectively) (Figure 6C,D). Approximately half (48.6%) of the dynamic cargo was preferentially enriched in LTD-AVs, while the other half (51.4%) was preferentially excluded from LTD-AVs. Enrichment analyses for cellular component (Figure 6E) indicated that the “LTD-excluded” cargo is enriched for proteins that localize in the postsynaptic endocytic zone and spine head, including cofilin-1, an actin filament-severing protein, with established function in dendritic spine shrinkage and elimination during LTD (Rust et al., 2010; Zhou et al., 2004). In line with the proteomic data, western blot analysis confirmed that the levels of cofilin-1 are significantly reduced in proteinase K-treated AVs in LTD compared to baseline conditions, as normalized against the cargo protein p62, whose levels are stable between the two conditions (Figure 6F).

The dynamic “LTD-enriched” cargo was found to consist mainly of proteins localized to the postsynaptic cytoskeleton, including myosin-II filament proteins and to postsynaptic membrane anchoring comlexes. This group also contained AMPA receptor complex proteins, including in particular AMPA receptor subunits (GRIA1,2/3). In line with the proteomic data, western blot analysis confirmed that the levels of these AMPA receptors were significantly enriched in proteinase K-treated LTD-AVs, compared to control (Figure 6G). As summarized in Figure 6H, the “LTD-enriched” cargo further included prominent scaffold proteins involved in receptor localization to the postsynaptic density, kinases involved in synaptic plasticity and cell adhesion molecules, as well as a number of presynaptic proteins involved in neurotransmitter release.

Notably, we observed that 80 proteins (20%) of the dynamic cargo are implicated in autism spectrum disorders (ASD) according to the gene scoring module within SFARI Gene 3.0 (https://gene.sfari.org) (Abrahams et al., 2013) (Table S2). Of these, the majority (49 proteins) are enriched in LTD-AVs and have functions in the localization of proteins and receptors to the synapse and in regulating ion transport, as determined by biological process enrichment analysis (Figure 6I). In view of findings that propose impaired LTD as an underlying cause of several ASD mouse models (Hansel, 2019), these findings may help explain how autophagy deficiency in pyramidal neurons can cause impaired proteostasis of key ASD-implicated proteins, thus leading to cognitive deficits characteristic of autism (Tang et al., 2014).

### Mild autophagy impairment results in cognitive flexibility deficit

Synapse destabilization is required for key cognitive functions and LTD has been particularly recognized for its role in cognitive flexibility. Therefore, we hypothesized that autophagy impairment may lead to deficits in cognitive flexibility by preventing LTD induction. However, as cell type-specific and complete ablation of autophagy is unlikely to occur during physiological perturbations, we opted to analyze the effects of global but mild autophagy impairment. To this end, we analyzed mice with conditional ablation of one *atg5* allele in the neural lineage, using the *nestin-Cre* delete mice. These animals, referred to as *nestin-cHET*, exhibit reduced atg5 protein levels by approximately half compared to control (*atg5^f/+^*) animals (Figure 7A). This atg5 deficiency is sufficient to trigger a mild autophagy impairment, as indicated by the increased levels of p62, an autophagy receptor and substrate, in *nestin-cHET* animals compared to control in immunostaining of cortical sections (Figure 7B). However, unlike *nestin*-cKO animals that exhibit late-onset neurodegeneration (Hara et al., 2006; Komatsu et al., 2006), similar numbers of TUNEL-positive cells were found at the forebrain of *nestin-cHET* and *atg5^f/+^* (control) animals at P60 (Figure S4A).

**Figure 7.**
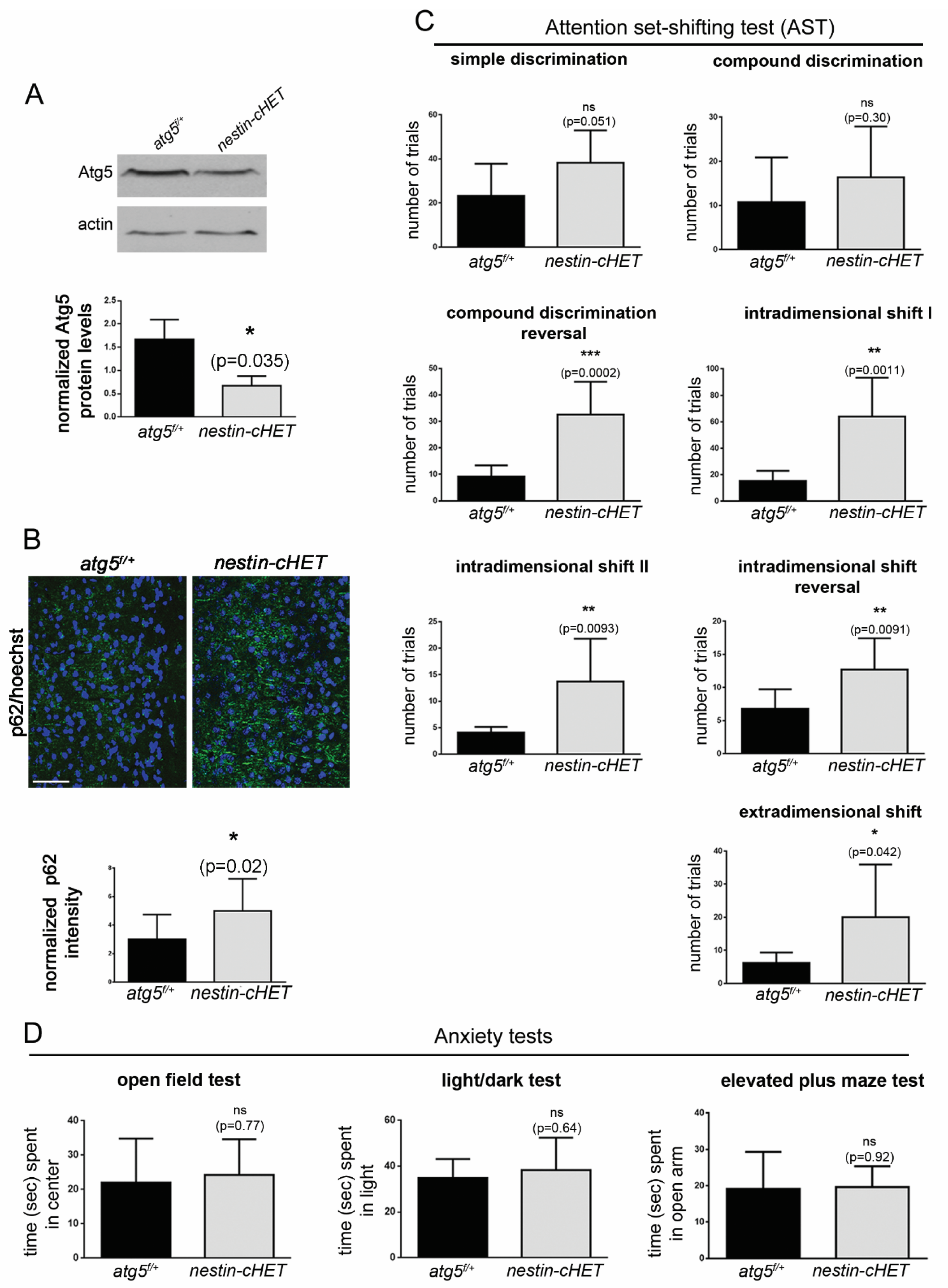
Autophagy is required for cognitive flexibility. (A) Western blot analysis in P22 forebrain lysates of the *atg5^f/+^ and nestin-cHET* animals with an antibody against Atg5 and normalized against actin. Graph shows quantification of normalized Atg5 protein levels. (N=3 animals per genotype). Bars represent mean values +/- SEM. Statistical analyses were performed by t-test. (B) Representative confocal images of forebrain cryosections of P40 *atg5^f/+^ and nestin-cHET* animals immuno-stained with an antibody against p62 and with hoechst dye to label nuclei. Bar graph shows the quantification of p62 levels normalized to the number of nuclei in the indicated genotypes. (N=3 animals per genotype and 3 sections per animal). (Scale bar: 50 μm). Bars represent mean values +/- SEM. Statistical analyses were performed by t-test. (C) Behavioral analysis of a*tg5^f/+^*and *nestin-cHET* mice in an attention set-shifting test. Graphs showing quantification of the number of trials per genotype in a test of simple discrimination, compound discrimination, compound discrimination reversal, intradimensional shift I, intradimensional shift II, intradimensional shift reversal and extradimensional shift. (D) Behavioral analysis of a*tg5^f/+^*and *nestin-cHET* mice in three anxiety tests. Graph showing the time spent in the centre of the cage in the open field test for the indicated genotypes, in the light compartment of the cage in the light/dark test for the indicated genotypes and in the open arm of the elevated plus maze for the indicated genotypes. Bars represent mean values +/- SEM. Statistical analyses were performed by t-test. (N=10 P40 animals/genotype for attention set shifting tests. N=10 P40 animals/genotype for anxiety tests.)

In order to assess cognitive flexibility, *nestin-cHET* and control (*atg5^f/+^*) animals were compared in an attentional set shifting behavioral paradigm, starting at P40. While *nestin-cHET* animals were indistinguishable from control littermates in simple discrimination and compound discrimination tasks, they exhibited a significant impairment in compound discrimination reversal, intradimentional shifts, intradimensional shift reversal and extradimentional shift tasks, collectively indicating deficits in flexible choice behavior (Figure 7C). Of note is that *nestin-cre;atg5^+/+^* animals were compared to *atg5^f/+^* mice in simple and compound discrimination tests. In both cases, no significant differences were observed between genotypes (Figure S4B).

By contrast, in an open field test, a light/dark test or an elevated plus maze test (Figure 7D), three different behavioral paradigms of anxiety, *nestin-cHET* animals were indistinguishable from control littermates. These findings demonstrate that mild and global impairment of autophagy is not associated with anxiety disorders but causes deficits in LTD-mediated behaviors, consistent with its requirement for LTD induction.

## Discussion

The role of autophagy in long-term synaptic plasticity, conducive to protein turnover, is beginning to be elucidated. Our previous work demonstrated that BDNF, a key regulator of long-term potentiation (LTP), suppresses autophagy in the adult brain. As a result, conditional BDNF mutants exhibit increased autophagic flux and an overabundance of AVs in the hippocampus. Interestingly, inhibition of autophagy fully rescues the LTP-impairment caused by sequestration of endogenous BDNF (Nikoletopoulou et al., 2017). In the present study, we demonstrate for the first time that autophagy is cell autonomously required in pyramidal neurons for the induction of both major forms of LTD that co-exist in the hippocampus, mediated by synaptic activation of either NMDA or group I metabotropic glutamate receptors. It therefore emerges that, depending on the type of plasticity, autophagy induction differentially modulates synaptic efficacy, with its upregulation being required for LTD induction but also implicated in LTP impairment (Nikoletopoulou et al., 2017).

Notably, the absolute requirement of autophagy in LTD induction is opposite to the function exerted by the ubiquitin proteasome system (UPS), the second major degradative pathway. Specifically, UPS inhibitors, such as MG132 and lactacystin, have been shown to promote LTD by facilitating the transition from early- to late-LTD, thereby revealing a modulatory role of the UPS in attenuating LTD progression (Li et al., 2016). It remains to be seen whether there is a cross regulation between autophagy and UPS during synaptic plasticity, as UPS impairment has been shown to cause an upregulation of the autophagic flux in other cellular contexts (Park and Cuervo, 2013).

We also found that both major forms of LTD are completely blocked by pharmacological inhibition of ULK1 kinase activity, which is required for the early steps of the cascade leading to the formation of autophagic vesicles. Therefore, these findings suggest that LTD induction relies on novel biogenesis of autophagic vesicles. As LTD is expressed by the shrinkage or elimination of dendritic spines, mainly enriched in distal dendrites, it is difficult to conceptualize how the previously described biogenesis of AVs at the axon tip (Maday and Holzbaur, 2016) can rapidly contribute to degradation in remote dendritic compartments. We therefore, used biochemical tools and electron microscopy to examine an alternative hypothesis, and revealed the local and rapid biogenesis of autophagic vesicles in postsynaptic dendrites following LTD. This regulation of the biogenesis cascade by synaptic activity allows for post-synaptic material to be rapidly accessible on-site to phagophores in order to facilitate their sequestration into nascent vesicles.

A key question relates to the nature of the synaptic and plasticity-related proteins that are degraded during LTD. Previous studies used candidate-based approaches to suggest that autophagy can degrade AMPA (Shehata et al., 2012) and GABAB receptor subunits (Rowland et al., 2006). In order to gain inclusive insight into autophagic cargo dynamics upon LTD, we performed unbiased quantitative proteomic analyses comparing intact AVs under baseline conditions or 1 hour after LTD induction. An important first conclusion is that the autophagic cargo is largely stable, as the vast majority of cargo proteins show comparable abundance between these conditions and only a small fraction of the cargo is dynamically regulated during LTD.

The dynamic cargo mainly consists of proteins that are relatively enriched or excluded in LTD compared to control baseline AVs. Interestingly, enriched proteins comprise key players of the post-synapse, including not only AMPA receptor subunits, whose removal from the synapse during NMDAR-LTD is well documented (Carroll et al., 1999a; Carroll et al., 1999b; Collingridge et al., 2004; Heynen et al., 2000; Luscher et al., 1999; Man et al., 2000), but also scaffold proteins, cell adhesion molecules and numerous modulators of the actin cytoskeleton. This group also includes a number of presynaptic proteins involved in neurotransmitter release, in line with evidence that LTD can involve a reduction in the probability of glutamate release (Enoki et al., 2009; Stanton et al., 2003). Part of the dynamic cargo consists of proteins that are less abundant upon LTD, and therefore selectively escape degradation during this process. One such example is cofilin-1, an actin filament severing protein with established role in structural plasticity (Noguchi et al., 2016), These findings establish autophagy as the major degradative route not only for specific receptors but more generally for synaptic components, whose degradation can collectively facilitate the shrinkage or elimination of spines. Moreover, they suggest that synaptic plasticity also regulates mechanisms that control synaptic cargo selectivity, emphasizing the need for their molecular characterization.

Our results further reveal that autophagic degradation is responsible for the turnover of many ASD-susceptibility proteins (Abrahams et al., 2013) during synaptic plasticity, in line with the fact that autophagy-deficient mouse models exhibit autistic-like behaviors (Tang et al., 2014) and that small exonic copy number variations from whole-exome sequence data of patients implicate dysregulation of autophagy in ASD (Poultney et al., 2013). Consistently, we found that reducing autophagy even by half in the brain is sufficient to impair LTD and behaviors such as cognitive flexibility that depend on functional LTD and are impaired in autism.

Therefore, in addition to elucidating the role of autophagy in long-term synaptic depression our study provides a molecular basis for developing novel therapeutic approaches to restore synaptic and behavioral deficits in disorders that entail impaired synaptic plasticity.

## Acknowledgements

The authors thank all lab members for their input. This work was supported by an ERC starting grant with the acronym “NEUROPHAGY” to V.N. and a PhD fellowship from the Hellenic Foundation for Research and Innovation (ELIDEK) to A.D.D.

## Author contributions

V.N. designed the study. K.S and A.K. performed electrophysiology experiments. Y.D. performed electron microscopy experiments. M.M.S., P.H. and F.S. performed the proteomic analyses and F.S performed the biostatistical analyses of the proteomics. M.P. helped with behavioral analyses. E.K., E.I. and A.D.D. carried out all other experiments and analyzed data. V.N., E.K., and A.D.D. wrote the manuscript.

## Methods

### Animals

The animal protocols of this study were approved by the FORTH Animal Ethics Committee (FEC). The mice were maintained in a pathogen-free environment and housed in groups of five animals per cage with constant temperature and humidity and a 12 hour/12 hour light/dark cycle. The animals used were of C57BL/6 genetic background. *Atg5^f/f^* ^(^Hara et al., 200^6^)*, nestin-Cre* mice (Nikoletopoulou et al., 2017) and *SLICK-H-Thy1-cre/^ERT2^-EYFP* (Jax labs strain 012708) mice were used for electrophysiology experiments and behavioral analyses. In *thy1-Cre^ERT2^;atg5^f/f^* progeny, tamoxifen was administered by intraperitoneal injections at a dose of 75 mg/kg body weight for five consecutive days starting at P15.

### Chemicals

| Chemical | Final concentration | Function |
| --- | --- | --- |
| NMDA (hellobio, HB0454) | 20 or 50 μM | Agonist of NMDAR |
| DHPG (hellobio, HB0026) | 20 or 50 μM | Agonist of mGLUR1 and 5 |
| Ifeprondil (Co 101244 hydrochloride) (Tocris, 2456) | 10 μM | Selective antagonist of NR2B-containing NMDA receptors |
| SBI-0206965 (Sigma-Aldrich, SML1540) | 500 nM | ULK1 kinase inhibitor |
| Puromycin (Applchem, A2856,0025) | 1 μg/ml | Translation inhibitor |
| Lactacystin | 5 nM | Proteasome inhibitor |
| L-azidohomoalanine (DC Biosciences, CCR-1066) | 4 mM | Analog of Methionine |
| MTEP hydrochloride (hellobio, HB0431) | 10 μM | Antagonist of mGluR5 receptor |
| JNJ16259685 (hellobio, HB0348) | 10 μM | Antagonist of mGluR1 receptor |
| Bafilomycin A1 (Sigma-Aldrich, B1793) | 1.5 nM | Inhibitor of V-ATPase-dependent acidification of lysosomes |
| Biotin-alkyne (PEG4-carboxiamide-propargyl Biotin) (CCR-1266, DC Biosciences) | 40 μM | Biotin conjugate |
| Ciliobrevin-D (Calbiochem, 250401) | 25 μM | Specific blocker of AAA+ ATPase motor cytoplasmic dynein |

### Neuronal Cultures

Cortices and hippocampi were isolated at embryonic day 15.5 (E16.5), rinsed in ice-cold PBS 1X and centrifuged for 5 minutes at 700g at room temperature. Neurons were treated with 0.25% trypsin in PBS 1X for 30 minutes at 37°C, followed by mechanical dissociation. After DMEM (Thermo Fisher Scientific, #41966) and FBS (Biosera) were applied in a one-to-one ratio for trypsin inactivation, the cells were centrifuged 5 minutes at 700g. Neurons were plated at an initial density of 125.000 cells/cm^2^ in 6 cm plates (3 ml/plate) and 12-well plates containing 18-mm glass coverslips (1 ml/plate) – all plates were coated overnight with poly-D-lysine (Sigma-Aldrich, A-003-E). The cells were cultured in Gibco^TM^ Neurobasal^TM^ Medium (Fisher Scientific, A13712-01) supplemented with 2% B-27, 200 mM L-glutamine, 5 mg/ml penicillin and 12.5 mg/ml streptomycin.

### Biochemical purification of synaptosomes

Synaptosomes were isolated as previously described (Carlin et al., 1980). Briefly, brains were isolated from C57BL/6 mice of both sexes at P20 and P25. The brains were rinsed and homogenized in solution A (0.32M Sucrose, 1mM NaHCO3, 1mM MgCl2, 0.5mM CaCl2•H2O, 10mM Na pyrophosphate, nanopure water, protease inhibitors) using a glass homogenizer. Following dilution to 10% weight/volume in solution A, and centrifugation for ten minutes at 710g at 4°C, the pellet was resuspended in solution A supernatant using the homogenizer. After a 10-minute centrifugation at 1.400g at 4°C) for the removal of nuclei, the supernatant was collected and centrifuged for 10 minutes at 13.800g at 4°C. The pellet was resuspended in solution B (0.32M sucrose, 1mM NaHCO3) using the homogenizer and 8 ml of the resulting sample were layered on a discontinuous sucrose gradient (10 ml-layers of 1.2M, 1M and 0.85 M sucrose). Following centrifugation of two hours at 82.500g at 4°C, synaptosomes were isolated between the 1.2M and 1M sucrose layers.

### Biochemical purification of autophagic vesicles from hippocampal slices

200 μm-thick hippocampal sections were isolated from 16 C57BL/6 mice of P22-28 using a vibratome (Leica, VT1200S), in the presence of ice-cold oxygenated a-CSF (125mM NaCl, 3.5mM KCl, 26mM NaHCO3, 1mM MgCl2, 2mM CaCl2, 10mM glucose). The sections were incubated in oxygenated a-CSF and were treated with an NMDA pulse (20μM for ten minutes), followed by two washes in oxygenated sCSF and an one hour incubation in oxygenated a-CSF. After centrifugation for two minutes at 1.000g at 4°C, the supernatant was discarded and the pelleted sections were collected in 10ml of Buffer B (0.25M sucrose, 1mM EDTA pH 8.8, and 20mM Hepes pH 7.4) for homogenization using a glass homogenizer (15 dounces, on ice). Following a centrifugation for two minutes at 2.000g at 4°C, the post-nuclear fraction (PNF) was collected and autophagic vesicles were purified as previously described (Nikoletopoulou et al., 2017). The purified vesicles were subjected to proteinase K (20μg/ml) treatment for twenty minutes on ice to digest proteins associated with the outer membrane. Proteinase K was then inactivated with 4mM PMSF for ten minutes on ice and the material was centrifuged at 16.000g for twenty minutes at 4°C to pellet the autophagic vesicles. Moreover, carbonate extraction for autophagic vesicles was performed, as previously reported (Nikoletopoulou et al., 2017), by incubation of the organelles with ice-cold 0.1M Na2CO3 for thirty minutes on ice followed by a thirty-minute centrifugation at 20psi using an airfuge (GMI, Beckman CLS Ultracentrifuge) at 4°C. The supernatant was precipitated with 10% TCA.

### Study of local protein synthesis in purified synaptosomes

Isolated and purified synaptosomes were pelleted by centrifugation for fifteen minutes at 16.200g at 4°C and resuspended in Medium S buffer (125mM NaCl, 25mM KCl, 0.25mM KH2PO4, 5mM NaHCO3, 100μM CaCl2, 10mM glucose). After a ten-minute incubation in Medium S, they were treated or not with a NMDA- or DHPG-pulse (50μM for ten minutes) at room temperature. Puromycin was added in parallel with the treatments to block protein synthesis. After the treatments, synaptosomes were incubated in Medium S supplemented with the translation inhibitor at room temperature for different time durations (from fifteen to ninety minutes). Synaptosomes were then sonicated to release synaptosomal components and their levels were detected by Western blot analysis.

### Immunolabeling

Cultured cortical and hippocampal neurons were rinsed in PBS 1X and then fixed for fifteen minutes in 4% paraformaldehyde (PFA) in PBS 1X. After fixation, cells were rinsed in PBS and incubated in blocking solution (10% Fetal Bovine Serum (FBS) and 0.2% Triton X-100 in PBS 1X) for 1 hour at room temperature. Neurons were then incubated with primary antibodies diluted in blocking solution at 4°C overnight. The primary antibodies used were: LC3 (1:1000, Santa Cruz, sc-376404), Atg13 (1:1000, Sigma-Aldrich, SAB4200100), MAP2 (1:1000, Synaptic Systems, #188004), FIP200 (1:1000, Cell Signaling, #12436), Atg101 (1:1000, Cell Signaling, #13492), ULK1 (1:1000, Cell Signaling, #8054), p62 (1:2000, Calbiochem, DR1057). Following three rinses with PBS 1X, cells were incubated with secondary antibodies diluted in PBS 1X for 1 hour at room temperature. The secondary antibodies used were: anti-rabbit Alexa 488 (1:1000, Abcam, ab150073), anti-mouse Alexa 594 (1:1000, Abcam, ab150116), and anti-guinea pig Alexa 647 (1:1000, Abcam, ab150187). Hoechst nuclear dye (1:5000) was used for nuclei staining. After three PBS 1X rinses, neurons were mounted onto slides and confocal images of the fluorescently labeled proteins were captured using the Leica TCS SP8 inverted confocal microscope. Similar procedures were followed for - cryosections of brains that were first perfused with 4% PFA and cryoprotected in 30% sucrose in PBS 1X, prior to sectioning.

### Western blotting

Western blots, with slight modifications, were performed as previously described (Nikoletopoulou et al., 2017). Cells, or purified synaptosomes, were collected in cold PBS 1X, incubated in RIPA buffer (500mM Tris-HCl pH 7.4, 150mM NaCl, 1mM EDTA, 1% Triton X-100, 0.1% Na-deoxycholate, 0.1% SDS) supplemented with protease inhibitors (Roche) for 1 hour on ice, and centrifuged (20 minutes at 16.200g, 4°C). Protein samples were separated on a 7.5%, 10% or 15% polyacrylamide gel and transferred to a nitrocellulose membrane (Millipore). Membranes were incubated in 5% milk (or 5% BSA in the case of phospho-Atg13 antibody) for 1 hour at room temperature, and then in 5% milk (or 5% BSA in the case of phospho-Atg13 antibody) with primary antibodies overnight at 4°C. The primary antibodies used were: LC3 (1:1000, Santa Cruz, sc-376404), p62 (1:5000, Calbiochem, DR1057), Atg13 (1:1000, Sigma-Aldrich, SAB4200100), Atg13 pS318 (1:1000, Rockland, #600-401-C49), FIP200 (1:1000, Cell Signaling, #12436), Atg101 (1:1000, Cell Signaling, #13492), ULK1 (1:1000, Cell Signaling, #8054), β III tubulin (Tuj1) (1:5000, Santa Cruz Biotechnology, sc-80005), actin (2Q1055) (1:5000, sc-58673), Atg5 (1:1000, abcam, ab109490), Atg16L1 E-10 (1:1000, Santa Cruz, sc-393274), GRP78Bip (1:1000, Abcam, ab21685), biotin (1:1000, Vector Labs, MB9100), cofilin-1 (1:1000, Proteintech, #66057-1-Ig), GRIA1 (1:1000, Milipore, #AB1504), GRIA2/3 (1:1000, Milipore, #AB1506). Following three 10-minute rinses with PBS-T 1X (100 mM Na2HPO4, 100 mM ΝαΗ2ΡΟ4, 0.5Ν NaCl, 0.1% Tween-20), or TBS-T 1X (150 mM NaCl, 20 mM Tris-HCl pH 7.4, 0.1% Tween-20) in the case of phospho-Atg13, membranes were incubated in 2% milk with the corresponding secondary horseradish peroxidase-conjugated antibodies (1:10.000, Abcam) for 1 hour at room temperature. After three 10-minute rinses with PBS-T 1X (or TBS-T 1X for phospho-Atg13), the membranes were developed by chemiluminescence (SuperSignal Chemiluminescent Substrate, West Pico and Femto, ThermoFisher Scientific) according to the manufacturer’s instructions.

### Immunoprecipitation

100 μg of lysates of hippocampal slices which were AHA-treated and subjected to a click-reaction under control or NMDAR-LTD conditions were immunoprecipitated with antibodies against the four components of the ULK1-complex (ULK1, Atg13, Atg101 and FIP200) with rotation overnight at 4°C, followed by western blot analysis with an antibody biotin (Vector labs, MB9100).

### TUNEL assay

TUNEL staining was performed using the *in situ* cell death detection kit (Roche), following manufacturer’s instructions.

### Measurement of proteasome activity

Proteasome activity was measured in 30 μg mouse forebrain lysates (hippocampus and cortex) or in 50 μg of cortical neuronal lysates from P25-P30 *atg5^f/f^*;*nestin-Cre* and *atg5^f/f^* mice, using the 20S Proteasome Activity Assay Kit (Millipore, APT280) according to manufacturer’s instructions. In both cases, lactacystin a proteasome inhibitor was used as a negative control to block the proteasome activity.

### *Ex vivo* hippocampal preparations for staining and bio-orthogonal labeling

Free-floating hippocampal sections of 100μm were prepared from P25 male C57BL/6 mice. The brain sections were kept in aCSF with simultaneous oxygen supply prior and during treatment. Sections were incubated with or without 50 μM NMDA or DHPG for 10 minutes. After treatment the slices were allowed for 20 minutes in aCSF in order the protein synthesis to take place. After fixation with 4% paraformaldehyde (PFA) and permeabilization overnight with 0.5% Triton X-100 in PBS 1X, non-specific binding sites were blocked for 5 hours using 20% BSA in PBS 1X, incubated overnight with primary antibodies (as described under immunolabeling section) and then with secondary antibodies for 5 hours with PBS 1X. Finally, sections were mounted in 80% glycerol and images were acquired using Leica TCS SP8 inverted confocal microscope.

### AHA labelling in *ex vivo* hippocampal preparations

Hippocampal sections, prepared as mentioned above, were labeled with 4mM AHA (DC Biosciences) according to (Tom Dieck et al., 2012). AHA was supplemented during treatment with or without NMDA and DHPG and for twenty minutes after the treatments. The sections were washed with PBS-MC (PBS 1X, pH 7.4, 1mM MgCl2, 0.1mM CaCl2), fixed in 4% paraformaldehyde (PFA), washed again in PBS 1X and finally permeabilized in 0.5% Triton X-100 in PBS 1X overnight at 4°C. The next day the sections were blocked (4% Fetal Bovine Serum (FBS) in PBS 1X) for five hours at room temperature, calibrated in PBS 1X pH 7.8 and then Click reaction (DC Biosciences) was added to the sections, according to manufactures’ instructions, for overnight incubation. The reaction was washed with 0.5% Triton X-100 in PBS 1X, blocked in 20% BSA in PBS 1X for five hours and primary antibodies: a-Biotin 1:1000, a-Arc 1:1000 (Synaptic systems, 156003), a-Atg13, a-Atg101, a-ULK1 1:500, a-MAP2 1:2000) were added for overnight incubation. Following three washes with PBS 1X, secondary antibodies were added for 5 hours at room temperature and sections were mounted with 80% glycerol. Images were acquired using Leica TCS SP8 inverted confocal microscope.

### Electrophysiological recordings

Electrophysiological experiments were performed in brains of P22 male mice that were isolated and sectioned in 400 μm thick slices using a vibratome (Leica, VT1000S, Leica Biosystems GmbH, Wetzlar, Germany) and transferred to a submerged chamber that was continuously superfused with oxygenated (95% O2/5% CO2) aCSF containing: 124mM NaCl, 3mM KCl, 26mM NaHCO3, 2mM CaCl2, 1mM MgSO4, 1.25mM NaH2PO4 and 10mM glucose (pH 7.4, 315mOsm/l) at room temperature (namely control aCSF). The slices were allowed to recover for at least 2 hours, and then transferred to a submerged recording chamber, which continuously superfused oxygenated (95% O2/5% CO2) aCSF containing: 124mM NaCl, 3mM KCl, 26mM NaHCO3, 2mM CaCl2, 1mM MgSO4, 1.25mM NaH2PO4 and 10mM glucose (pH 7.4, 315 mOsm/l) at room temperature. Extracellular recording electrodes filled with 3M NaCl were placed in the *stratum radiatum* (SR) layer of the CA1 region. Platinum/iridium metal microelectrodes (Harvard apparatus UK, Cambridge, UK) were also placed in the SR layer, about 300 mm away from the recording electrode, and were used to evoke fEPSPs. The voltage responses were amplified using a Dagan BVC-700A amplifier (Dagan Corporation, Minneapolis, MN, USA), digitized using the ITC-18 board (Instrutech) on a PC using custom-made procedures in IgorPro (Wavemetrics, Lake Oswego, OR, USA). The electrical stimulus consisted of a single square waveform of 100 msec duration given at an intensity that generated 40% of the maximum fEPSP, using a stimulator equipped with a stimulus isolation unit (World Precision Instruments). Data were acquired and analyzed using custom-written procedures in IgorPro software (Wavemetrics, Lake Oswego, OR, USA). The voltage response was analyzed in order to measure the fEPSP peak and slope.

For LFS-induced LTD, baseline responses were monitored for at least ten minutes, then stimulated at 1Hz for fifteen minutes with 900 pulses. The fEPSP was monitored for at least fifty minutes following the end of the LFS protocol. In order to test the contribution of autophagy to LFS-induced LTD, brain slices were either incubated in aCSF with the selective autophagy inhibitor (SBI-0206965, 500nM) (Egan et al., 2015).

Brain slices were incubated in aCSF with the dynein inhibitor Ciliobrevin-D (25μM) for at least 20 minutes before, during the LFS stimulation and for at least 50 minutes following the end of the LFS protocol.

For mGluR-LTD, DHPG (20μM) was applied for ten minutes in the aCSF during the recording and following at least ten minutes of monitoring baseline responses. The fEPSP was recorded for at least fifty minutes following wash-out of DHPG.

The fEPSP peak of each response was normalized to the average ten minutes pre-LFS, pre-NMDA or pre-DHPG average fEPSP slope. Statistical analyses were performed by multi-way ANOVA. The average normalized fEPSP peak forty-fifty minutes following each LTD-inducing stimuli was compared across the various conditions using unpaired t-tests.

### Electron Microscopy of purified AVs

Purified forebrain AVs (15mg) were thawed from -80°C and pelleted by centrifugation at 16.000g for thirty minutes at 4°C. The pellet of AVs was resuspended and incubated for twenty minutes in 2.5% glutaraldehyde in 0.1M cacodylate buffer. The AVs were washed for twenty minutes twice in 0.1M cacodylate buffer and post-fixed in 1% OsO4 in 0.1M cacodylate buffer. After two washes of twenty minutes each in 0.1M cacodylate buffer the pellet was dehydrated in a series of increasing concentrations of ethanol (30%, 50%, 70%, 90% and 100%) for ten minutes each time. Two more washes of fifteen min each in absolute ethanol followed at room temperature and two washes in propylene oxide for twenty minutes each. Then, the pellet of AVs was treated in 1:3, 1:1, 3:1 of epoxy resin:propylenoxide, for one hour per case at room temperature and in the end the AVs were resuspended in pure epoxy resin for 1 hour at room temperature. Finally, the AVs were transferred in pure epoxy resin for an overnight incubation by mild shaking. Next day, the AVs were transferred again in pure epoxy resin for one hour incubation at room temperature and then in molds with fresh resin for forty-eight hours at 60°C overnight for their analysis by electron microscopy. Ultrathin 70nm-thick sections were placed on pioloform-coated copper slot grids, stained with lead nitrate and were observed using a JEM-2100 transmission electron microscope (JEOL Ltd, Akishima, Tokyo, JAPAN) at 80kV. Photographs were taken with an Orius camera (Gatan, Pleasanton, CA, USA). Electron micrographs were analyzed using the open source ImageJ software.

### Neurobiotin-labeling and electron microscopy of hippocampal neurons

In a series of experiments, the recording electrode was filled with 1.5% neurobiotin in 3M NaCl and the recording in slices was carried out as previously described. After the recording, the slices were fixed overnight in a fixative containing 4% paraformaldehyde in 0.1M phosphate buffer (PB). After several washes in PB, the slices were embedded in 4% low melting agarose and cut in 70µm thick sections in a vibrating microtome (Leica, VT1000S). Sections around the tip of the recording electrode were collected and cryoprotected in 10% and 20% sucrose solutions in PB, freezed-thawed in liquid nitrogen and processed for the visualization of neurobiotin labeled structures as previously described (Kogo et al., 2004). In brief, sections were incubated for approximately sixty-five hours in avidin–biotin–horseradish peroxidase complex (ABC Elite kit, 1:200; Vector), and neurobiotin was visualized with diaminobenzidine (0.5 mg/mL) and 0.02% H2O2 as substrate. Sections were treated with 1.33% OsO4 and contrasted in 1% uranyl acetate, dehydrated in a series of ethanol and propylene oxide and flat embedded in epoxy resin (Durcupan ACM, Fluca,

Sigma-Aldrich, Gillingham, UK) on microscope slides and. After polymerization of resin (60°C, overnight), regions of interest were reembedded in epoxy resin blocks. Serial electron microscopic sections (70–75nm) were collected on pioloform-coated copper slot grids, contrasted with lead and observed using a JEM-2100 transmission electron microscope (JEOL Ltd, Akishima, Tokyo, JAPAN) at 80kV. Photographs were taken with an Orius camera (Gatan, Pleasanton, CA, USA). Electron micrographs were analyzed using the open source ImageJ software.

### Sample Preparation and Mass Spectrometric Identification of Proteins of purified autophagic vesicles

For the mass spectrometric comparison of purified AVs from control and LTD conditions, membrane enriched fractions were reconstituted in 40µl and soluble TCA-precipitated cargo enriched fractions in 25µl of 1% SDS in 100mM Hepes/NaOH, pH 8.4 and protease inhibitor cocktail (Roche, #11873580001). 20µl of lysates were subjected to an in-solution tryptic digest using a modified version of the Single-Pot Solid-Phase-enhanced Sample Preparation (SP3) protocol(Hughes et al., 2014; Moggridge et al., 2018). Here, lysates were added to Sera-Mag Beads (Thermo Scientific, #4515-2105-050250, 6515-2105-050250) in 10µl 15% formic acid and 30 µl of ethanol. Binding of proteins was achieved by shaking for fifteen minutes at room temperature. SDS was removed by 4 subsequent washes with 200µl of 70% ethanol. Proteins were digested with 0.4 µg of sequencing grade modified trypsin (Promega, #V5111) in 40 µl Hepes/NaOH, pH 8.4 in the presence of 1.25 mM TCEP and 5 mM chloroacetamide (Sigma-Aldrich, #C0267) overnight at room temperature. Beads were separated, washed with 10µl of an aqueous solution of 2% DMSO and the combined eluates were dried down. Peptides were reconstituted in 10µl of H2O and reacted with 80µg of TMT10plex (Thermo Scientific, #90111) (Werner et al., 2014) label reagent dissolved in 4µl of acetonitrile for one hour at room temperature. Excess TMT reagent was quenched by the addition of 4µl of an aqueous solution of 5% hydroxylamine (Sigma-Aldrich, 438227). Peptides were mixed to achieve a 1:1 ratio across all TMT-channels. Mixed peptides were subjected to a reverse phase clean-up step (OASIS HLB 96-well µElution Plate, Waters #186001828BA) and analyzed by LC-MS/MS on a Q Exactive Plus (Thermo Scentific) as previously described (Becher et al., 2018).

Briefly, peptides were separated using an UltiMate 3000 RSLC (Thermo Scientific) equipped with a trapping cartridge (Precolumn; C18 PepMap 100, 5lm, 300lm i.d. × 5mm, 100A°) and an analytical column (Waters nanoEase HSS C18 T3, 75 lm × 25 cm, 1.8 lm, 100 A°). Solvent A: aqueous 0.1% formic acid; Solvent B: 0.1% formic acid in acetonitrile (all solvents were of LC-MS grade). Peptides were loaded on the trapping cartridge using solvent A for three minutes with a flow of 30 µl/minutes. Peptides were separated on the analytical column with a constant flow of 0.3 µl/minutes applying a two hour gradient of 2 – 28% of solvent B in A, followed by an increase to 40% B. Peptides were directly analyzed in positive ion mode applying with a spray voltage of 2.3 kV and a capillary temperature of 320°C using a Nanospray-Flex ion source and a Pico-Tip Emitter 360 lm OD × 20 lm ID; 10 lm tip (New Objective). MS spectra with a mass range of 375–1.200 m/z were acquired in profile mode using a resolution of 70.000 [maximum fill time of 250ms or a maximum of 3e6 ions (automatic gain control, AGC)]. Fragmentation was triggered for the top 10 peaks with charge 2–4 on the MS scan (data-dependent acquisition) with a 30-second dynamic exclusion window (normalized collision energy was 32). Precursors were isolated with a 0.7 m/z window and MS/MS spectra were acquired in profile mode with a resolution of 35,000 (maximum fill time of 120 ms or an AGC target of 2e5 ions).

Acquired data were analyzed using IsobarQuant (Franken et al., 2015) and Mascot V2.4 (Matrix Science) using a reverse UniProt FASTA Mus musculus database (UP000000589) including common contaminants. The following modifications were taken into account: Carbamidomethyl (C, fixed), TMT10plex (K, fixed), Acetyl (N-term, variable), Oxidation (M, variable) and TMT10plex (N-term, variable). The mass error tolerance for full scan MS spectra was set to 10ppm and for MS/MS spectra to 0.02 Da. A maximum of two missed cleavages were allowed. A minimum of two unique peptides with a peptide length of at least seven amino acids and a false discovery rate below 0.01 were required on the peptide and protein level (Savitski et al., 2015).

### Behavioral analyses

*Attentional set-shifting test (AST).* Control (*atg5^f/+^*), *nestin-cre* and *nestin-cHET* male mice, 2 months old, were handled by the experimenter and food-restricted to 90% of their initial weight before the start of the experiment. In addition, mice were habituated to the test chamber and were trained to dig through the bowls in order to get their reward. In the first stage (simple discrimination, SD) of the task two different bedding materials were provided (smoking pipe cleaning rod and silver thread) one of which had the reward (smoking pipe cleaning rod). In the second stage (compound discrimination, CD), a new dimension was introduced using two different odors (strawberry and lavender). During this stage, the same substrate still remained relevant to the reward while the odors are irrelevant. The third stage (compound discrimination reversal, CDR) used the same substrates and odors as the second phase, but the reward was in the other substrate. During the fourth stage (intradimentional shift I, IDI), two new digging media (cardboard and wool) and two new odors (jasmine and apple) are introduced. One of the digging media is still the relevant variable in this phase (cardboard). The next stage (intradimentional shift II, IDII) uses again two new substrates (colorful gobbled paper and cotton) and two new odors (vanilla and cherry). This time the gobbled paper is the rewarded digging media. In the 6^th^ stage (intradimentional reversal, IDR), the correct and the wrong digging media are reversed while vanilla and cherry odors are irrelevant to the reward. In the last stage (extradimentional shift, EDS), the reward now is governed by the odors. The digging media (confetti and cloth) and the odors (ocean and freesia) used in this stage are introduced for the first time in this experiment with the ocean odor being the relevant to the reward variable.

*Open field test.* Control (*atg5f/+*) and *nestin-cHET* male mice, 2 months old, were habituated at least one hour before the experimental procedure. Anxiety-like behavior was measured in an open field box, which consists of four sections: center, periphery, wall and edges. Each mouse was placed individually into the center of the open field box and was allowed to freely explore the open field for fifteen minutes. The experiment was recorded and exploratory behavior was expressed by the percentage of the number of entries in the center and periphery over the number of entries in the center, periphery, edges and wall (JWatcher V1.0).

*Light/dark test.* Unconditioned anxiety response in control (*atg5f/+*) and *nestin-cHET* male mice, 2 months old, was examined by testing their preference for darker over lighter areas. For this purpose, a two-compartment device, consisting of a dark chamber and a light chamber, was used with a small opening in the separator wall between the two chambers that allowed the mice to move freely between the two compartments. Mice were placed in the behavioral room at least one hour before experimentation. Preference was recorded ten seconds after placing each mouse individually in the dark chamber and for a duration of five minutes. Anxiety was calculated as the percentage of the number of entries in the light compartment over the total number of entries in the dark as well as in the light compartment (JWatcher V1.0).

*Elevated Plus Maze (EPM).* Control (*atg5f/+*) and *nestin-cHET* male mice, two months old, were tested in their tendency to be thigmotaxic by using an elevated, plus-shaped maze with two open and two enclosed arms. The mice were firstly placed in the behavioral room to habituate at least one hour before examination. Each mouse was individually placed on the intersection facing a closed arm, opposite of the examiner and the task lasted for five minutes, immediately after the mouse placed on the device. Anxiety was then expressed as the percentage of the number of entries into the open arms over the total entries in both opened and closed arm regions (JWatcher V1.0).

### Statistical analysis

Analyses were performed by student’s t-test or ANOVA or Man Whitney U test, as indicated.

