## Supplement for "Long-term synaptic depression triggers local biogenesis of autophagic vesicles in dendrites and requires autophagic degradation"

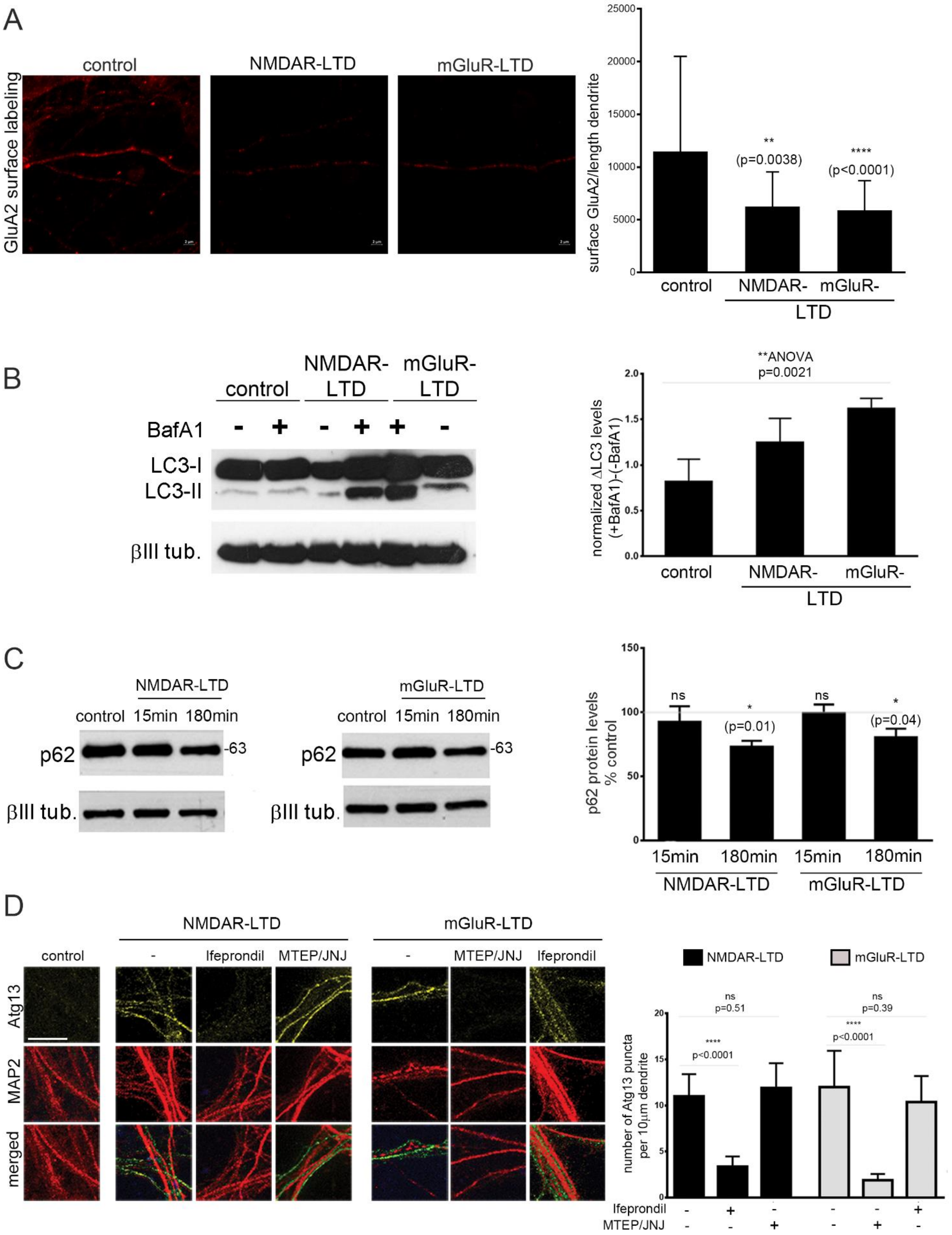

A
control
NMDAR-LTD
mGluR-LTD
GluA2 surface labeling
surface GluA2/length dendrite
**
(p=0.0038)
****
(p<0.0001)
control
NMDAR-
mGluR-
LTD
B
control
NMDAR-LTD
mGluR-LTD
BafA1
LC3-I
LC3-II
βIII tub.
**ANOVA
p=0.0021
normalized ΔLC3 levels (+BafA1)-(-BafA1)
control
NMDAR-
mGluR-
LTD
C
NMDAR-LTD
control 15min 180min
mGluR-LTD
control 15min 180min
p62
-63
βIII tub.
p62 protein levels % control
ns
*
(p=0.01)
ns
*
(p=0.04)
15min 180min 15min 180min
NMDAR-LTD
mGluR-LTD
D
NMDAR-LTD
mGluR-LTD
control
-
Ifeprondil
MTEP/JNJ
-
MTEP/JNJ
Ifeprondil
Atg13
MAP2
merged
NMDAR-LTD
mGluR-LTD
ns
p=0.51
****
p<0.0001
ns
p=0.39
****
p<0.0001
number of Atg13 puncta per 10μm dendrite
Ifeprondil
MTEP/JNJ

**Supplementary Figure 1. Chemical LTD triggers autophagy in cultured neurons.**

(A) Surface labeling for GluA2 in control neurons or one hour following an NMDA or DHPG pulse to induce chemical NMDAR- and mGLUR-LTD respectively.

(B) Western blot analysis for LC3 and β-III tubulin in lysates prepared from neurons that were exposed to control conditions or to a five-minute pulse 50μM DHPG, and in the presence or absence of 1.5nM Bafilomycin A1 for one hour after the pulse. Lysates were obtained one hour after the pulse. Graph shows the autophagic flux in the indicated conditions, calculated as the difference of LC3-II levels in the presence and absence of Bafilomycin A1. Bars represent mean values +/- SEM. (N=3 independent experiments).

(C) Western blot analysis in lysates of control neurons or fifteen and one hundred eighty minutes after an NMDA or DHPG pulse with an antibody against p62 and normalized for β-III tubulin. Graph shows the normalized levels of p62 in the indicated conditions. (N=3 independent experiments). Bars represent mean values +/- SEM.

(D) Representative confocal images of cultured neurons under control conditions or 15 minutes after chemical NMDAR- or mGluR-LTD, immunostained with an antibody against Atg13 to label autophagic structures and MAP2 to label dendrites. (Scale bar: 50μm). Neurons were either control, fifteen minutes after NMDAR-LTD and mGluR-LTD or with treatment for one hour before the NMDA or DHPG pulse with Ifeprondil or MTEP and JNJ to pharmacologically inhibit NR2B and mGluR1/5 receptors, respectively. (Scale bar: 25μm). Graph showing the number of dendritic Atg13-positive puncta, normalized to the dendritic length, in the indicated conditions. (N=3 independent experiments per condition).

Unless indicated otherwise, statistical analyses were performed using student’s t-test.

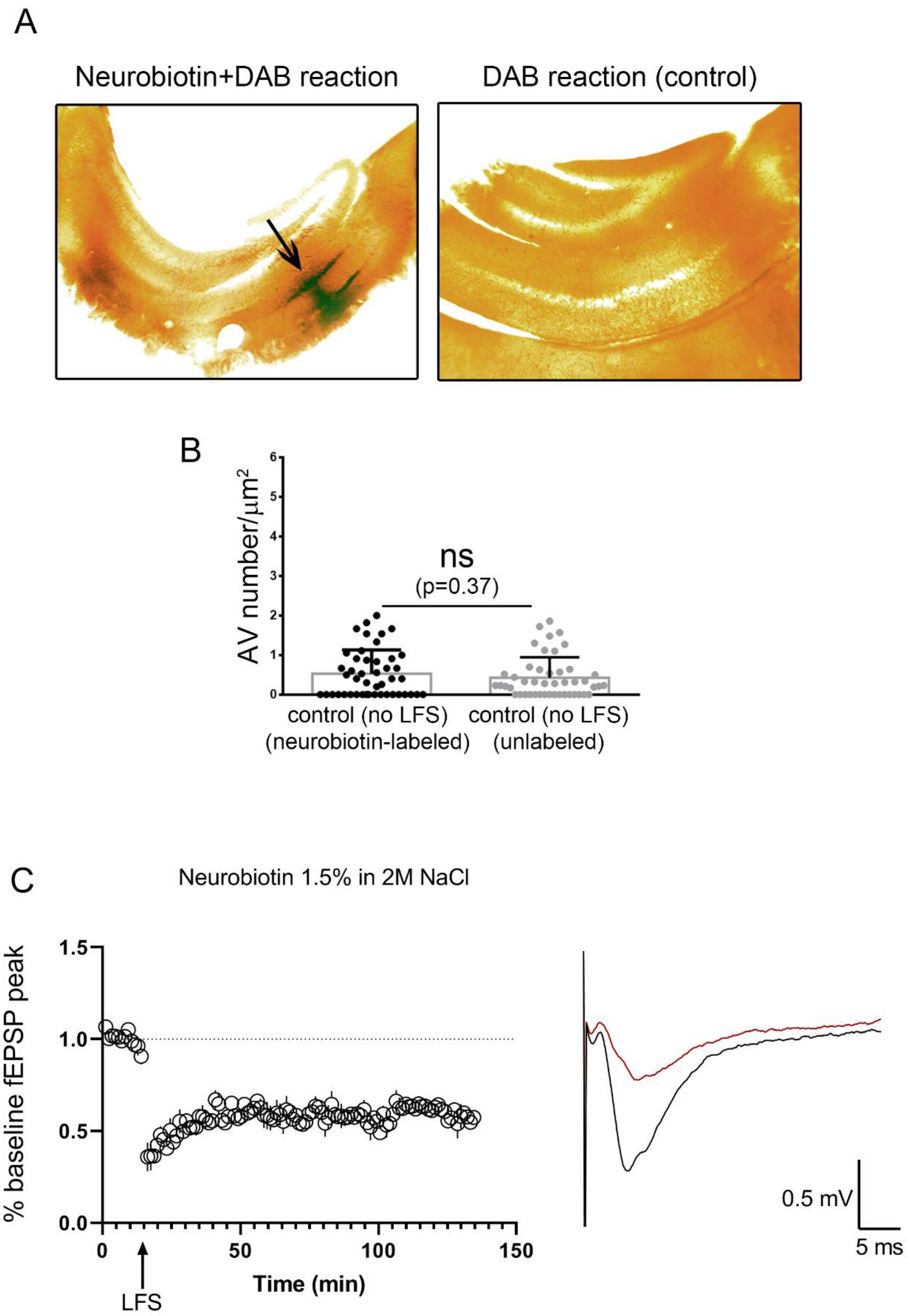
A
Neurobiotin+DAB reaction
DAB reaction (control)
B
AV number/μm²
ns
(p=0.37)
control (no LFS)
(neurobiotin-labeled)
control (no LFS)
(unlabeled)
C
Neurobiotin 1.5% in 2M NaCl
% baseline fEPSP peak
Time (min)
LFS
0.5 mV
5 ms

**Supplementary Figure 2. Neurobiotin labeling of dendrites during LTD induction is restricted to the CA1 of hippocampus and does not interfere with LTD.**

(A) DAB reaction in neurobiotin-labeled or control hippocampal slices. In both cases, the brain slices were fixed 60 minutes after input-output stimulation.

(B) Graph showing the density of AVs in neurobiotin-labeled or unlabeled distal post-synaptic dendrites of pyramidal neurons in the *stratum radiatum* of CA1 in control (without LTD) hippocampal slices. Bars represent mean values +/- SEM. (N=46 dendrites per condition). Statistical analyses were performed using Man Whitney U test.

(C) Time-plot of the normalized fEPSPs and representative traces before and after LFS stimulation of neurobiotin-labelled CA1 area. Note that neurobiotin-labeling does not interfere with LTD induction and maintenance.

A

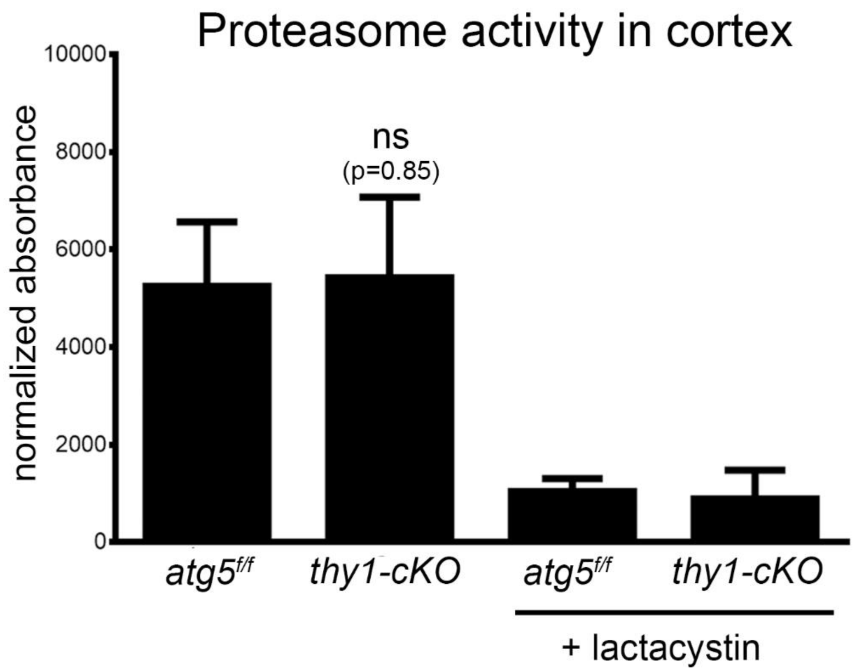
Proteasome activity in cortex
normalized absorbance
10000
8000
6000
4000
2000
0
ns
(p=0.85)
atg5f/f
thy1-cKO
atg5f/f
thy1-cKO
+ lactacystin

B

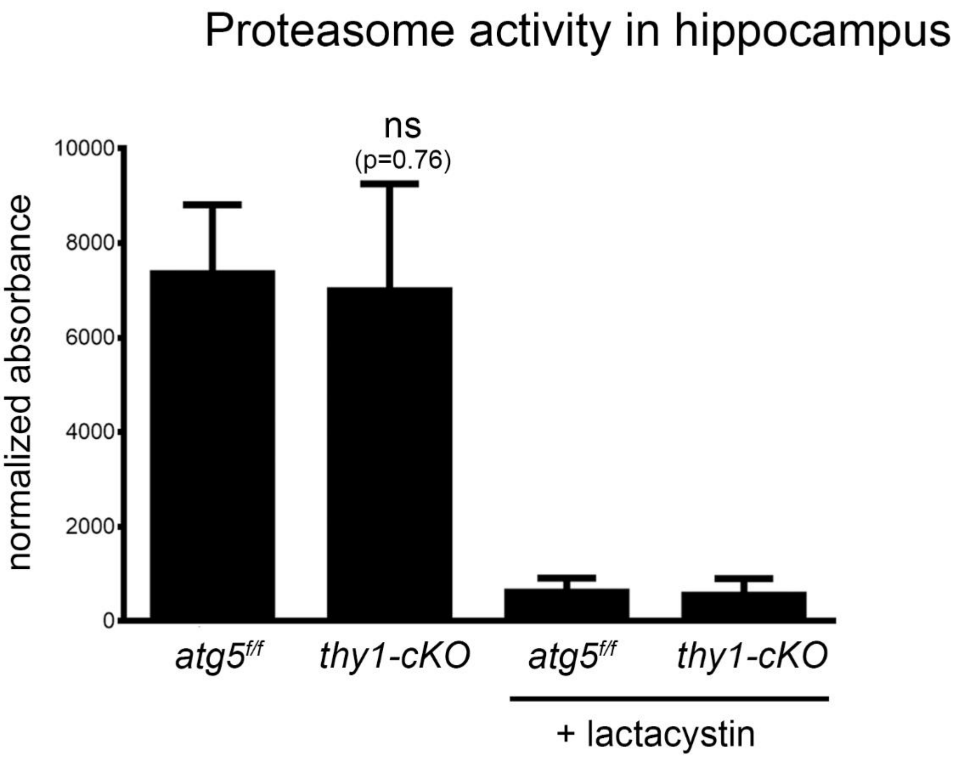
Proteasome activity in hippocampus
normalized absorbance
10000
8000
6000
4000
2000
0
ns
(p=0.76)
atg5f/f
thy1-cKO
atg5f/f
thy1-cKO
+ lactacystin

**Supplementary Figure 3. Proteasome activity in the forebrain is not affected by genetic ablation of autophagy.**

Proteasome activity in (A) cortical and (B) hippocampal lysates of *atg5*$^{f/f}$, and *thy1-cKO* animals at postnatal day forty (P40) in the presence or absence of proteasomal inhibitor lactacystin. Bars represent mean values +/-SEM. (N=3 animals per genotype). Statistical analysis was performed using student's t-test.

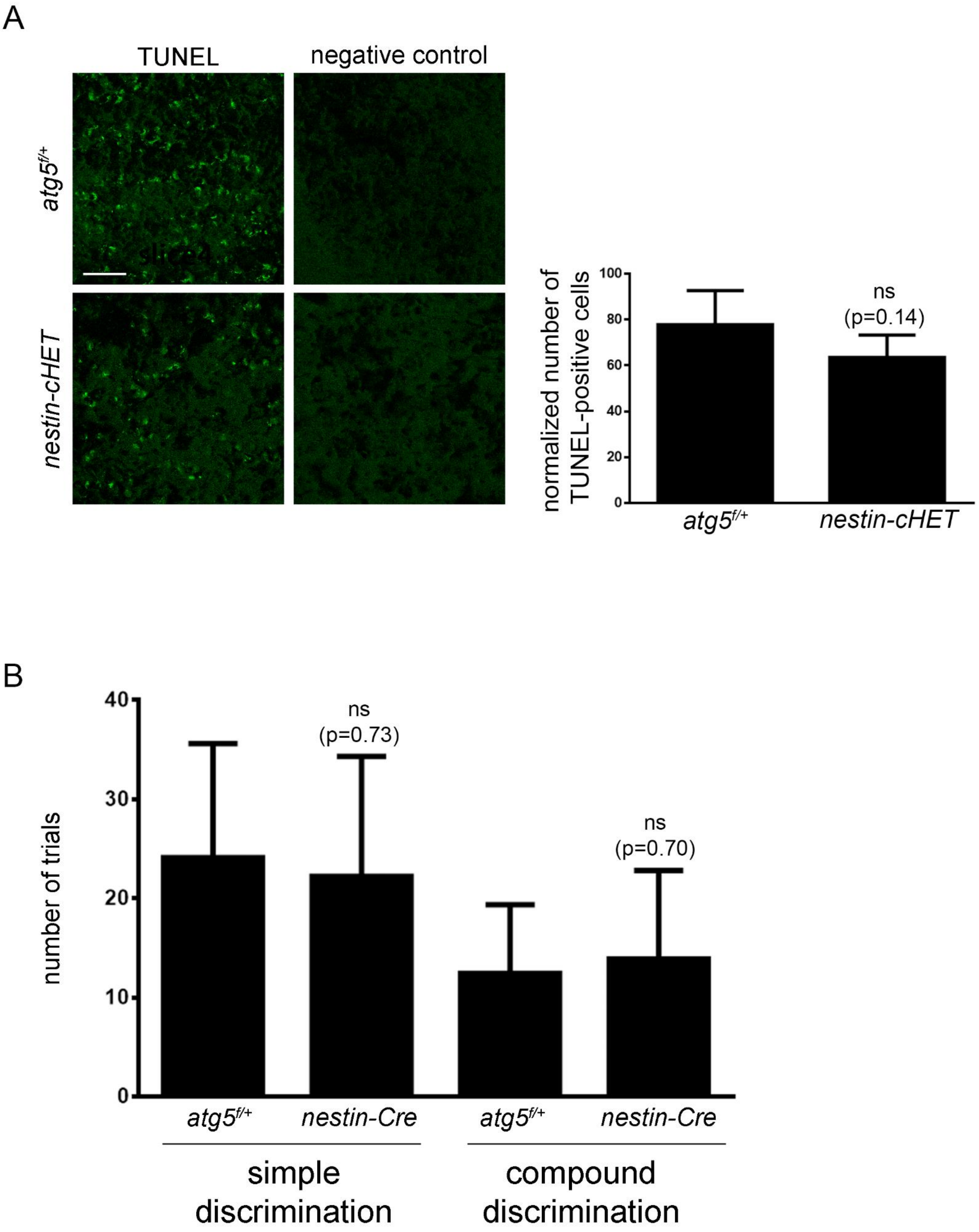
A
TUNEL
negative control
atg5f/+
nestin-cHET
normalized number of TUNEL-positive cells
100
80
60
40
20
0
ns
(p=0.14)
atg5f/+
nestin-cHET
B
number of trials
40
30
20
10
0
ns
(p=0.73)
ns
(p=0.70)
atg5f/+
nestin-Cre
atg5f/+
nestin-Cre
simple discrimination
compound discrimination

**Supplementary Figure 4. Characterization of *nestin-cHET* animals.**

(A) Representative images of TUNEL assay of *atg5*$^{f/+}$ and *nestin-cHET* forebrain cryosections (Scale bar: 50μm). Graph shows the normalized TUNEL-positive cells in the forebrain in the two genotypes. Bars represent mean values +/-SEM (N=3 independent experiments). Statistical analysis was performed using student's t-test.

(B) Simple and compound discrimination tasks in *atg5*$^{f/+}$ and nestin-cHET animals. Graph shows the number of trials mice performed in the each task. (N=8 animals per genotype). Statistical analyses were performed using student's t-test.

### Supplementary Table Legends

**Supplementary Table 1. The dynamic autophagic cargo under LTD.** List of proteins comprising the dynamic cargo of autophagosomes after NMDAR-LTD. Proteins are identified as hits, candidates or trends (according to their FDR value) in both supernatant and pellet fractions.

**Supplementary Table 2. The ASD-implicated proteins of the dynamic LTD autophagic cargo.** List of ASD-implicated proteins identified in the dynamic autophagic cargo under NMDAR-LTD. Scores are based on the gene scoring module within SFARI Gene 3.0 (https://gene.sfari.org) (S for syndromic and scores 1>2>3>4>5).
