## Supplementary material for "Long-term synaptic depression triggers local biogenesis of autophagic vesicles in dendrites and requires autophagic degradation": table_S1

Table_1

| gene_name | increased or decreased in LTD | protein_id | description | keep_supernatant | keep_pellet | comparison | sample | hit_annota | pvalue | fdr | hit | hit_annotation |
|---|---|---|---|---|---|---|---|---|---|---|---|---|
| 2010300C02RIK | increased | E9Q3M9 | Protein 2010300C02Rik OS=Mus musculus GN=2010300C02Rik PE=1 SV=1 | TRUE | TRUE | NMDA - control | pellet | fdrtool | 0,074667 | 0,584087 | FALSE | trend |
| 2310035C23RIK\|KIAA | increased | A0A087WSS1\|E9QM90\|Q148V7\|Q148V7-2 | Protein 2310035C23Rik OS=Mus musculus GN=2310035C23Rik PE=1 SV=1\|Protein 2310035C23Rik O | TRUE | FALSE | NMDA - control | supernatant | fdrtool | 0,080056 | 0,589077 | FALSE | trend |
| ABR | increased | E9PUE7\|Q5SSL4\|Q5SSL4-2\|Q5SSL4-3\|Q5SSL4-4 | Active breakpoint cluster region-related protein OS=Mus musculus GN=Abr PE=1 SV=1\|Isoform 2 of | TRUE | TRUE | NMDA - control | supernatant | fdrtool | 0,08128 | 0,592743 | FALSE | trend |
| ADAM22 | increased | D3YUP9\|Q9R1V6\|Q9R1V6-10\|Q9R1V6-11\|Q9R1V6-12\|Q9R1V6-13\|Q9R1V6-14\|Q9R1V6-15\|Q9R1V6-17\|Q9 | Disintegrin and metalloproteinase domain-containing protein 22 OS=Mus musculus GN=Adam22 PE= | FALSE | TRUE | NMDA - control | pellet | fdrtool | 0,037245 | 0,476812 | FALSE | trend |
| ADAM23 | increased | Q5SRA0\|Q9R1V7\|Q9R1V7-2\|Q9R1V7-3\|Q9R1V7-4 | Disintegrin and metalloproteinase domain-containing protein 23 OS=Mus musculus GN=Adam23 PE= | FALSE | TRUE | NMDA - control | pellet | fdrtool | 0,027266 | 0,420425 | FALSE | trend |
| ADCY5 | increased | P84309\|P84309-2 | Adenylate cyclase type 5 OS=Mus musculus GN=Adcy5 PE=1 SV=2\|Isoform 2 of Adenylate cyclase ty | FALSE | TRUE | NMDA - control | pellet | fdrtool | 0,032478 | 0,452473 | FALSE | trend |
| ADD2 | increased | Q9QYB8\|Q9QYB8-2 | Beta-adducin OS=Mus musculus GN=Add2 PE=1 SV=4\|Isoform 2 of Beta-adducin OS=Mus musculus | TRUE | TRUE | NMDA - control | pellet | fdrtool | 0,000394 | 0,052021 | FALSE | candidate |
| ADRBK1 | increased | Q3U1V3\|Q7TS64\|Q99MK8 | Protein-serine/threonine kinase OS=Mus musculus GN=Adrbk1 PE=1 SV=1\|Protein-serine/threonine | TRUE | TRUE | NMDA - control | supernatant | fdrtool | 0,020215 | 0,265777 | FALSE | trend |
| AGPAT1 | increased | A0A0R4J263\|O35083 | 1-acyl-sn-glycerol-3-phosphate acyltransferase OS=Mus musculus GN=Agpat1 PE=1 SV=1\|1-acyl-sn-g | FALSE | TRUE | NMDA - control | pellet | fdrtool | 0,070833 | 0,577093 | FALSE | trend |
| AK5 | increased | Q920P5 | Adenylate kinase isoenzyme 5 OS=Mus musculus GN=Ak5 PE=1 SV=2 | TRUE | TRUE | NMDA - control | supernatant | fdrtool | 0,008435 | 0,144167 | FALSE | candidate |
| ALDH1B1 | increased | Q9CZS1 | Aldehyde dehydrogenase X, mitochondrial OS=Mus musculus GN=Aldh1b1 PE=1 SV=1 | TRUE | TRUE | NMDA - control | pellet | fdrtool | 0,035821 | 0,469967 | FALSE | trend |
| ANK2 | increased | Q8C8R3-2\|Q8C8R3-5\|Q8C8R3-7 | Isoform 2 of Ankyrin-2 OS=Mus musculus GN=Ank2\|Isoform 5 of Ankyrin-2 OS=Mus musculus GN=A | TRUE | TRUE | NMDA - control | supernatant | fdrtool | 0,002543 | 0,06099 | FALSE | candidate |
| ANKS1B | increased | A0A0R4J2A6\|A0A0R4J2A8\|E9QPP6\|Q8BIZ1-2\|Q8BIZ1-4\|S4R1H2\|S4R1Q0\|S4R2H2\|S4R2I2\|S4R2Q2 | Ankyrin repeat and sterile alpha motif domain-containing protein 1B OS=Mus musculus GN=Anks1b | TRUE | TRUE | NMDA - control | supernatant | fdrtool | 0,065806 | 0,540941 | FALSE | trend |
| AP3D1 | increased | O54774 | AP-3 complex subunit delta-1 OS=Mus musculus GN=Ap3d1 PE=1 SV=1 | TRUE | TRUE | NMDA - control | supernatant | fdrtool | 0,002846 | 0,065475 | FALSE | candidate |
| ARF6 | increased | P62331\|Q3U0D7 | ADP-ribosylation factor 6 OS=Mus musculus GN=Arf6 PE=1 SV=2\|ADP-ribosylation factor 6 OS=Mus | TRUE | TRUE | NMDA - control | pellet | fdrtool | 0,065556 | 0,566459 | FALSE | trend |
| ARHGAP35 | increased | B2RTN5\|Q91YM2 | Glucocorticoid receptor DNA binding factor 1 OS=Mus musculus GN=Arhgap35 PE=1 SV=1\|Rho GTPa | TRUE | FALSE | NMDA - control | supernatant | fdrtool | 0,005789 | 0,111323 | FALSE | candidate |
| ARMC10 | increased | Q9D0L7\|Q9D0L7-2 | Armadillo repeat-containing protein 10 OS=Mus musculus GN=Armc10 PE=1 SV=1\|Isoform 2 of Arma | TRUE | TRUE | NMDA - control | supernatant | fdrtool | 0,048767 | 0,46617 | FALSE | trend |
| ASAP1 | increased | E9QMI7\|E9QMJ1\|E9QN63\|H3BJY2\|H3BKD4\|H3BKE6\|H3BL41\|Q9QWY8\|Q9QWY8-2\|Q9QWY8-3\|Q9QWY8- | Arf-GAP with SH3 domain, ANK repeat and PH domain-containing protein 1 OS=Mus musculus GN=A | TRUE | FALSE | NMDA - control | supernatant | fdrtool | 0,001651 | 0,045308 | FALSE | candidate |
| ASPHD2 | increased | Q80VP9 | Aspartate beta-hydroxylase domain-containing protein 2 OS=Mus musculus GN=Asphd2 PE=1 SV=1 | FALSE | TRUE | NMDA - control | pellet | fdrtool | 0,035543 | 0,468588 | FALSE | trend |
| ATP1A1 | increased | Q8VDN2 | Sodium/potassium-transporting ATPase subunit alpha-1 OS=Mus musculus GN=Atp1a1 PE=1 SV=1 | FALSE | TRUE | NMDA - control | pellet | fdrtool | 0,030333 | 0,440063 | FALSE | trend |
| ATP1A3 | increased | A0A0G2JGX4\|Q6PIC6\|Q8VCE0 | Sodium/potassium-transporting ATPase subunit alpha-3 OS=Mus musculus GN=Atp1a3 PE=1 SV=1\|S | TRUE | TRUE | NMDA - control | supernatant | fdrtool | 5,53E-13 | 2,46E-10 | TRUE | hit |
| ATP1B1 | increased | P14094\|Q545P0 | Sodium/potassium-transporting ATPase subunit beta-1 OS=Mus musculus GN=Atp1b1 PE=1 SV=1\|Sc | TRUE | TRUE | NMDA - control | supernatant | fdrtool | 0,000269 | 0,011506 | TRUE | hit |
| ATP2A2 | increased | J3KMM5\|O55143 | Calcium-transporting ATPase OS=Mus musculus GN=Atp2a2 PE=1 SV=2\|Sarcoplasmic/endoplasmic r | FALSE | TRUE | NMDA - control | pellet | fdrtool | 0,042392 | 0,499012 | FALSE | trend |
| ATP5A1 | increased | Q03265 | ATP synthase subunit alpha, mitochondrial OS=Mus musculus GN=Atp5a1 PE=1 SV=1 | TRUE | TRUE | NMDA - control | supernatant | fdrtool | 0,046408 | 0,453855 | FALSE | trend |
| ATP5C1 | increased | Q3UD06\|Q8C2Q8\|Q91VR2 | ATP synthase subunit gamma OS=Mus musculus GN=Atp5c1 PE=1 SV=1\|ATP synthase subunit gamm | TRUE | TRUE | NMDA - control | supernatant | fdrtool | 0,067996 | 0,54906 | FALSE | trend |
| ATP5I | increased | Q06185 | ATP synthase subunit e, mitochondrial OS=Mus musculus GN=Atp5i PE=1 SV=2 | TRUE | TRUE | NMDA - control | supernatant | fdrtool | 0,028229 | 0,335768 | FALSE | trend |
| ATP5J2 | increased | P56135 | ATP synthase subunit f, mitochondrial OS=Mus musculus GN=Atp5j2 PE=1 SV=3 | FALSE | TRUE | NMDA - control | pellet | fdrtool | 0,077262 | 0,588513 | FALSE | trend |
| ATP6V0A1 | increased | K3W4T3\|Q9Z1G4\|Q9Z1G4-2\|Q9Z1G4-3 | Isoform A1-I of V-type proton ATPase 116 kDa subunit a isoform 1 OS=Mus musculus GN=Atp6v0a1\| | TRUE | TRUE | NMDA - control | supernatant | fdrtool | 0,006991 | 0,127214 | FALSE | candidate |
| ATP6V0D1 | increased | P51863 | V-type proton ATPase subunit d 1 OS=Mus musculus GN=Atp6v0d1 PE=1 SV=2 | TRUE | TRUE | NMDA - control | pellet | fdrtool | 0,009367 | 0,232477 | FALSE | trend |
| ATP6V1D | increased | P57746\|Q3U861 | V-type proton ATPase subunit D OS=Mus musculus GN=Atp6v1d PE=1 SV=1\|ATPase, H+ transporting | TRUE | TRUE | NMDA - control | supernatant | fdrtool | 0,075876 | 0,576037 | FALSE | trend |
| ATP8A1 | increased | A0A0M3HEP7\|F6WYQ5\|P70704-2\|P70704-3 | Phospholipid-transporting ATPase OS=Mus musculus GN=Atp8a1 PE=1 SV=1\|Phospholipid-transport | FALSE | TRUE | NMDA - control | pellet | fdrtool | 0,034732 | 0,464494 | FALSE | trend |
| BRAF | increased | F6SZ47\|P28028 | Serine/threonine-protein kinase B-raf (Fragment) OS=Mus musculus GN=Braf PE=1 SV=1\|Serine/thre | TRUE | FALSE | NMDA - control | supernatant | fdrtool | 0,009025 | 0,150508 | FALSE | candidate |
| BSN | increased | O88737\|O88737-2 | Isoform 2 of Protein bassoon OS=Mus musculus GN=Bsn\|Protein bassoon OS=Mus musculus GN=Bs | TRUE | TRUE | NMDA - control | supernatant | fdrtool | 0,001092 | 0,033083 | TRUE | hit |
| CACNA1E | increased | A0A087WS83\|Q61290 | Voltage-dependent R-type calcium channel subunit alpha-1E OS=Mus musculus GN=Cacna1e PE=1 S | FALSE | TRUE | NMDA - control | pellet | fdrtool | 0,06499 | 0,565241 | FALSE | trend |
| CACNA2D1 | increased | O08532-5 | Isoform 2E of Voltage-dependent calcium channel subunit alpha-2/delta-1 OS=Mus musculus GN=Ca | FALSE | TRUE | NMDA - control | pellet | fdrtool | 0,067219 | 0,569946 | FALSE | trend |
| CACNG3 | increased | Q9JJV5 | Voltage-dependent calcium channel gamma-3 subunit OS=Mus musculus GN=Cacng3 PE=2 SV=2 | FALSE | TRUE | NMDA - control | pellet | fdrtool | 0,045532 | 0,510884 | FALSE | trend |
| CADPS2 | increased | E9Q5C0\|E9Q835\|Q8BYR5-2\|Q8BYR5-3\|Q8BYR5-4\|S4R1C6 | Calcium-dependent secretion activator 2 OS=Mus musculus GN=Cadps2 PE=1 SV=2\|Calcium-depend | TRUE | FALSE | NMDA - control | supernatant | fdrtool | 0,015903 | 0,223064 | FALSE | trend |
| CAMK2A | increased | P11798 | Calcium/calmodulin-dependent protein kinase type II subunit alpha OS=Mus musculus GN=Camk2a | TRUE | TRUE | NMDA - control | supernatant | fdrtool | 8,88E-16 | 1,19E-12 | TRUE | hit |
| CAMK2A | increased | P11798 | Calcium/calmodulin-dependent protein kinase type II subunit alpha OS=Mus musculus GN=Camk2a | TRUE | TRUE | NMDA - control | pellet | fdrtool | 0,008652 | 0,226697 | FALSE | trend |
| CAMK2B | increased | P28652\|Q5SVI2\|Q5SVJ0 | Calcium/calmodulin-dependent protein kinase II, beta, isoform CRA b OS=Mus musculus GN=Camk2 | TRUE | TRUE | NMDA - control | supernatant | fdrtool | 0,000435 | 0,016587 | FALSE | candidate |
| CAMK2D | increased | A0A0G2JGS4\|E9Q1T1\|E9Q1W0\|Q6PHZ2-5 | Calcium/calmodulin-dependent protein kinase type II subunit delta OS=Mus musculus GN=Camk2d | TRUE | TRUE | NMDA - control | supernatant | fdrtool | 0,006877 | 0,125771 | FALSE | candidate |
| CAMK2G | increased | Q923T9\|Q923T9-2\|Q923T9-3 | Calcium/calmodulin-dependent protein kinase type II subunit gamma OS=Mus musculus GN=Camk2 | TRUE | TRUE | NMDA - control | supernatant | fdrtool | 7,21E-09 | 1,20E-06 | TRUE | hit |
| CAMKV | increased | A0A0A6YW88\|Q3UHL1 | CaM kinase-like vesicle-associated protein OS=Mus musculus GN=Camkv PE=1 SV=1\|CaM kinase-like | TRUE | TRUE | NMDA - control | supernatant | fdrtool | 0,020688 | 0,270314 | FALSE | trend |
| CDS2 | increased | Q99L43 | Phosphatidate cytidylyltransferase 2 OS=Mus musculus GN=Cds2 PE=1 SV=1 | FALSE | TRUE | NMDA - control | pellet | fdrtool | 0,083221 | 0,598529 | FALSE | trend |
| CISD2 | increased | Q9CQB5 | CDGSH iron-sulfur domain-containing protein 2 OS=Mus musculus GN=Cisd2 PE=1 SV=1 | FALSE | TRUE | NMDA - control | pellet | fdrtool | 0,078948 | 0,591266 | FALSE | trend |
| CIT | increased | D3YU89\|D3Z1U0\|E9QL53\|F6SBR5\|P49025\|P49025-3\|P49025-4\|P49025-5 | Citron Rho-interacting kinase OS=Mus musculus GN=Cit PE=1 SV=1\|Citron Rho-interacting kinase OS | TRUE | TRUE | NMDA - control | supernatant | fdrtool | 0,000639 | 0,021816 | FALSE | candidate |
| CLASP2 | increased | E9Q8N5\|F7DCH5\|Q08EB6\|Q8BRT1 | CLIP-associating protein 2 OS=Mus musculus GN=Clasp2 PE=1 SV=1\|CLIP-associating protein 2 OS=M | TRUE | TRUE | NMDA - control | supernatant | fdrtool | 0,000618 | 0,021315 | FALSE | candidate |
| CORO1A | increased | G3UYK8\|O89053\|Q3U1N0 | Coronin-1A OS=Mus musculus GN=Coro1a PE=1 SV=5\|Coronin OS=Mus musculus GN=Coro1a PE=1 S | TRUE | TRUE | NMDA - control | supernatant | fdrtool | 0,006209 | 0,117073 | FALSE | candidate |
| CORO1C | increased | Q499X7\|Q9WUM4 | Coronin OS=Mus musculus GN=Coro1c PE=1 SV=1\|Coronin-1C OS=Mus musculus GN=Coro1c PE=1 S | TRUE | TRUE | NMDA - control | supernatant | fdrtool | 0,038368 | 0,407245 | FALSE | trend |
| CORO2B | increased | Q8BH44 | Coronin-2B OS=Mus musculus GN=Coro2b PE=1 SV=2 | TRUE | FALSE | NMDA - control | supernatant | fdrtool | 0,066806 | 0,544682 | FALSE | trend |
| COX5B | increased | P19536\|Q9D881 | Cytochrome c oxidase subunit 5B, mitochondrial OS=Mus musculus GN=Cox5b PE=1 SV=1\|Cytochron | TRUE | TRUE | NMDA - control | pellet | fdrtool | 0,005496 | 0,191044 | FALSE | candidate |
| CPT1C | increased | Q8BGD5 | Carnitine O-palmitoyltransferase 1, brain isoform OS=Mus musculus GN=Cpt1c PE=1 SV=1 | FALSE | TRUE | NMDA - control | pellet | fdrtool | 0,046669 | 0,514916 | FALSE | trend |
| CRELD1 | increased | A8C1T7\|Q91XD7 | Cysteine-rich with EGF-like domain protein 1 OS=Mus musculus GN=Creld1 PE=1 SV=1\|Cysteine-rich | FALSE | TRUE | NMDA - control | pellet | fdrtool | 0,057709 | 0,548026 | FALSE | trend |
| CTNNA2 | increased | E0CXB9\|Q61301\|Q61301-2\|Q61301-3 | Catenin alpha-2 OS=Mus musculus GN=Ctnna2 PE=1 SV=1\|Catenin alpha-2 OS=Mus musculus GN=Ct | TRUE | TRUE | NMDA - control | pellet | fdrtool | 0,056565 | 0,545031 | FALSE | trend |
| DCLK1 | increased | A0A0G2JGQ6\|Q9JLM8 | Serine/threonine-protein kinase DCLK1 OS=Mus musculus GN=Dclk1 PE=1 SV=1 | TRUE | TRUE | NMDA - control | supernatant | fdrtool | 8,97E-07 | 8,55E-05 | TRUE | hit |
| DCTN1 | increased | E9Q3M3\|E9Q586\|O08788\|O08788-2 | Dynactin subunit 1 OS=Mus musculus GN=Dctn1 PE=1 SV=1\|Dynactin subunit 1 OS=Mus musculus G | TRUE | TRUE | NMDA - control | supernatant | fdrtool | 0,057958 | 0,509282 | FALSE | trend |
| DGKE | increased | Q9R1C6 | Diacylglycerol kinase epsilon OS=Mus musculus GN=Dgke PE=1 SV=1 | FALSE | TRUE | NMDA - control | pellet | fdrtool | 0,010022 | 0,242525 | FALSE | trend |
| DLAT | increased | Q8BMF4 | Dihydrolipoyllysine-residue acetyltransferase component of pyruvate dehydrogenase complex, mito | TRUE | TRUE | NMDA - control | supernatant | fdrtool | 0,029034 | 0,342066 | FALSE | trend |
| DLG1 | increased | D3Z3B8\|E9Q9H0\|H7BWY4\|Q811D0\|Q811D0-2\|Q811D0-3 | Disks large homolog 1 OS=Mus musculus GN=Dlg1 PE=1 SV=1\|Isoform 2 of Disks large homolog 1 OS | TRUE | TRUE | NMDA - control | supernatant | fdrtool | 0,000607 | 0,02107 | FALSE | candidate |
| DLG4 | increased | Q62108\|Q62108-2\|Q62108-3 | Disks large homolog 4 OS=Mus musculus GN=Dlg4 PE=1 SV=1\|Isoform 2 of Disks large homolog 4 OS | TRUE | TRUE | NMDA - control | supernatant | fdrtool | 0,004417 | 0,090845 | FALSE | candidate |
| DLST | increased | Q9D2G2 | Dihydrolipoyllysine-residue succinyltransferase component of 2-oxoglutarate dehydrogenase comple | TRUE | TRUE | NMDA - control | supernatant | fdrtool | 0,079063 | 0,586051 | FALSE | trend |
| DMXL2 | increased | B0V2P5\|Q8BPN8\|Q8BPN8-2 | DmX-like protein 2 OS=Mus musculus GN=Dmxl2 PE=1 SV=1\|DmX-like protein 2 OS=Mus musculus G | TRUE | TRUE | NMDA - control | supernatant | fdrtool | 0,055403 | 0,498013 | FALSE | trend |
| DNAJA1 | increased | P63037\|Q5NTY0 | DnaJ homolog subfamily A member 1 OS=Mus musculus GN=Dnaja1 PE=1 SV=1\|DnaJ (Hsp40) homol | TRUE | FALSE | NMDA - control | supernatant | fdrtool | 0,06073 | 0,520953 | FALSE | trend |
| ERC2 | increased | Q6PH08\|Q6PH08-2\|Q6PH08-3 | ERC protein 2 OS=Mus musculus GN=Erc2 PE=1 SV=2\|Isoform 2 of ERC protein 2 OS=Mus musculus | TRUE | TRUE | NMDA - control | supernatant | fdrtool | 0,002412 | 0,058918 | FALSE | candidate |
| EXOC3 | increased | Q6KAR6\|Q8K0E2 | Exocyst complex component 3 OS=Mus musculus GN=Exoc3 PE=1 SV=2\|Exocyst complex componen | TRUE | FALSE | NMDA - control | supernatant | fdrtool | 0,04231 | 0,431055 | FALSE | trend |
| EXOC7 | increased | A2AAN0\|O35250\|O35250-2 | Exocyst complex component 7 OS=Mus musculus GN=Exoc7 PE=1 SV=1\|Exocyst complex componen | TRUE | TRUE | NMDA - control | supernatant | fdrtool | 0,034454 | 0,381554 | FALSE | trend |
| FAM81A | increased | Q3UXZ6 | Protein FAM81A OS=Mus musculus GN=Fam81a PE=1 SV=2 | FALSE | TRUE | NMDA - control | pellet | fdrtool | 0,019049 | 0,353177 | FALSE | trend |
| FUS | increased | G3UXT7\|P56959\|Q564D0\|Q8CFQ9 | RNA-binding protein FUS (Fragment) OS=Mus musculus GN=Fus PE=1 SV=1\|RNA-binding protein FUS | FALSE | TRUE | NMDA - control | pellet | fdrtool | 0,000123 | 0,033705 | TRUE | hit |
| FYN | increased | D3YZ57\|P39688\|P39688-2 | Isoform 2 of Tyrosine-protein kinase Fyn OS=Mus musculus GN=Fyn\|Tyrosine-protein kinase Fyn OS | TRUE | TRUE | NMDA - control | supernatant | fdrtool | 0,010227 | 0,162544 | FALSE | candidate |
| GAS7 | increased | B1ATI9\|Q3U432\|Q60780\|Q60780-2 | Growth arrest-specific protein 7 OS=Mus musculus GN=Gas7 PE=1 SV=1\|Growth arrest-specific prot | TRUE | FALSE | NMDA - control | supernatant | fdrtool | 0,083664 | 0,599702 | FALSE | trend |
| GRIA2 | increased | P23819-4 | Isoform 4 of Glutamate receptor 2 OS=Mus musculus GN=Gria2 | FALSE | TRUE | NMDA - control | pellet | fdrtool | 0,060949 | 0,556062 | FALSE | trend |
| GRIA3 | increased | B0QZW1\|Q9Z2W9 | Glutamate receptor 3 OS=Mus musculus GN=Gria3 PE=1 SV=1\|Glutamate receptor 3 OS=Mus muscu | FALSE | TRUE | NMDA - control | pellet | fdrtool | 0,067273 | 0,570056 | FALSE | trend |
| GRIN1 | increased | A2AI14\|A2AI21 | Glutamate receptor ionotropic, NMDA 1 OS=Mus musculus GN=Grin1 PE=1 SV=1\|Glutamate recepto | FALSE | TRUE | NMDA - control | pellet | fdrtool | 0,069244 | 0,574021 | FALSE | trend |
| GSK3A | increased | D3Z7E5\|Q2NL51 | Glycogen synthase kinase-3 alpha OS=Mus musculus GN=Gsk3a PE=1 SV=1\|Glycogen synthase kinas | TRUE | FALSE | NMDA - control | supernatant | fdrtool | 0,011091 | 0,170516 | FALSE | candidate |
| H1F0 | increased | P10922 | Histone H1.0 OS=Mus musculus GN=H1f0 PE=2 SV=4 | FALSE | TRUE | NMDA - control | pellet | fdrtool | 0,008942 | 0,229121 | FALSE | trend |
| HNRNPA1 | increased | P49312\|P49312-2\|Q5EBP8 | Heterogeneous nuclear ribonucleoprotein A1 OS=Mus musculus GN=Hnrnpa1 PE=1 SV=2\|Isoform Sh | TRUE | TRUE | NMDA - control | supernatant | fdrtool | 0,015557 | 0,219579 | FALSE | trend |
| HSPA5 | increased | P20029 | 78 kDa glucose-regulated protein OS=Mus musculus GN=Hspa5 PE=1 SV=3 | TRUE | TRUE | NMDA - control | supernatant | fdrtool | 0,050827 | 0,476478 | FALSE | trend |
| HSPA9 | increased | P38647 | Stress-70 protein, mitochondrial OS=Mus musculus GN=Hspa9 PE=1 SV=3 | TRUE | TRUE | NMDA - control | pellet | fdrtool | 0,045684 | 0,511432 | FALSE | trend |
| HYOU1 | increased | Q9JKR6 | Hypoxia up-regulated protein 1 OS=Mus musculus GN=Hyou1 PE=1 SV=1 | TRUE | TRUE | NMDA - control | supernatant | fdrtool | 0,000685 | 0,022855 | FALSE | candidate |
| ICA1 | increased | D3Z118\|D3Z119\|D3Z376\|P97411\|P97411-2\|S4R217 | Islet cell autoantigen 1 OS=Mus musculus GN=Ica1 PE=1 SV=1\|Islet cell autoantigen 1 OS=Mus musc | TRUE | FALSE | NMDA - control | supernatant | fdrtool | 0,082691 | 0,596892 | FALSE | trend |
| IL1RAPL1 | increased | B1ASU0\|P59823 | Interleukin-1 receptor accessory protein-like 1 OS=Mus musculus GN=Il1rapl1 PE=1 SV=1\|Interleukin | FALSE | TRUE | NMDA - control | pellet | fdrtool | 0,021933 | 0,379678 | FALSE | trend |
| INA | increased | P46660 | Alpha-internexin OS=Mus musculus GN=Ina PE=1 SV=3 | TRUE | TRUE | NMDA - control | supernatant | fdrtool | 0,074019 | 0,569973 | FALSE | trend |
| IQSEC1 | increased | E9PUA3\|Q8R0S2\|Q8R0S2-2 | IQ motif and SEC7 domain-containing protein 1 OS=Mus musculus GN=Iqsec1 PE=1 SV=1\|IQ motif ar | TRUE | TRUE | NMDA - control | supernatant | fdrtool | 6,91E-07 | 7,03E-05 | TRUE | hit |
| ITPKA | increased | Q8R071 | Inositol-trisphosphate 3-kinase A OS=Mus musculus GN=Itpka PE=1 SV=1 | TRUE | TRUE | NMDA - control | supernatant | fdrtool | 6,06E-14 | 4,04E-11 | TRUE | hit |
| KALRN | increased | A0A0A0MQF1\|A2CG49\|A2CG49-2\|A2CG49-7\|A2CG49-8\|A2CG49-9\|F6QYT9 | Isoform 2 of Kalirin OS=Mus musculus GN=Kalrn\|Isoform 7 of Kalirin OS=Mus musculus GN=Kalrn\|Is | TRUE | TRUE | NMDA - control | supernatant | fdrtool | 0,0432 | 0,436165 | FALSE | trend |
| KCNA1 | increased | P16388 | Potassium voltage-gated channel subfamily A member 1 OS=Mus musculus GN=Kcna1 PE=1 SV=1 | FALSE | TRUE | NMDA - control | pellet | fdrtool | 0,038794 | 0,483889 | FALSE | trend |
| KCNQ2 | increased | A0A0G2JFQ2\|B7ZBV4\|B7ZBV6\|B7ZBV7\|B7ZBV8\|B7ZBV9\|B7ZBW1\|B7ZBW2\|Q9Z351\|Q9Z351-12\|Q9Z351-1 | Potassium voltage-gated channel subfamily KQT member 2 OS=Mus musculus GN=Kcnq2 PE=1 SV=1 | FALSE | TRUE | NMDA - control | pellet | fdrtool | 0,080066 | 0,593041 | FALSE | trend |
| KIAA0196 | increased | Q8C2E7 | WASH complex subunit strumpellin OS=Mus musculus GN=Kiaa0196 PE=1 SV=2 | TRUE | FALSE | NMDA - control | supernatant | fdrtool | 0,083217 | 0,598417 | FALSE | trend |
| KIF2A | increased | E0CZ72\|F8VQ42\|P28740\|P28740-1\|P28740-2 | Isoform 1 of Kinesin-like protein KIF2A OS=Mus musculus GN=Kif2a\|Isoform 2 of Kinesin-like protein | TRUE | TRUE | NMDA - control | supernatant | fdrtool | 3,11E-09 | 6,91E-07 | TRUE | hit |
| KIF5C | increased | P28738 | Kinesin heavy chain isoform 5C OS=Mus musculus GN=Kif5c PE=1 SV=3 | TRUE | TRUE | NMDA - control | supernatant | fdrtool | 0,065448 | 0,539587 | FALSE | trend |
| LGI1 | increased | A0A0G2JGB7\|A0A0G2JGE8\|Q9JIA1 | Leucine-rich glioma-inactivated protein 1 OS=Mus musculus GN=Lgi1 PE=1 SV=1\|Leucine-rich glioma | TRUE | FALSE | NMDA - control | supernatant | fdrtool | 0,006597 | 0,122188 | FALSE | candidate |
| MACROD1 | increased | Q922B1 | O-acetyl-ADP-ribose deacetylase MACROD1 OS=Mus musculus GN=Macrod1 PE=1 SV=2 | TRUE | FALSE | NMDA - control | supernatant | fdrtool | 0,07252 | 0,564951 | FALSE | trend |
| MADD | increased | A0A0R4J1F4\|A2AGQ2\|A2AGQ3\|A2AGQ4\|A2AGQ5\|A2AGQ6\|A2AGQ7\|A2AGQ8\|A2AGQ9\|A2AGR0\|A2AGR1\| | Isoform 10 of MAP kinase-activating death domain protein OS=Mus musculus GN=Madd\|Isoform 11 | TRUE | TRUE | NMDA - control | supernatant | fdrtool | 1,26E-07 | 1,69E-05 | TRUE | hit |
| MAP6 | increased | D3Z6W1\|Q7TSJ2\|Q7TSJ2-2 | Isoform 2 of Microtubule-associated protein 6 OS=Mus musculus GN=Map6\|Microtubule-associated | TRUE | TRUE | NMDA - control | supernatant | fdrtool | 0,012089 | 0,182066 | FALSE | candidate |
| MAPRE2 | increased | D3YYK8\|E9Q6X0\|Q8R001\|Q8R001-2 | Microtubule-associated protein RP/EB family member 2 (Fragment) OS=Mus musculus GN=Mapre2 | TRUE | TRUE | NMDA - control | supernatant | fdrtool | 0,000236 | 0,010615 | TRUE | hit |
| MARK1 | increased | Q8VHJ5 | Serine/threonine-protein kinase MARK1 OS=Mus musculus GN=Mark1 PE=1 SV=2 | TRUE | TRUE | NMDA - control | supernatant | fdrtool | 0,000255 | 0,011144 | TRUE | hit |
| MPP2 | increased | Q9WV34\|Q9WV34-2 | MAGUK p55 subfamily member 2 OS=Mus musculus GN=Mpp2 PE=1 SV=1\|Isoform 2 of MAGUK p55 | TRUE | TRUE | NMDA - control | supernatant | fdrtool | 0,07113 | 0,560189 | FALSE | trend |
| MPST | increased | Q3UW66\|Q99J99 | Sulfurtransferase OS=Mus musculus GN=Mpst PE=1 SV=1\|3-mercaptopyruvate sulfurtransferase OS= | TRUE | TRUE | NMDA - control | pellet | fdrtool | 0,051643 | 0,531071 | FALSE | trend |
| MRPL1 | increased | Q99N96\|Q9D3F3 | 39S ribosomal protein L1, mitochondrial OS=Mus musculus GN=Mrpl1 PE=1 SV=2\|39S ribosomal pro | TRUE | TRUE | NMDA - control | supernatant | fdrtool | 0,055816 | 0,499871 | FALSE | trend |

| | | | | | | | | | | | | |
|---|---|---|---|---|---|---|---|---|---|---|---|---|
| MYH10 | increased | Q3UH59\|Q5SV64\|Q61879 | Myosin-10 OS=Mus musculus GN=Myh10 PE=1 SV=1\|Myosin-10 OS=Mus musculus GN=Myh10 PE=1 | TRUE | TRUE | NMDA - control | supernatant | fdrtool | 0,033313 | 0,373641 | FALSE | trend |
| MYH9 | increased | Q8VDD5 | Myosin-9 OS=Mus musculus GN=Myh9 PE=1 SV=4 | TRUE | TRUE | NMDA - control | supernatant | fdrtool | 0,031414 | 0,360008 | FALSE | trend |
| MYL12B | increased | Q3THE2 | Myosin regulatory light chain 12B OS=Mus musculus GN=Myl12b PE=1 SV=2 | TRUE | TRUE | NMDA - control | supernatant | fdrtool | 0,018739 | 0,251242 | FALSE | trend |
| MYO5A | increased | D3YZ62\|D3Z4J3 | Unconventional myosin-Va OS=Mus musculus GN=Myo5a PE=1 SV=1\|Unconventional myosin-Va OS | TRUE | TRUE | NMDA - control | supernatant | fdrtool | 0,000143 | 0,007535 | TRUE | hit |
| NCEH1 | increased | Q8BLF1 | Neutral cholesterol ester hydrolase 1 OS=Mus musculus GN=Nceh1 PE=1 SV=1 | FALSE | TRUE | NMDA - control | pellet | fdrtool | 0,060385 | 0,554709 | FALSE | trend |
| NCKAP1 | increased | A2AS98\|P28660 | Nck-associated protein 1 OS=Mus musculus GN=Nckap1 PE=1 SV=1\|Nck-associated protein 1 OS=Mu | TRUE | TRUE | NMDA - control | supernatant | fdrtool | 0,008278 | 0,142424 | FALSE | candidate |
| NDUFA10 | increased | Q99LC3 | NADH dehydrogenase [ubiquinone] 1 alpha subcomplex subunit 10, mitochondrial OS=Mus musculu | TRUE | TRUE | NMDA - control | pellet | fdrtool | 0,024397 | 0,399675 | FALSE | trend |
| NDUFA5 | increased | Q9CPP6 | NADH dehydrogenase [ubiquinone] 1 alpha subcomplex subunit 5 OS=Mus musculus GN=Ndufa5 PE | TRUE | TRUE | NMDA - control | pellet | fdrtool | 0,05884 | 0,550904 | FALSE | trend |
| NDUFA8 | increased | Q9DCJ5 | NADH dehydrogenase [ubiquinone] 1 alpha subcomplex subunit 8 OS=Mus musculus GN=Ndufa8 PE | TRUE | TRUE | NMDA - control | pellet | fdrtool | 0,028984 | 0,431721 | FALSE | trend |
| NDUFS2 | increased | Q91WD5 | NADH dehydrogenase [ubiquinone] iron-sulfur protein 2, mitochondrial OS=Mus musculus GN=Nduf | TRUE | TRUE | NMDA - control | pellet | fdrtool | 0,042484 | 0,499379 | FALSE | trend |
| NDUFS3 | increased | Q9DCT2 | NADH dehydrogenase [ubiquinone] iron-sulfur protein 3, mitochondrial OS=Mus musculus GN=Nduf | TRUE | TRUE | NMDA - control | pellet | fdrtool | 0,012333 | 0,27427 | FALSE | trend |
| NDUFS5 | increased | B1ARW4\|Q99LY9 | NADH dehydrogenase [ubiquinone] iron-sulfur protein 5 (Fragment) OS=Mus musculus GN=Ndufs5 P | TRUE | TRUE | NMDA - control | pellet | fdrtool | 0,024136 | 0,397663 | FALSE | trend |
| NEFL | increased | P08551 | Neurofilament light polypeptide OS=Mus musculus GN=Nefl PE=1 SV=5 | TRUE | TRUE | NMDA - control | supernatant | fdrtool | 0,018923 | 0,253091 | FALSE | trend |
| NEFM | increased | A0A0R4J036\|P08553 | Neurofilament 3, medium OS=Mus musculus GN=Nefm PE=1 SV=1\|Neurofilament medium polypept | TRUE | TRUE | NMDA - control | supernatant | fdrtool | 0,007903 | 0,138166 | FALSE | candidate |
| NGEF | increased | E9QK62\|Q8CHT1\|Q8CHT1-2 | Ephexin-1 OS=Mus musculus GN=Ngef PE=1 SV=1\|Ephexin-1 OS=Mus musculus GN=Ngef PE=1 SV=1 | TRUE | TRUE | NMDA - control | supernatant | fdrtool | 0,00132 | 0,038371 | FALSE | candidate |
| NMT2 | increased | A2AJH3\|O70311\|Q3THR4\|Q8CFK1 | Glycylpeptide N-tetradecanoyltransferase OS=Mus musculus GN=Nmt2 PE=1 SV=1\|Glycylpeptide N- | TRUE | FALSE | NMDA - control | supernatant | fdrtool | 0,0002 | 0,009506 | FALSE | candidate |
| OLFM1 | increased | A3KGE4\|G3X9Q7\|O88998\|O88998-3\|Q8R357 | Noelin OS=Mus musculus GN=Olfm1 PE=1 SV=1\|Noelin OS=Mus musculus GN=Olfm1 PE=1 SV=1\|No | TRUE | TRUE | NMDA - control | supernatant | fdrtool | 0,082854 | 0,597364 | FALSE | trend |
| PACS1 | increased | Q8K212 | Phosphofurin acidic cluster sorting protein 1 OS=Mus musculus GN=Pacs1 PE=1 SV=2 | TRUE | TRUE | NMDA - control | supernatant | fdrtool | 0,001794 | 0,048085 | FALSE | candidate |
| PDE2A | increased | Q3TXZ6\|Q5PR72\|Q922S4 | Phosphodiesterase OS=Mus musculus GN=Pde2a PE=1 SV=1\|Phosphodiesterase OS=Mus musculus | TRUE | TRUE | NMDA - control | supernatant | fdrtool | 0,027696 | 0,331523 | FALSE | trend |
| PFKL | increased | P12382 | ATP-dependent 6-phosphofructokinase, liver type OS=Mus musculus GN=Pfkl PE=1 SV=4 | TRUE | TRUE | NMDA - control | pellet | fdrtool | 0,001932 | 0,110732 | FALSE | candidate |
| PHF24 | increased | Q80TL4\|Q80TL4-2 | Isoform 2 of PHD finger protein 24 OS=Mus musculus GN=Phf24\|PHD finger protein 24 OS=Mus mus | TRUE | TRUE | NMDA - control | supernatant | fdrtool | 0,035235 | 0,386855 | FALSE | trend |
| PICK1 | increased | E9QAC6 | Galectin OS=Mus musculus GN=Pick1 PE=1 SV=1 | TRUE | FALSE | NMDA - control | supernatant | fdrtool | 0,018995 | 0,253812 | FALSE | trend |
| PIP5K1C | increased | F8WHW6\|O70161\|O70161-2\|O70161-3 | Isoform 2 of Phosphatidylinositol 4-phosphate 5-kinase type-1 gamma OS=Mus musculus GN=Pip5k1 | TRUE | TRUE | NMDA - control | supernatant | fdrtool | 0,024203 | 0,30236 | FALSE | trend |
| PLCB1 | increased | F6X322\|Q9Z1B3\|Q9Z1B3-2\|Q9Z1B3-3 | Phosphoinositide phospholipase C OS=Mus musculus GN=Plcb1 PE=1 SV=2\|1-phosphatidylinositol 4 | TRUE | TRUE | NMDA - control | pellet | fdrtool | 0,020729 | 0,369055 | FALSE | trend |
| PLEC | increased | E9Q3W4\|Q9QXS1\|Q9QXS1-10\|Q9QXS1-11\|Q9QXS1-12\|Q9QXS1-13\|Q9QXS1-14\|Q9QXS1-15\|Q9QXS1-16\|C | Plectin OS=Mus musculus GN=Plec PE=1 SV=1\|Plectin OS=Mus musculus GN=Plec PE=1 SV=3\|Isofor | TRUE | TRUE | NMDA - control | supernatant | fdrtool | 7,58E-07 | 7,54E-05 | TRUE | hit |
| POMGNT2 | increased | Q8BW41\|Q8BW41-2 | Protein O-linked-mannose beta-1,4-N-acetylglucosaminyltransferase 2 OS=Mus musculus GN=Pomgr | FALSE | TRUE | NMDA - control | pellet | fdrtool | 0,000177 | 0,035835 | TRUE | hit |
| PPP1R9A | increased | H3BJD0\|H3BJD6\|H3BKQ7\|H3BL28\|Q7TN74 | Neurabin I OS=Mus musculus GN=Ppp1r9a PE=1 SV=1\|Protein Ppp1r9a (Fragment) OS=Mus musculu | TRUE | TRUE | NMDA - control | supernatant | fdrtool | 8,55E-05 | 0,005162 | TRUE | hit |
| PPP1R9B | increased | Q6R891 | Neurabin-2 OS=Mus musculus GN=Ppp1r9b PE=1 SV=1 | TRUE | TRUE | NMDA - control | pellet | fdrtool | 0,034914 | 0,465424 | FALSE | trend |
| PRKAR1A | increased | Q9DBC7 | cAMP-dependent protein kinase type I-alpha regulatory subunit OS=Mus musculus GN=Prkar1a PE= | TRUE | FALSE | NMDA - control | supernatant | fdrtool | 0,051486 | 0,479695 | FALSE | trend |
| PRKCA | increased | Q4VA93 | Protein kinase C OS=Mus musculus GN=Prkca PE=1 SV=1 | TRUE | FALSE | NMDA - control | supernatant | fdrtool | 0,035664 | 0,389734 | FALSE | trend |
| PRKCB | increased | P68404\|P68404-2 | Isoform Beta-II of Protein kinase C beta type OS=Mus musculus GN=Prkcb\|Protein kinase C beta typ | TRUE | TRUE | NMDA - control | pellet | fdrtool | 0,0027 | 0,135199 | FALSE | candidate |
| PRKCB | increased | P68404\|P68404-2 | Isoform Beta-II of Protein kinase C beta type OS=Mus musculus GN=Prkcb\|Protein kinase C beta typ | TRUE | TRUE | NMDA - control | supernatant | fdrtool | 0,076131 | 0,576855 | FALSE | trend |
| PRKCE | increased | P16054 | Protein kinase C epsilon type OS=Mus musculus GN=Prkce PE=1 SV=1 | TRUE | TRUE | NMDA - control | supernatant | fdrtool | 1,26E-07 | 1,68E-05 | TRUE | hit |
| PRKCG | increased | P63318 | Protein kinase C gamma type OS=Mus musculus GN=Prkcg PE=1 SV=1 | TRUE | TRUE | NMDA - control | pellet | fdrtool | 0,000766 | 0,068779 | FALSE | candidate |
| PSD3 | increased | E9PUC5\|F6Z9E6\|Q2PFD7\|Q2PFD7-2\|Q2PFD7-3\|Q2PFD7-5\|Q8C0E9 | Isoform 2 of PH and SEC7 domain-containing protein 3 OS=Mus musculus GN=Psd3\|Isoform 3 of PH | TRUE | TRUE | NMDA - control | supernatant | fdrtool | 0,000436 | 0,016611 | TRUE | hit |
| PSMC2 | increased | P46471\|Q8BVQ9 | 26S protease regulatory subunit 7 OS=Mus musculus GN=Psmc2 PE=1 SV=5\|26S protease regulatory | TRUE | TRUE | NMDA - control | supernatant | fdrtool | 0,049841 | 0,471597 | FALSE | trend |
| PSMC5 | increased | P62196\|Q8K1K2 | 26S protease regulatory subunit 8 OS=Mus musculus GN=Psmc5 PE=1 SV=1 | TRUE | TRUE | NMDA - control | supernatant | fdrtool | 0,037552 | 0,402066 | FALSE | trend |
| RAB14 | increased | Q50HX4\|Q91V41 | RAB14 protein OS=Mus musculus GN=Rab14 PE=1 SV=1\|Ras-related protein Rab-14 OS=Mus muscul | TRUE | TRUE | NMDA - control | supernatant | fdrtool | 0,039408 | 0,413719 | FALSE | trend |
| RAB1A | increased | Q0PD67\|Q5SW88 | Protein Rab1a OS=Mus musculus GN=Rab1a PE=1 SV=1\|RAB1, member RAS oncogene family, isoforr | TRUE | TRUE | NMDA - control | supernatant | fdrtool | 0,00438 | 0,09024 | FALSE | candidate |
| RAB1B | increased | Q0PD66\|Q9D1G1 | RAB1B, member RAS oncogene family, isoform CRA_c OS=Mus musculus GN=Rab1b PE=1 SV=1\|Ras- | TRUE | TRUE | NMDA - control | pellet | fdrtool | 0,079376 | 0,59195 | FALSE | trend |
| RAB2A | increased | P53994\|Q0PD65 | Ras-related protein Rab-2A OS=Mus musculus GN=Rab2a PE=1 SV=1\|RAB2, member RAS oncogene f | TRUE | TRUE | NMDA - control | supernatant | fdrtool | 0,001819 | 0,048559 | FALSE | candidate |
| RAB3A | increased | P63011\|Q0PD63 | Ras-related protein Rab-3A OS=Mus musculus GN=Rab3a PE=1 SV=1\|RAB3A, member RAS oncogene | TRUE | TRUE | NMDA - control | supernatant | fdrtool | 0,000176 | 0,008711 | FALSE | candidate |
| RAB3C | increased | P62823\|Q542T7 | Ras-related protein Rab-3C OS=Mus musculus GN=Rab3c PE=1 SV=1\|RAB3C, member RAS oncogene | TRUE | TRUE | NMDA - control | supernatant | fdrtool | 1,04E-09 | 2,79E-07 | TRUE | hit |
| RAP2A | increased | Q80ZJ1 | Ras-related protein Rap-2a OS=Mus musculus GN=Rap2a PE=1 SV=2 | FALSE | TRUE | NMDA - control | pellet | fdrtool | 0,082194 | 0,596315 | FALSE | trend |
| RAP2B | increased | P61226\|Q6ZWR0 | Ras-related protein Rap-2b OS=Mus musculus GN=Rap2b PE=1 SV=1\|MCG5466 OS=Mus musculus G | FALSE | TRUE | NMDA - control | pellet | fdrtool | 0,00105 | 0,078213 | FALSE | candidate |
| RASAL1 | increased | Q9Z268 | RasGAP-activating-like protein 1 OS=Mus musculus GN=Rasal1 PE=1 SV=2 | TRUE | FALSE | NMDA - control | supernatant | fdrtool | 0,076598 | 0,578346 | FALSE | trend |
| RGS7 | increased | A0A0A6YW36\|O54829\|Q45FE1\|Q45FE2\|Q80XD3 | Regulator of G-protein-signaling 7 OS=Mus musculus GN=Rgs7 PE=1 SV=1\|Regulator of G-protein sig | TRUE | TRUE | NMDA - control | supernatant | fdrtool | 0,00312 | 0,069253 | FALSE | candidate |
| RHEB | increased | Q540E6\|Q921J2 | GTP-binding protein Rheb OS=Mus musculus GN=Rheb PE=1 SV=1\|GTP-binding protein Rheb OS=Mu | TRUE | TRUE | NMDA - control | pellet | fdrtool | 0,065476 | 0,566288 | FALSE | trend |
| ROCK2 | increased | F8VPK5\|P70336\|P70336-2 | Rho-associated protein kinase OS=Mus musculus GN=Rock2 PE=1 SV=1\|Rho-associated protein kina | TRUE | TRUE | NMDA - control | supernatant | fdrtool | 0,011075 | 0,170372 | FALSE | candidate |
| RPH3A | increased | P47708 | Rabphilin-3A OS=Mus musculus GN=Rph3a PE=1 SV=2 | TRUE | TRUE | NMDA - control | supernatant | fdrtool | 0,001338 | 0,038766 | TRUE | hit |
| RPL30 | increased | P62889\|Q58DZ3 | 60S ribosomal protein L30 OS=Mus musculus GN=Rpl30 PE=1 SV=2\|60S ribosomal protein L30 OS=M | TRUE | TRUE | NMDA - control | pellet | fdrtool | 0,048731 | 0,521888 | FALSE | trend |
| RPS10 | increased | P63325\|Q5M9K7 | 40S ribosomal protein S10 OS=Mus musculus GN=Rps10 PE=1 SV=1\|40S ribosomal protein S10 OS=M | FALSE | TRUE | NMDA - control | pellet | fdrtool | 0,037434 | 0,477695 | FALSE | trend |
| RPS6KA2 | increased | Q9WUT3 | Ribosomal protein S6 kinase alpha-2 OS=Mus musculus GN=Rps6ka2 PE=1 SV=1 | TRUE | FALSE | NMDA - control | supernatant | fdrtool | 0,008236 | 0,141955 | FALSE | candidate |
| RTN1 | increased | Q4FJL2\|Q7M6W1\|Q8K0T0 | Reticulon-1 OS=Mus musculus GN=Rtn1 PE=1 SV=1\|Reticulon OS=Mus musculus GN=Rtn1 PE=1 SV= | TRUE | TRUE | NMDA - control | supernatant | fdrtool | 2,24E-06 | 0,0002 | TRUE | hit |
| RTN3 | increased | Q9ES97\|Q9ES97-2\|Q9ES97-3 | Isoform 2 of Reticulon-3 OS=Mus musculus GN=Rtn3\|Isoform 3 of Reticulon-3 OS=Mus musculus GN | TRUE | TRUE | NMDA - control | supernatant | fdrtool | 0,001946 | 0,050926 | FALSE | candidate |
| SCN1B | increased | P97952 | Sodium channel subunit beta-1 OS=Mus musculus GN=Scn1b PE=1 SV=1 | FALSE | TRUE | NMDA - control | pellet | fdrtool | 0,026871 | 0,417713 | FALSE | trend |
| SCN2A1 | increased | A0A0J9YTW6\|B1AWN6 | Protein Scn2a1 OS=Mus musculus GN=Scn2a1 PE=1 SV=1\|Protein Scn2a1 OS=Mus musculus GN=Scn | FALSE | TRUE | NMDA - control | pellet | fdrtool | 0,05795 | 0,548647 | FALSE | trend |
| SCN3B | increased | H3BJ27\|H3BJR6\|Q0P666\|Q8BHK2 | Sodium channel subunit beta-3 (Fragment) OS=Mus musculus GN=Scn3b PE=1 SV=1\|Sodium channe | FALSE | TRUE | NMDA - control | pellet | fdrtool | 0,009327 | 0,232173 | FALSE | trend |
| SGIP1 | increased | F7CPX0\|Q8VD37\|Q8VD37-2\|Q8VD37-3\|Q8VD37-4\|Q8VD37-6\|Q8VD37-8 | Isoform 2 of SH3-containing GRB2-like protein 3-interacting protein 1 OS=Mus musculus GN=Sgip1\|I | TRUE | TRUE | NMDA - control | supernatant | fdrtool | 0,04314 | 0,435823 | FALSE | trend |
| SIK3 | increased | E9PU87\|F6S7W6\|F6U6U5\|F6U8X4\|Q6P4S6\|Q6P4S6-2 | Isoform 2 of Serine/threonine-protein kinase SIK3 OS=Mus musculus GN=Sik3\|Serine/threonine-pro | TRUE | TRUE | NMDA - control | supernatant | fdrtool | 0,065356 | 0,539235 | FALSE | trend |
| SLC12A5 | increased | Q91V14\|Q91V14-2 | Isoform 2 of Solute carrier family 12 member 5 OS=Mus musculus GN=Slc12a5\|Solute carrier family | TRUE | TRUE | NMDA - control | supernatant | fdrtool | 0,047493 | 0,45959 | FALSE | trend |
| SLC17A7 | increased | Q3TXX4 | Vesicular glutamate transporter 1 OS=Mus musculus GN=Slc17a7 PE=1 SV=2 | FALSE | TRUE | NMDA - control | pellet | fdrtool | 0,004357 | 0,172785 | FALSE | candidate |
| SLC2A3 | increased | P32037\|Q3TPL8 | Solute carrier family 2, facilitated glucose transporter member 3 OS=Mus musculus GN=Slc2a3 PE=1 | FALSE | TRUE | NMDA - control | pellet | fdrtool | 0,053336 | 0,53608 | FALSE | trend |
| SLC30A3 | increased | P97441\|Q3TMQ7\|S4R169 | Zinc transporter 3 OS=Mus musculus GN=Slc30a3 PE=1 SV=1\|Zinc transporter 3 OS=Mus musculus G | FALSE | TRUE | NMDA - control | pellet | fdrtool | 0,065083 | 0,565443 | FALSE | trend |
| SLC8A2 | increased | Q8K596\|Q8K596-2 | Sodium/calcium exchanger 2 OS=Mus musculus GN=Slc8a2 PE=1 SV=1\|Isoform 2 of Sodium/calcium | FALSE | TRUE | NMDA - control | pellet | fdrtool | 0,042153 | 0,498063 | FALSE | trend |
| SNPH | increased | Q80U23\|Q80U23-2\|Q80U23-3 | Isoform 2 of Syntaphilin OS=Mus musculus GN=Snph\|Isoform 3 of Syntaphilin OS=Mus musculus GN | TRUE | TRUE | NMDA - control | supernatant | fdrtool | 0,017512 | 0,238718 | FALSE | trend |
| SPTBN2 | increased | Q68FG2 | Protein Sptbn2 OS=Mus musculus GN=Sptbn2 PE=1 SV=1 | TRUE | TRUE | NMDA - control | supernatant | fdrtool | 0,025291 | 0,311714 | FALSE | trend |
| SRCIN1 | increased | B1AQX6\|B1AQX7\|B1AQX9\|Q9QWI6\|Q9QWI6-2\|Q9QWI6-3 | Isoform 2 of SRC kinase signaling inhibitor 1 OS=Mus musculus GN=Srcin1\|Isoform 3 of SRC kinase si | TRUE | TRUE | NMDA - control | supernatant | fdrtool | 9,76E-05 | 0,005665 | TRUE | hit |
| STX1A | increased | O35526\|Q497P1 | Syntaxin-1A OS=Mus musculus GN=Stx1a PE=1 SV=3\|Syntaxin 1A (Brain) OS=Mus musculus GN=Stx1 | FALSE | TRUE | NMDA - control | pellet | fdrtool | 0,014307 | 0,300354 | FALSE | trend |
| STX1B | increased | P61264 | Syntaxin-1B OS=Mus musculus GN=Stx1b PE=1 SV=1 | TRUE | TRUE | NMDA - control | supernatant | fdrtool | 5,57E-09 | 1,03E-06 | TRUE | hit |
| STXBP1 | increased | O08599\|O08599-2 | Syntaxin-binding protein 1 OS=Mus musculus GN=Stxbp1 PE=1 SV=2\|Isoform 2 of Syntaxin-binding p | TRUE | TRUE | NMDA - control | supernatant | fdrtool | 0,000483 | 0,017886 | FALSE | candidate |
| SUB1 | increased | P11031 | Activated RNA polymerase II transcriptional coactivator p15 OS=Mus musculus GN=Sub1 PE=1 SV=3 | FALSE | TRUE | NMDA - control | pellet | fdrtool | 0,012086 | 0,271131 | FALSE | trend |
| SV2B | increased | Q8BG39 | Synaptic vesicle glycoprotein 2B OS=Mus musculus GN=Sv2b PE=1 SV=1 | FALSE | TRUE | NMDA - control | pellet | fdrtool | 0,013814 | 0,294055 | FALSE | trend |
| SYN1 | increased | O88935 | Synapsin-1 OS=Mus musculus GN=Syn1 PE=1 SV=2 | TRUE | TRUE | NMDA - control | supernatant | fdrtool | 0,011152 | 0,171055 | FALSE | candidate |
| SYN2 | increased | Q64332\|Q64332-2 | Isoform IIb of Synapsin-2 OS=Mus musculus GN=Syn2\|Synapsin-2 OS=Mus musculus GN=Syn2 PE=1 | TRUE | TRUE | NMDA - control | supernatant | fdrtool | 0,035653 | 0,389656 | FALSE | trend |
| SYNGAP1 | increased | A0A0A6YVS6\|A0A0J9YVH8\|A0A140T8K9\|F6SEU4\|J3QQ18 | Ras/Rap GTPase-activating protein SynGAP (Fragment) OS=Mus musculus GN=Syngap1 PE=1 SV=2\|R | TRUE | TRUE | NMDA - control | supernatant | fdrtool | 0,00012 | 0,006642 | FALSE | candidate |
| SYNGR3 | increased | Q8R191 | Synaptogyrin-3 OS=Mus musculus GN=Syngr3 PE=1 SV=1 | TRUE | TRUE | NMDA - control | supernatant | fdrtool | 0,019566 | 0,259456 | FALSE | trend |
| SYP | increased | Q62277 | Synaptophysin OS=Mus musculus GN=Syp PE=1 SV=2 | TRUE | TRUE | NMDA - control | supernatant | fdrtool | 0,003116 | 0,069196 | FALSE | candidate |
| SYT1 | increased | H6RXZ1\|P46096 | Synaptotagmin I OS=Mus musculus GN=Syt1 PE=1 SV=1\|Synaptotagmin-1 OS=Mus musculus GN=Sy | TRUE | TRUE | NMDA - control | supernatant | fdrtool | 2,71E-11 | 9,05E-09 | TRUE | hit |
| TARS | increased | Q9D0R2 | Threonine--tRNA ligase, cytoplasmic OS=Mus musculus GN=Tars PE=1 SV=2 | TRUE | FALSE | NMDA - control | supernatant | fdrtool | 0,016452 | 0,228511 | FALSE | trend |
| TIMM44 | increased | O35857 | Mitochondrial import inner membrane translocase subunit TIM44 OS=Mus musculus GN=Timm44 PI | TRUE | TRUE | NMDA - control | supernatant | fdrtool | 0,040467 | 0,420168 | FALSE | trend |
| TMEM109 | increased | Q3UBX0 | Transmembrane protein 109 OS=Mus musculus GN=Tmem109 PE=1 SV=2 | FALSE | TRUE | NMDA - control | pellet | fdrtool | 0,077559 | 0,589005 | FALSE | trend |
| TMEM35 | increased | Q9D328 | Transmembrane protein 35 OS=Mus musculus GN=Tmem35 PE=1 SV=1 | FALSE | TRUE | NMDA - control | pellet | fdrtool | 0,045226 | 0,509776 | FALSE | trend |
| TNIK | increased | B2RQ80\|B9EKN8\|E0CXD6\|E0CY98\|E0CZD7\|E0CZF8\|E9PUL9\|P83510\|P83510-2 | Isoform 2 of Traf2 and NCK-interacting protein kinase OS=Mus musculus GN=Tnik\|Tnik protein OS=M | TRUE | TRUE | NMDA - control | supernatant | fdrtool | 0,000287 | 0,011987 | TRUE | hit |
| TOMM70 | increased | Q9CZW5 | Mitochondrial import receptor subunit TOM70 OS=Mus musculus GN=Tomm70 PE=1 SV=2 | TRUE | TRUE | NMDA - control | supernatant | fdrtool | 0,000164 | 0,008289 | FALSE | candidate |
| TPPP | increased | Q3URG1\|Q7TQD2 | RIKEN cDNA 2900041A09, isoform CRA_a OS=Mus musculus GN=Tppp PE=1 SV=1\|Tubulin polymeriz | TRUE | TRUE | NMDA - control | supernatant | fdrtool | 0,005053 | 0,100678 | FALSE | candidate |
| TRIM2 | increased | E9QKC6\|Q3UHH4\|Q9ESN6 | Tripartite motif-containing protein 2 OS=Mus musculus GN=Trim2 PE=1 SV=1\|Tripartite motif-contai | TRUE | TRUE | NMDA - control | supernatant | fdrtool | 0,00035 | 0,014035 | FALSE | candidate |
| TRIO | increased | Q0KL02\|Q0KL02-2\|Q0KL02-3\|Q0KL02-4 | Isoform 2 of Triple functional domain protein OS=Mus musculus GN=Trio\|Isoform 3 of Triple functio | TRUE | TRUE | NMDA - control | supernatant | fdrtool | 0,000844 | 0,026812 | FALSE | candidate |
| TUBA4A | increased | A0A0A0MQA5\|P68368 | Tubulin alpha-4A chain (Fragment) OS=Mus musculus GN=Tuba4a PE=1 SV=1\|Tubulin alpha-4A chair | TRUE | TRUE | NMDA - control | pellet | fdrtool | 0,006135 | 0,199982 | FALSE | candidate |
| TUFM | increased | Q8BFR5 | Elongation factor Tu, mitochondrial OS=Mus musculus GN=Tufm PE=1 SV=1 | TRUE | TRUE | NMDA - control | pellet | fdrtool | 0,000154 | 0,03506 | TRUE | hit |
| VSNL1 | increased | P62761\|Q4W4C9 | Visinin-like protein 1 OS=Mus musculus GN=Vsnl1 PE=1 SV=2\|Neural visinin-like type 1 protein OS=N | TRUE | TRUE | NMDA - control | pellet | fdrtool | 0,064726 | 0,564668 | FALSE | trend |
| WDR37 | increased | H3BJB3\|Q8CBE3 | WD repeat-containing protein 37 (Fragment) OS=Mus musculus GN=Wdr37 PE=1 SV=7\|WD repeat-c | TRUE | TRUE | NMDA - control | supernatant | fdrtool | 0,001556 | 0,043397 | FALSE | candidate |
| 2010300C02RIK | decreased | E9Q3M9 | Protein 2010300C02Rik OS=Mus musculus GN=2010300C02Rik PE=1 SV=1 | TRUE | TRUE | NMDA - control | supernatant | fdrtool | 0,049336 | 0,469057 | FALSE | trend |
| AAK1 | decreased | Q3UHJ0\|Q3UHJ0-2 | AP2-associated protein kinase 1 OS=Mus musculus GN=Aak1 PE=1 SV=2\|Isoform 2 of AP2-associatec | TRUE | TRUE | NMDA - control | pellet | fdrtool | 0,014549 | 0,303388 | FALSE | trend |
| ACADM | decreased | P45952 | Medium-chain specific acyl-CoA dehydrogenase, mitochondrial OS=Mus musculus GN=Acadm PE=1 S | TRUE | FALSE | NMDA - control | supernatant | fdrtool | 0,017907 | 0,242802 | FALSE | trend |
| ACADS | decreased | Q07417 | Short-chain specific acyl-CoA dehydrogenase, mitochondrial OS=Mus musculus GN=Acads PE=1 SV=2 | TRUE | FALSE | NMDA - control | supernatant | fdrtool | 0,035474 | 0,38846 | FALSE | trend |
| ACADVL | decreased | B1AR28\|P50544 | Very long-chain-specific acyl-CoA dehydrogenase, mitochondrial OS=Mus musculus GN=Acadvl PE=1 | TRUE | TRUE | NMDA - control | supernatant | fdrtool | 0,043396 | 0,437279 | FALSE | trend |
| ACOT7 | decreased | Q91V12-2 | Isoform A of Cytosolic acyl coenzyme A thioester hydrolase OS=Mus musculus GN=Acot7 | TRUE | FALSE | NMDA - control | supernatant | fdrtool | 0,076504 | 0,578047 | FALSE | trend |
| ACOX1 | decreased | Q9R0H0\|Q9R0H0-2 | Peroxisomal acyl-coenzyme A oxidase 1 OS=Mus musculus GN=Acox1 PE=1 SV=5\|Isoform 2 of Perox | TRUE | FALSE | NMDA - control | supernatant | fdrtool | 0,036568 | 0,395699 | FALSE | trend |
| ACSBG1 | decreased | Q99PU5 | Long-chain-fatty-acid--CoA ligase ACSBG1 OS=Mus musculus GN=Acsbg1 PE=1 SV=1 | TRUE | TRUE | NMDA - control | supernatant | fdrtool | 0,022028 | 0,282866 | FALSE | trend |
| ACSF2 | decreased | Q8VCW8 | Acyl-CoA synthetase family member 2, mitochondrial OS=Mus musculus GN=Acsf2 PE=1 SV=1 | TRUE | FALSE | NMDA - control | supernatant | fdrtool | 0,000787 | 0,025437 | FALSE | candidate |

| | | | | | | | | | | | | |
|---|---|---|---|---|---|---|---|---|---|---|---|---|
| ADD1 | decreased | F8WGR0\|F8WHZ9\|Q9QYC0\|Q9QYC0-2 | Alpha-adducin OS=Mus musculus GN=Add1 PE=1 SV=1\|Alpha-adducin OS=Mus musculus GN=Add1 | TRUE | TRUE | NMDA - control | pellet | fdrtool | 0,006434 | 0,203819 | FALSE | trend |
| ADD2 | decreased | Q9QYB8\|Q9QYB8-2 | Beta-adducin OS=Mus musculus GN=Add2 PE=1 SV=4\|Isoform 2 of Beta-adducin OS=Mus musculus | TRUE | TRUE | NMDA - control | supernatant | fdrtool | 4,90E-05 | 0,003365 | FALSE | candidate |
| AHCYL1 | decreased | Q80SW1 | Putative adenosylhomocysteinase 2 OS=Mus musculus GN=Ahcyl1 PE=1 SV=1 | TRUE | TRUE | NMDA - control | pellet | fdrtool | 0,001155 | 0,08173 | FALSE | candidate |
| AHCYL2 | decreased | F8WGT1\|Q68FL4 | Adenosylhomocysteinase OS=Mus musculus GN=Ahcyl2 PE=1 SV=1\|Putative adenosylhomocysteina | TRUE | TRUE | NMDA - control | pellet | fdrtool | 0,000618 | 0,063523 | FALSE | candidate |
| AK3 | decreased | Q9WTP7 | GTP:AMP phosphotransferase AK3, mitochondrial OS=Mus musculus GN=Ak3 PE=1 SV=3 | TRUE | TRUE | NMDA - control | pellet | fdrtool | 0,066082 | 0,567575 | FALSE | trend |
| ALDH1A1 | decreased | P24549 | Retinal dehydrogenase 1 OS=Mus musculus GN=Aldh1a1 PE=1 SV=5 | TRUE | FALSE | NMDA - control | supernatant | fdrtool | 0,055919 | 0,500331 | FALSE | trend |
| ALDH4A1 | decreased | Q8CHT0 | Delta-1-pyrroline-5-carboxylate dehydrogenase, mitochondrial OS=Mus musculus GN=Aldh4a1 PE=1 | TRUE | FALSE | NMDA - control | supernatant | fdrtool | 0,077566 | 0,581405 | FALSE | trend |
| ALDH6A1 | decreased | Q9EQ20 | Methylmalonate-semialdehyde dehydrogenase [acylating], mitochondrial OS=Mus musculus GN=Ald | TRUE | TRUE | NMDA - control | supernatant | fdrtool | 0,080711 | 0,591048 | FALSE | trend |
| ALDOC | decreased | P05063 | Fructose-bisphosphate aldolase C OS=Mus musculus GN=Aldoc PE=1 SV=4 | TRUE | TRUE | NMDA - control | supernatant | fdrtool | 0,046947 | 0,45672 | FALSE | trend |
| AMPH | decreased | A0A0G2JEG8\|Q7TQF7 | Amphiphysin OS=Mus musculus GN=Amph PE=1 SV=1\|Amphiphysin OS=Mus musculus GN=Amph P | TRUE | TRUE | NMDA - control | pellet | fdrtool | 0,005196 | 0,186686 | FALSE | candidate |
| ANKFY1 | decreased | Q810B6 | Rabankyrin-5 OS=Mus musculus GN=Ankfy1 PE=1 SV=2 | TRUE | FALSE | NMDA - control | supernatant | fdrtool | 0,052926 | 0,486582 | FALSE | trend |
| AP3B2 | decreased | Q9JME5 | AP-3 complex subunit beta-2 OS=Mus musculus GN=Ap3b2 PE=1 SV=2 | TRUE | TRUE | NMDA - control | pellet | fdrtool | 0,008031 | 0,221133 | FALSE | trend |
| ARHGAP32 | decreased | Q811P8 | Rho GTPase-activating protein 32 OS=Mus musculus GN=Arhgap32 PE=1 SV=2 | FALSE | TRUE | NMDA - control | pellet | fdrtool | 0,072599 | 0,580386 | FALSE | trend |
| ARHGEF11 | decreased | E9Q0A3\|Q68FM7 | Protein Arhgef11 (Fragment) OS=Mus musculus GN=Arhgef11 PE=1 SV=7\|Protein Arhgef11 OS=Mus | FALSE | TRUE | NMDA - control | pellet | fdrtool | 0,078438 | 0,590443 | FALSE | trend |
| ARHGEF7 | decreased | A0A0R4J0C5\|A0A0R4J0X8\|D3Z0V2\|Q9ES28\|Q9ES28-2\|Q9ES28-3\|Q9ES28-4\|Q9ES28-5\|Q9ES28-6\|Q9ES28-7 | Rho guanine nucleotide exchange factor 7 OS=Mus musculus GN=Arhgef7 PE=1 SV=1\|Rho guanine n | TRUE | TRUE | NMDA - control | pellet | fdrtool | 0,004711 | 0,178979 | FALSE | candidate |
| ASAH1 | decreased | Q78P93\|Q9WV54 | Acid ceramidase OS=Mus musculus GN=Asah1 PE=1 SV=1\|Acid ceramidase OS=Mus musculus GN=A | TRUE | FALSE | NMDA - control | supernatant | fdrtool | 6,27E-05 | 0,004093 | TRUE | hit |
| ASRGL1 | decreased | Q8C0M9 | Isoaspartyl peptidase/L-asparaginase OS=Mus musculus GN=Asrgl1 PE=1 SV=1 | TRUE | FALSE | NMDA - control | supernatant | fdrtool | 0,068259 | 0,550013 | FALSE | trend |
| ATP5E | decreased | P56382\|Q545F5 | ATP synthase subunit epsilon, mitochondrial OS=Mus musculus GN=Atp5e PE=1 SV=2\|ATP synthase | TRUE | TRUE | NMDA - control | pellet | fdrtool | 0,066946 | 0,569383 | FALSE | trend |
| ATP6V1G2 | decreased | B0V2H4\|Q54A87\|Q9WTT4 | ATPase, H+ transporting, lysosomal V1 subunit G2, isoform CRA_a OS=Mus musculus GN=Atp6v1g2 | TRUE | TRUE | NMDA - control | pellet | fdrtool | 0,000413 | 0,053105 | FALSE | candidate |
| ATPIF1 | decreased | E9PV44\|O35143 | ATPase inhibitor, mitochondrial OS=Mus musculus GN=Atpif1 PE=1 SV=1\|ATPase inhibitor, mitochor | TRUE | TRUE | NMDA - control | supernatant | fdrtool | 0,022263 | 0,285025 | FALSE | trend |
| ATXN10 | decreased | P28658 | Ataxin-10 OS=Mus musculus GN=Atxn10 PE=1 SV=2 | TRUE | FALSE | NMDA - control | supernatant | fdrtool | 0,041988 | 0,429178 | FALSE | trend |
| BCL2L13 | decreased | P59017 | Bcl-2-like protein 13 OS=Mus musculus GN=Bcl2l13 PE=1 SV=2 | FALSE | TRUE | NMDA - control | pellet | fdrtool | 0,051609 | 0,530968 | FALSE | trend |
| BIN1 | decreased | O08539 | Myc box-dependent-interacting protein 1 OS=Mus musculus GN=Bin1 PE=1 SV=1 | TRUE | TRUE | NMDA - control | pellet | fdrtool | 0,003524 | 0,155781 | FALSE | candidate |
| BRK1 | decreased | Q91VR8 | Protein BRICK1 OS=Mus musculus GN=Brk1 PE=1 SV=1 | FALSE | TRUE | NMDA - control | pellet | fdrtool | 0,010701 | 0,252424 | FALSE | trend |
| CALB2 | decreased | Q08331 | Calretinin OS=Mus musculus GN=Calb2 PE=1 SV=3 | TRUE | FALSE | NMDA - control | supernatant | fdrtool | 0,004795 | 0,096758 | FALSE | candidate |
| CAMSAP2 | decreased | A0A0A6YXV8\|A0A0A6YY67\|H7BX08\|Q8C1B1\|Q8C1B1-2 | Calmodulin-regulated spectrin-associated protein 2 (Fragment) OS=Mus musculus GN=Camsap2 PE= | FALSE | TRUE | NMDA - control | pellet | fdrtool | 0,076762 | 0,587679 | FALSE | trend |
| CAT | decreased | P24270 | Catalase OS=Mus musculus GN=Cat PE=1 SV=4 | TRUE | FALSE | NMDA - control | supernatant | fdrtool | 0,080431 | 0,590206 | FALSE | trend |
| CBLN1 | decreased | Q7TNF5\|Q9R171 | Cbln1 protein OS=Mus musculus GN=Cbln1 PE=1 SV=2\|Cerebellin-1 OS=Mus musculus GN=Cbln1 PE | TRUE | FALSE | NMDA - control | supernatant | fdrtool | 1,95E-07 | 2,36E-05 | TRUE | hit |
| CBS | decreased | Q91WT9\|Q91WT9-2 | Cystathionine beta-synthase OS=Mus musculus GN=Cbs PE=1 SV=3\|Isoform 2 of Cystathionine beta- | TRUE | FALSE | NMDA - control | supernatant | fdrtool | 0,02132 | 0,276291 | FALSE | trend |
| CCT7 | decreased | A0A0N4SV00\|P80313\|Q3TIJ7 | Chaperonin subunit 7 (Eta), isoform CRA_a OS=Mus musculus GN=Cct7 PE=1 SV=1\|T-complex protei | TRUE | TRUE | NMDA - control | pellet | fdrtool | 0,005564 | 0,191994 | FALSE | candidate |
| CDC42BPB | decreased | Q7TT50 | Serine/threonine-protein kinase MRCK beta OS=Mus musculus GN=Cdc42bpb PE=1 SV=2 | FALSE | TRUE | NMDA - control | pellet | fdrtool | 0,080923 | 0,594376 | FALSE | trend |
| CEP170 | decreased | H7BX26\|Q6A065 | Centrosomal protein of 170 kDa OS=Mus musculus GN=Cep170 PE=1 SV=1\|Centrosomal protein of 1 | FALSE | TRUE | NMDA - control | pellet | fdrtool | 0,059505 | 0,552561 | FALSE | trend |
| CFL1 | decreased | P18760 | Cofilin-1 OS=Mus musculus GN=Cfl1 PE=1 SV=3 | TRUE | TRUE | NMDA - control | pellet | fdrtool | 0,007215 | 0,212889 | FALSE | trend |
| CFL2 | decreased | P45591\|Q3UHW9 | Cofilin-2 OS=Mus musculus GN=Cfl2 PE=1 SV=1\|Cofilin-2 OS=Mus musculus GN=Cfl2 PE=1 SV=1 | TRUE | FALSE | NMDA - control | supernatant | fdrtool | 0,013928 | 0,20257 | FALSE | trend |
| CLASP1 | decreased | A0A087WNW8\|A0A087WP42\|A0A087WQ31\|A0A087WQ95\|A0A087WQF1\|A0A087WRE8\|A0A087WRK0\|A | CLIP-associating protein 1 OS=Mus musculus GN=Clasp1 PE=1 SV=2\|CLIP-associating protein 1 OS= | FALSE | TRUE | NMDA - control | pellet | fdrtool | 0,01079 | 0,253676 | FALSE | trend |
| CNBP | decreased | A0A0N4SVS6\|P53996\|P53996-2\|P53996-3\|Q3ULK8 | Cellular nucleic acid-binding protein OS=Mus musculus GN=Cnbp PE=1 SV=1\|Cellular nucleic acid-bi | FALSE | TRUE | NMDA - control | pellet | fdrtool | 0,000212 | 0,037438 | TRUE | hit |
| CORO1B | decreased | A2RS22\|Q9WUM3 | Coronin OS=Mus musculus GN=Coro1b PE=1 SV=1\|Coronin-1B OS=Mus musculus GN=Coro1b PE=1 | FALSE | TRUE | NMDA - control | pellet | fdrtool | 0,0015 | 0,096106 | FALSE | candidate |
| CORO1C | decreased | Q499X7\|Q9WUM4 | Coronin OS=Mus musculus GN=Coro1c PE=1 SV=1\|Coronin-1C OS=Mus musculus GN=Coro1c PE=1 | TRUE | TRUE | NMDA - control | pellet | fdrtool | 0,000858 | 0,071419 | FALSE | candidate |
| CPLX1 | decreased | P63040 | Complexin-1 OS=Mus musculus GN=Cplx1 PE=1 SV=1 | TRUE | TRUE | NMDA - control | pellet | fdrtool | 0,004438 | 0,174246 | FALSE | candidate |
| CPLX1 | decreased | P63040 | Complexin-1 OS=Mus musculus GN=Cplx1 PE=1 SV=1 | TRUE | TRUE | NMDA - control | supernatant | fdrtool | 4,07E-06 | 0,000339 | TRUE | hit |
| CPLX2 | decreased | P84086 | Complexin-2 OS=Mus musculus GN=Cplx2 PE=1 SV=1 | TRUE | TRUE | NMDA - control | supernatant | fdrtool | 0,007415 | 0,13241 | FALSE | candidate |
| CRMP1 | decreased | P97427\|Q6P1J1 | Crmp1 protein OS=Mus musculus GN=Crmp1 PE=1 SV=1\|Dihydropyrimidinase-related protein 1 OS= | TRUE | TRUE | NMDA - control | pellet | fdrtool | 0,005394 | 0,189593 | FALSE | candidate |
| CSRP1 | decreased | P97315\|Q4FJX4 | Cysteine and glycine-rich protein 1 OS=Mus musculus GN=Csrp1 PE=1 SV=3\|Csrp1 protein OS=Mus | TRUE | FALSE | NMDA - control | supernatant | fdrtool | 0,017455 | 0,238182 | FALSE | trend |
| CTNNA2 | decreased | E0CXB9\|Q61301\|Q61301-2\|Q61301-3 | Catenin alpha-2 OS=Mus musculus GN=Ctnna2 PE=1 SV=1\|Catenin alpha-2 OS=Mus musculus GN=C | TRUE | TRUE | NMDA - control | supernatant | fdrtool | 0,05746 | 0,507127 | FALSE | trend |
| CTNNB1 | decreased | Q02248 | Catenin beta-1 OS=Mus musculus GN=Ctnnb1 PE=1 SV=1 | TRUE | TRUE | NMDA - control | pellet | fdrtool | 0,060211 | 0,554288 | FALSE | trend |
| CTSD | decreased | P18242\|Q3UCD9 | Cathepsin D OS=Mus musculus GN=Ctsd PE=1 SV=1\|Cathepsin D OS=Mus musculus GN=Ctsd PE=1 S | TRUE | FALSE | NMDA - control | supernatant | fdrtool | 0,003699 | 0,078946 | FALSE | candidate |
| CTTN | decreased | Q60598 | Src substrate cortactin OS=Mus musculus GN=Cttn PE=1 SV=2 | TRUE | FALSE | NMDA - control | supernatant | fdrtool | 0,010525 | 0,165353 | FALSE | candidate |
| DBI | decreased | P31786\|Q4VWZ5\|Q548W7 | Acyl-CoA-binding protein OS=Mus musculus GN=Dbi PE=1 SV=2\|Acyl-CoA-binding protein OS=Mus | TRUE | FALSE | NMDA - control | supernatant | fdrtool | 0,053884 | 0,491065 | FALSE | trend |
| DECR1 | decreased | Q4FJK0\|Q9CQ62 | 2,4-dienoyl-CoA reductase, mitochondrial OS=Mus musculus GN=Decr1 PE=1 SV=1\|2,4-dienoyl-CoA | TRUE | TRUE | NMDA - control | supernatant | fdrtool | 0,04522 | 0,447435 | FALSE | trend |
| DLG2 | decreased | D3YUZ8\|Q91XM9\|Q91XM9-2\|Q91XM9-3\|Q91XM9-4\|Q91XM9-6\|Q91XM9-7\|Q91XM9-8 | Disks large homolog 2 OS=Mus musculus GN=Dlg2 PE=1 SV=1\|Disks large homolog 2 OS=Mus muscu | TRUE | TRUE | NMDA - control | pellet | fdrtool | 0,008702 | 0,227124 | FALSE | trend |
| DMTN | decreased | Q9WV69\|Q9WV69-2\|Q9WV69-3 | Dematin OS=Mus musculus GN=Dmtn PE=1 SV=1\|Isoform 2 of Dematin OS=Mus musculus GN=Dmtr | TRUE | TRUE | NMDA - control | pellet | fdrtool | 0,004888 | 0,181887 | FALSE | candidate |
| DNAJB4 | decreased | Q9D832 | DnaJ homolog subfamily B member 4 OS=Mus musculus GN=Dnajb4 PE=1 SV=1 | FALSE | TRUE | NMDA - control | pellet | fdrtool | 0,038894 | 0,484337 | FALSE | trend |
| DNM1 | decreased | A0A0J9YUN4\|P39053-3\|P39053-4\|P39053-5\|P39053-6 | Dynamin-1 OS=Mus musculus GN=Dnm1 PE=1 SV=1\|Isoform 3 of Dynamin-1 OS=Mus musculus GN= | TRUE | TRUE | NMDA - control | pellet | fdrtool | 0,031859 | 0,448995 | FALSE | trend |
| DNM2 | decreased | G3X9G4\|Q3TCR7 | Dynamin-2 OS=Mus musculus GN=Dnm2 PE=1 SV=1\|Dynamin-2 OS=Mus musculus GN=Dnm2 PE=1 | TRUE | TRUE | NMDA - control | pellet | fdrtool | 0,031803 | 0,448675 | FALSE | trend |
| DPYSL2 | decreased | O08553 | Dihydropyrimidinase-related protein 2 OS=Mus musculus GN=Dpysl2 PE=1 SV=2 | TRUE | TRUE | NMDA - control | pellet | fdrtool | 0,000119 | 0,033467 | TRUE | hit |
| DPYSL3 | decreased | Q62188 | Dihydropyrimidinase-related protein 3 OS=Mus musculus GN=Dpysl3 PE=1 SV=1 | TRUE | TRUE | NMDA - control | pellet | fdrtool | 0,002565 | 0,131288 | FALSE | candidate |
| DPYSL4 | decreased | O35098\|Q3TMU8 | Dihydropyrimidinase-related protein 4 OS=Mus musculus GN=Dpysl4 PE=1 SV=1\|Dihydropyrimidina | TRUE | TRUE | NMDA - control | pellet | fdrtool | 0,007179 | 0,2125 | FALSE | trend |
| DPYSL5 | decreased | Q9EQF6 | Dihydropyrimidinase-related protein 5 OS=Mus musculus GN=Dpysl5 PE=1 SV=1 | TRUE | TRUE | NMDA - control | pellet | fdrtool | 0,000743 | 0,068056 | FALSE | candidate |
| DSTN | decreased | Q4FK36\|Q9R0P5 | Destrin OS=Mus musculus GN=Dstn PE=1 SV=1\|Destrin OS=Mus musculus GN=Dstn PE=1 SV=3 | TRUE | FALSE | NMDA - control | supernatant | fdrtool | 0,040366 | 0,419556 | FALSE | trend |
| ECH1 | decreased | O35459 | Delta(3,5)-Delta(2,4)-dienoyl-CoA isomerase, mitochondrial OS=Mus musculus GN=Ech1 PE=1 SV=1 | TRUE | FALSE | NMDA - control | supernatant | fdrtool | 0,049004 | 0,467377 | FALSE | trend |
| ECI1 | decreased | P42125 | Enoyl-CoA delta isomerase 1, mitochondrial OS=Mus musculus GN=Eci1 PE=1 SV=2 | TRUE | FALSE | NMDA - control | supernatant | fdrtool | 0,063868 | 0,533507 | FALSE | trend |
| EEF1A1 | decreased | P10126\|Q58E64 | Elongation factor 1-alpha 1 OS=Mus musculus GN=Eef1a1 PE=1 SV=3\|Elongation factor 1-alpha OS= | TRUE | TRUE | NMDA - control | pellet | fdrtool | 0,012506 | 0,276439 | FALSE | trend |
| EEF1G | decreased | Q4FZK2\|Q9D8N0 | Elongation factor 1-gamma OS=Mus musculus GN=Eef1g PE=1 SV=1\|Elongation factor 1-gamma OS= | TRUE | TRUE | NMDA - control | pellet | fdrtool | 0,020937 | 0,370932 | FALSE | trend |
| EFHD2 | decreased | Q8C845\|Q9D8Y0 | EF-hand domain-containing protein D2 OS=Mus musculus GN=Efhd2 PE=1 SV=1\|EF-hand domain-co | TRUE | TRUE | NMDA - control | pellet | fdrtool | 0,040634 | 0,491843 | FALSE | trend |
| EIF2S2 | decreased | Q3ULL5\|Q99L45 | Eif2s2 protein OS=Mus musculus GN=Eif2s2 PE=1 SV=1\|Eukaryotic translation initiation factor 2 sub | FALSE | TRUE | NMDA - control | pellet | fdrtool | 0,072594 | 0,580375 | FALSE | trend |
| EIF3A | decreased | P23116 | Eukaryotic translation initiation factor 3 subunit A OS=Mus musculus GN=Eif3a PE=1 SV=5 | FALSE | TRUE | NMDA - control | pellet | fdrtool | 0,07818 | 0,590024 | FALSE | trend |
| EIF4B | decreased | Q8BGD9 | Eukaryotic translation initiation factor 4B OS=Mus musculus GN=Eif4b PE=1 SV=1 | FALSE | TRUE | NMDA - control | pellet | fdrtool | 0,000664 | 0,065302 | FALSE | candidate |
| EIF4H | decreased | Q9WUK2 | Eukaryotic translation initiation factor 4H OS=Mus musculus GN=Eif4h PE=1 SV=3 | FALSE | TRUE | NMDA - control | pellet | fdrtool | 0,028935 | 0,431409 | FALSE | trend |
| EML2 | decreased | E9QK48 | Echinoderm microtubule-associated protein-like 2 OS=Mus musculus GN=Eml2 PE=1 SV=1 | FALSE | TRUE | NMDA - control | pellet | fdrtool | 0,023721 | 0,394406 | FALSE | trend |
| EPB41L1 | decreased | A2AUK5\|A2AUK8 | Band 4.1-like protein 1 OS=Mus musculus GN=Epb41l1 PE=1 SV=1 | TRUE | TRUE | NMDA - control | pellet | fdrtool | 0,015025 | 0,30923 | FALSE | trend |
| EPB41L3 | decreased | Q9WV92\|Q9WV92-2\|Q9WV92-3\|Q9WV92-8 | Band 4.1-like protein 3 OS=Mus musculus GN=Epb41l3 PE=1 SV=1\|Isoform 2 of Band 4.1-like protein | TRUE | TRUE | NMDA - control | pellet | fdrtool | 0,043135 | 0,501926 | FALSE | trend |
| ETFB | decreased | Q9DCW4 | Electron transfer flavoprotein subunit beta OS=Mus musculus GN=Etfb PE=1 SV=3 | TRUE | TRUE | NMDA - control | pellet | fdrtool | 0,005893 | 0,196696 | FALSE | candidate |
| FABP5 | decreased | Q05816\|Q497I3 | Fatty acid-binding protein, epidermal OS=Mus musculus GN=Fabp5 PE=1 SV=3\|Fatty acid binding pr | TRUE | FALSE | NMDA - control | supernatant | fdrtool | 0,002581 | 0,061577 | FALSE | candidate |
| FABP7 | decreased | E9Q0H6\|P51880 | Fatty acid-binding protein, brain OS=Mus musculus GN=Fabp7 PE=1 SV=1\|Fatty acid-binding protein | TRUE | TRUE | NMDA - control | supernatant | fdrtool | 6,91E-05 | 0,004411 | TRUE | hit |
| FABP7 | decreased | E9Q0H6\|P51880 | Fatty acid-binding protein, brain OS=Mus musculus GN=Fabp7 PE=1 SV=1\|Fatty acid-binding protein | TRUE | TRUE | NMDA - control | pellet | fdrtool | 0,018634 | 0,349055 | FALSE | trend |
| FAM73B | decreased | Q8BK03\|Q8BK03-2 | Mitoguardin-2 OS=Mus musculus GN=Fam73b PE=1 SV=1\|Isoform 2 of Mitoguardin-2 OS=Mus musc | FALSE | TRUE | NMDA - control | pellet | fdrtool | 0,032115 | 0,450444 | FALSE | trend |
| FDXR | decreased | Q61578 | NADPH:adrenodoxin oxidoreductase, mitochondrial OS=Mus musculus GN=Fdxr PE=1 SV=1 | TRUE | TRUE | NMDA - control | pellet | fdrtool | 0,012108 | 0,271407 | FALSE | trend |
| FERMT2 | decreased | Q3TLE2\|Q8CIB5 | Fermitin family homolog 2 OS=Mus musculus GN=Fermt2 PE=1 SV=1\|Fermitin family homolog 2 OS= | TRUE | FALSE | NMDA - control | supernatant | fdrtool | 0,070754 | 0,558884 | FALSE | trend |
| GAD1 | decreased | P48318\|Q548L6 | Glutamate decarboxylase 1 OS=Mus musculus GN=Gad1 PE=1 SV=2\|Glutamate decarboxylase OS=M | TRUE | TRUE | NMDA - control | supernatant | fdrtool | 0,010342 | 0,163631 | FALSE | candidate |
| GFAP | decreased | P03995 | Glial fibrillary acidic protein OS=Mus musculus GN=Gfap PE=1 SV=4 | TRUE | TRUE | NMDA - control | supernatant | fdrtool | 0,00151 | 0,042462 | FALSE | candidate |
| GFER | decreased | P56213 | FAD-linked sulfhydryl oxidase ALR OS=Mus musculus GN=Gfer PE=1 SV=2 | FALSE | TRUE | NMDA - control | pellet | fdrtool | 0,049129 | 0,523188 | FALSE | trend |
| GLO1 | decreased | A5GZX3\|Q9CPU0 | Lactoylglutathione lyase OS=Mus musculus GN=Glo1 PE=1 SV=1\|Lactoylglutathione lyase OS=Mus | TRUE | FALSE | NMDA - control | supernatant | fdrtool | 0,082267 | 0,595653 | FALSE | trend |
| GLUD1 | decreased | P26443 | Glutamate dehydrogenase 1, mitochondrial OS=Mus musculus GN=Glud1 PE=1 SV=1 | TRUE | TRUE | NMDA - control | supernatant | fdrtool | 0,009998 | 0,160339 | FALSE | candidate |
| GLUL | decreased | P15105 | Glutamine synthetase OS=Mus musculus GN=Glul PE=1 SV=6 | TRUE | TRUE | NMDA - control | supernatant | fdrtool | 0,020932 | 0,272635 | FALSE | trend |
| GLUL | decreased | P15105 | Glutamine synthetase OS=Mus musculus GN=Glul PE=1 SV=6 | TRUE | TRUE | NMDA - control | pellet | fdrtool | 0,068741 | 0,573026 | FALSE | trend |
| GPHN | decreased | A0JNY3\|Q8BUV3 | Gephyrin OS=Mus musculus GN=Gphn PE=1 SV=1\|Gephyrin OS=Mus musculus GN=Gphn PE=1 SV=2 | TRUE | TRUE | NMDA - control | pellet | fdrtool | 0,001985 | 0,112308 | FALSE | candidate |
| GPRIN1 | decreased | Q3UNH4 | G protein-regulated inducer of neurite outgrowth 1 OS=Mus musculus GN=Gprin1 PE=1 SV=2 | TRUE | TRUE | NMDA - control | pellet | fdrtool | 0,003899 | 0,163889 | FALSE | candidate |
| GSTK1 | decreased | Q9DCM2 | Glutathione S-transferase kappa 1 OS=Mus musculus GN=Gstk1 PE=1 SV=3 | TRUE | FALSE | NMDA - control | supernatant | fdrtool | 0,080831 | 0,591405 | FALSE | trend |
| GSTM5 | decreased | P48774 | Glutathione S-transferase Mu 5 OS=Mus musculus GN=Gstm5 PE=1 SV=1 | TRUE | FALSE | NMDA - control | supernatant | fdrtool | 0,010095 | 0,161275 | FALSE | candidate |
| HADH | decreased | Q61425 | Hydroxyacyl-coenzyme A dehydrogenase, mitochondrial OS=Mus musculus GN=Hadh PE=1 SV=2 | TRUE | FALSE | NMDA - control | supernatant | fdrtool | 0,002748 | 0,064066 | FALSE | candidate |
| HADHB | decreased | Q99JY0 | Trifunctional enzyme subunit beta, mitochondrial OS=Mus musculus GN=Hadhb PE=1 SV=1 | TRUE | TRUE | NMDA - control | pellet | fdrtool | 0,053261 | 0,535862 | FALSE | trend |
| HBA | decreased | P01942\|Q91VB8 | Hemoglobin subunit alpha OS=Mus musculus GN=Hba PE=1 SV=2\|Alpha globin 1 OS=Mus musculus | TRUE | FALSE | NMDA - control | supernatant | fdrtool | 0,075119 | 0,573585 | FALSE | trend |
| HDHD2 | decreased | Q3UGR5 | Haloacid dehalogenase-like hydrolase domain-containing protein 2 OS=Mus musculus GN=Hdhd2 PE | TRUE | TRUE | NMDA - control | pellet | fdrtool | 0,027836 | 0,424259 | FALSE | trend |
| HMGCS1 | decreased | Q8JZK9 | Hydroxymethylglutaryl-CoA synthase, cytoplasmic OS=Mus musculus GN=Hmgcs1 PE=1 SV=1 | TRUE | FALSE | NMDA - control | supernatant | fdrtool | 0,044983 | 0,446138 | FALSE | trend |
| HSD11B1 | decreased | P50172\|Q3TJI8\|Q4JHD9 | Corticosteroid 11-beta-dehydrogenase isozyme 1 OS=Mus musculus GN=Hsd11b1 PE=1 SV=3\|Cortic | FALSE | TRUE | NMDA - control | pellet | fdrtool | 0,080997 | 0,594491 | FALSE | trend |
| HSDL2 | decreased | Q2TPA8 | Hydroxysteroid dehydrogenase-like protein 2 OS=Mus musculus GN=Hsdl2 PE=1 SV=1 | TRUE | FALSE | NMDA - control | supernatant | fdrtool | 0,021241 | 0,275552 | FALSE | trend |
| HSP90AB1 | decreased | P11499\|Q71LX8 | Heat shock protein HSP 90-beta OS=Mus musculus GN=Hsp90ab1 PE=1 SV=3\|Heat shock protein 84 | TRUE | TRUE | NMDA - control | pellet | fdrtool | 0,002946 | 0,141886 | FALSE | candidate |
| HSPA8 | decreased | P63017 | Heat shock cognate 71 kDa protein OS=Mus musculus GN=Hspa8 PE=1 SV=1 | TRUE | TRUE | NMDA - control | pellet | fdrtool | 0,018137 | 0,344002 | FALSE | trend |
| HUWE1 | decreased | A2AFQ0\|F6XP90\|Q7TMY8\|Q7TMY8-2\|Q7TMY8-3\|Q7TMY8-4 | E3 ubiquitin-protein ligase HUWE1 OS=Mus musculus GN=Huwe1 PE=1 SV=1\|E3 ubiquitin-protein li | FALSE | TRUE | NMDA - control | pellet | fdrtool | 0,06796 | 0,571458 | FALSE | trend |
| ICT1 | decreased | A2A6T4\|Q8R035\|Q8R035-2 | Immature colon carcinoma transcript 1, isoform CRA_c OS=Mus musculus GN=Ict1 PE=1 SV=1\|Isofor | TRUE | TRUE | NMDA - control | pellet | fdrtool | 0,052403 | 0,533347 | FALSE | trend |
| IDH2 | decreased | P54071 | Isocitrate dehydrogenase [NADP], mitochondrial OS=Mus musculus GN=Idh2 PE=1 SV=3 | TRUE | TRUE | NMDA - control | supernatant | fdrtool | 0,073755 | 0,569096 | FALSE | trend |

| | | | | | | | | | | | | |
|---|---|---|---|---|---|---|---|---|---|---|---|---|
| IDI1 | decreased | G3XA48\|P58044 | Isopentenyl-diphosphate Delta-isomerase 1 OS=Mus musculus GN=Idi1 PE=1 SV=1\|Isopentenyl-diph | TRUE | FALSE | NMDA - control | supernatant | fdrtool | 0,043651 | 0,43872 | FALSE | trend |
| INPP4A | decreased | A0A0R4J0Q2\|D3YUD3\|D3YUL8\|D3Z230\|E9Q9A0\|E9QAD0\|F6R4N2\|F6V2U0\|Q3UEQ1\|Q9EPW0\|Q9EPW0-2 | Type I inositol 3,4-bisphosphate 4-phosphatase OS=Mus musculus GN=Inpp4a PE=1 SV=1\|Type I ino | TRUE | TRUE | NMDA - control | pellet | fdrtool | 0,050559 | 0,527738 | FALSE | trend |
| IRGQ | decreased | Q8VIM9 | Immunity-related GTPase family Q protein OS=Mus musculus GN=Irgq PE=1 SV=1 | TRUE | FALSE | NMDA - control | supernatant | fdrtool | 0,01241 | 0,185749 | FALSE | candidate |
| KARS | decreased | Q3TIV6\|Q8R2P8\|Q99MN1 | Lysine--tRNA ligase OS=Mus musculus GN=Kars PE=1 SV=1\|Lysine--tRNA ligase OS=Mus musculus GN | TRUE | TRUE | NMDA - control | pellet | fdrtool | 0,082191 | 0,59631 | FALSE | trend |
| KLC1 | decreased | E9Q7C9\|Q5UE59\|Q7TNF4\|Q8CD76 | Kinesin light chain 1 OS=Mus musculus GN=Klc1 PE=1 SV=1\|Kinesin light chain 1 OS=Mus musculus ( | TRUE | TRUE | NMDA - control | pellet | fdrtool | 0,016228 | 0,323343 | FALSE | trend |
| LAMTOR2 | decreased | Q9JHS3 | Ragulator complex protein LAMTOR2 OS=Mus musculus GN=Lamtor2 PE=1 SV=1 | TRUE | TRUE | NMDA - control | supernatant | fdrtool | 0,010979 | 0,169514 | FALSE | candidate |
| LAMTOR2 | decreased | Q9JHS3 | Ragulator complex protein LAMTOR2 OS=Mus musculus GN=Lamtor2 PE=1 SV=1 | TRUE | TRUE | NMDA - control | pellet | fdrtool | 0,075882 | 0,586188 | FALSE | trend |
| MAP1A | decreased | A2ARP8\|Q9QYR6\|Q9QYR6-2 | Microtubule-associated protein 1A OS=Mus musculus GN=Map1a PE=1 SV=1\|Microtubule-associate | TRUE | TRUE | NMDA - control | pellet | fdrtool | 0,00031 | 0,046302 | TRUE | hit |
| MAP1B | decreased | P14873 | Microtubule-associated protein 1B OS=Mus musculus GN=Map1b PE=1 SV=2 | TRUE | TRUE | NMDA - control | pellet | fdrtool | 0,000825 | 0,070499 | FALSE | candidate |
| MAP2 | decreased | G3UZJ2\|P20357 | Microtubule-associated protein (Fragment) OS=Mus musculus GN=Map2 PE=1 SV=7\|Microtubule-as | TRUE | TRUE | NMDA - control | pellet | fdrtool | 3,14E-08 | 4,44E-05 | TRUE | hit |
| MAP4 | decreased | A0A0B4J1F2\|A0A0B4J1M9\|E9PZ43\|P27546\|P27546-2\|P27546-3 | Microtubule-associated protein OS=Mus musculus GN=Map4 PE=1 SV=1\|Microtubule-associated pr | FALSE | TRUE | NMDA - control | pellet | fdrtool | 0,025588 | 0,408599 | FALSE | trend |
| MAPK8IP3 | decreased | E9Q6B6\|E9Q6E0\|J3QNR6\|K3W4S4\|Q9ESN9\|Q9ESN9-2\|Q9ESN9-3\|Q9ESN9-4\|Q9ESN9-5\|Q9ESN9-6 | C-Jun-amino-terminal kinase-interacting protein 3 OS=Mus musculus GN=Mapk8ip3 PE=1 SV=1\|C-Ju | FALSE | TRUE | NMDA - control | pellet | fdrtool | 0,06966 | 0,574835 | FALSE | trend |
| MAPRE2 | decreased | D3YYK8\|E9Q6X0\|Q8R001\|Q8R001-2 | Microtubule-associated protein RP/EB family member 2 (Fragment) OS=Mus musculus GN=Mapre2 F | TRUE | TRUE | NMDA - control | pellet | fdrtool | 0,009182 | 0,231043 | FALSE | trend |
| MAPT | decreased | A0A0A0MQC7\|A2A5Y6\|P10637\|P10637-2\|P10637-4\|P10637-5\|P10637-6 | Isoform Tau-A of Microtubule-associated protein tau OS=Mus musculus GN=Mapt\|Isoform Tau-C of | TRUE | TRUE | NMDA - control | pellet | fdrtool | 0,034487 | 0,463235 | FALSE | trend |
| MARCKS | decreased | P26645 | Myristoylated alanine-rich C-kinase substrate OS=Mus musculus GN=Marcks PE=1 SV=2 | FALSE | TRUE | NMDA - control | pellet | fdrtool | 0,016201 | 0,323041 | FALSE | trend |
| MARCKSL1 | decreased | P28667 | MARCKS-related protein OS=Mus musculus GN=Marcksl1 PE=1 SV=2 | FALSE | TRUE | NMDA - control | pellet | fdrtool | 0,000115 | 0,033266 | TRUE | hit |
| MCCC2 | decreased | B2RUK5\|Q3ULD5 | Methylcrotonoyl-CoA carboxylase beta chain, mitochondrial OS=Mus musculus GN=Mccc2 PE=1 SV= | TRUE | TRUE | NMDA - control | pellet | fdrtool | 0,047232 | 0,51686 | FALSE | trend |
| MRPL12 | decreased | Q9DB15 | 39S ribosomal protein L12, mitochondrial OS=Mus musculus GN=Mrpl12 PE=1 SV=2 | FALSE | TRUE | NMDA - control | pellet | fdrtool | 0,001011 | 0,076948 | FALSE | candidate |
| MRPL45 | decreased | Q9D0Q7 | 39S ribosomal protein L45, mitochondrial OS=Mus musculus GN=Mrpl45 PE=1 SV=1 | TRUE | TRUE | NMDA - control | pellet | fdrtool | 0,030533 | 0,441266 | FALSE | trend |
| MTFR1L | decreased | D3YW54\|D3Z0X8\|D6RCX5\|Q9CWE0 | Mitochondrial fission regulator 1-like (Fragment) OS=Mus musculus GN=Mtfr1l PE=1 SV=1\|Mitochor | TRUE | TRUE | NMDA - control | supernatant | fdrtool | 0,014964 | 0,213504 | FALSE | trend |
| MUT | decreased | P16332 | Methylmalonyl-CoA mutase, mitochondrial OS=Mus musculus GN=Mut PE=1 SV=2 | TRUE | TRUE | NMDA - control | pellet | fdrtool | 0,069258 | 0,574047 | FALSE | trend |
| NAP1L4 | decreased | A2RSB1\|B7ZNL2\|Q78ZA7 | Nucleosome assembly protein 1-like 4 OS=Mus musculus GN=Nap1l4 PE=1 SV=1\|Nap1l4 protein OS= | TRUE | TRUE | NMDA - control | pellet | fdrtool | 0,063992 | 0,563055 | FALSE | trend |
| NARS | decreased | Q8BP47 | Asparagine--tRNA ligase, cytoplasmic OS=Mus musculus GN=Nars PE=1 SV=2 | TRUE | TRUE | NMDA - control | pellet | fdrtool | 0,034333 | 0,462438 | FALSE | trend |
| NDUFA8 | decreased | Q9DCJ5 | NADH dehydrogenase [ubiquinone] 1 alpha subcomplex subunit 8 OS=Mus musculus GN=Ndufa8 PE | TRUE | TRUE | NMDA - control | supernatant | fdrtool | 0,079202 | 0,586477 | FALSE | trend |
| NEBL | decreased | Q9DC07 | LIM zinc-binding domain-containing Nebulette OS=Mus musculus GN=Nebl PE=1 SV=1 | TRUE | FALSE | NMDA - control | supernatant | fdrtool | 0,007127 | 0,128894 | FALSE | candidate |
| NF1 | decreased | Q04690\|Q04690-2\|Q04690-3\|Q04690-4 | Isoform 1 of Neurofibromin OS=Mus musculus GN=Nf1\|Isoform 3 of Neurofibromin OS=Mus muscul | TRUE | TRUE | NMDA - control | pellet | fdrtool | 0,035898 | 0,470342 | FALSE | trend |
| NFS1 | decreased | Q9Z1J3\|Q9Z1J3-2 | Cysteine desulfurase, mitochondrial OS=Mus musculus GN=Nfs1 PE=1 SV=3\|Isoform Cytoplasmic of | TRUE | TRUE | NMDA - control | pellet | fdrtool | 0,022764 | 0,386667 | FALSE | trend |
| NLGN3 | decreased | A2AGI2\|Q8BYM5 | Neuroligin-3 OS=Mus musculus GN=Nlgn3 PE=1 SV=1\|Neuroligin-3 OS=Mus musculus GN=Nlgn3 PE= | FALSE | TRUE | NMDA - control | pellet | fdrtool | 0,020774 | 0,369461 | FALSE | trend |
| NPC2 | decreased | Q9Z0J0 | Epididymal secretory protein E1 OS=Mus musculus GN=Npc2 PE=1 SV=1 | TRUE | FALSE | NMDA - control | supernatant | fdrtool | 0,037031 | 0,398713 | FALSE | trend |
| NRGN | decreased | P60761 | Neurogranin OS=Mus musculus GN=Nrgn PE=1 SV=1 | TRUE | TRUE | NMDA - control | pellet | fdrtool | 0,00697 | 0,210176 | FALSE | trend |
| OLA1 | decreased | Q9CZ30 | Obg-like ATPase 1 OS=Mus musculus GN=Ola1 PE=1 SV=1 | TRUE | TRUE | NMDA - control | pellet | fdrtool | 0,050435 | 0,527353 | FALSE | trend |
| OXCT1 | decreased | Q9D0K2 | Succinyl-CoA:3-ketoacid coenzyme A transferase 1, mitochondrial OS=Mus musculus GN=Oxct1 PE=1 | TRUE | TRUE | NMDA - control | pellet | fdrtool | 0,076622 | 0,587444 | FALSE | trend |
| PABPC1 | decreased | P29341 | Polyadenylate-binding protein 1 OS=Mus musculus GN=Pabpc1 PE=1 SV=2 | TRUE | TRUE | NMDA - control | pellet | fdrtool | 0,003311 | 0,150939 | FALSE | candidate |
| PACS1 | decreased | Q8K212 | Phosphofurin acidic cluster sorting protein 1 OS=Mus musculus GN=Pacs1 PE=1 SV=2 | TRUE | TRUE | NMDA - control | pellet | fdrtool | 0,080398 | 0,59356 | FALSE | trend |
| PAK1 | decreased | G5E884 | Serine/threonine-protein kinase PAK 1 OS=Mus musculus GN=Pak1 PE=1 SV=1 | TRUE | TRUE | NMDA - control | pellet | fdrtool | 0,073651 | 0,582288 | FALSE | trend |
| PCCB | decreased | Q99MN9 | Propionyl-CoA carboxylase beta chain, mitochondrial OS=Mus musculus GN=Pccb PE=1 SV=2 | TRUE | TRUE | NMDA - control | pellet | fdrtool | 0,072486 | 0,580177 | FALSE | trend |
| PCSK1N | decreased | Q9QXV0 | ProSAAS OS=Mus musculus GN=Pcsk1n PE=1 SV=2 | FALSE | TRUE | NMDA - control | pellet | fdrtool | 0,001572 | 0,098771 | FALSE | candidate |
| PEA15 | decreased | Q62048 | Astrocytic phosphoprotein PEA-15 OS=Mus musculus GN=Pea15 PE=1 SV=1 | TRUE | FALSE | NMDA - control | supernatant | fdrtool | 0,067081 | 0,545704 | FALSE | trend |
| PITPNM1 | decreased | O35954 | Membrane-associated phosphatidylinositol transfer protein 1 OS=Mus musculus GN=Pitpnm1 PE=1 S | TRUE | TRUE | NMDA - control | pellet | fdrtool | 0,065013 | 0,565291 | FALSE | trend |
| PKM | decreased | P52480\|P52480-2 | Pyruvate kinase PKM OS=Mus musculus GN=Pkm PE=1 SV=4\|Isoform M1 of Pyruvate kinase PKM OS | TRUE | TRUE | NMDA - control | pellet | fdrtool | 0,007481 | 0,215698 | FALSE | trend |
| PLCB1 | decreased | F6X322\|Q9Z1B3\|Q9Z1B3-2\|Q9Z1B3-3 | Phosphoinositide phospholipase C OS=Mus musculus GN=Plcb1 PE=1 SV=2\|1-phosphatidylinositol 4 | TRUE | TRUE | NMDA - control | supernatant | fdrtool | 0,041897 | 0,428648 | FALSE | trend |
| PLEC | decreased | E9Q3W4\|Q9QXS1\|Q9QXS1-10\|Q9QXS1-11\|Q9QXS1-12\|Q9QXS1-13\|Q9QXS1-14\|Q9QXS1-15\|Q9QXS1-16\|C | Plectin OS=Mus musculus GN=Plec PE=1 SV=1\|Plectin OS=Mus musculus GN=Plec PE=1 SV=3\|Isofor | TRUE | TRUE | NMDA - control | pellet | fdrtool | 0,030561 | 0,441431 | FALSE | trend |
| PPP1R9B | decreased | Q6R891 | Neurabin-2 OS=Mus musculus GN=Ppp1r9b PE=1 SV=1 | TRUE | TRUE | NMDA - control | supernatant | fdrtool | 0,013944 | 0,202744 | FALSE | trend |
| PPP2R5D | decreased | Q91V89 | Protein Ppp2r5d OS=Mus musculus GN=Ppp2r5d PE=1 SV=1 | FALSE | TRUE | NMDA - control | pellet | fdrtool | 0,056132 | 0,543873 | FALSE | trend |
| PPT1 | decreased | O88531\|Q3U6J9 | Palmitoyl-protein thioesterase 1 OS=Mus musculus GN=Ppt1 PE=1 SV=2\|Palmitoyl-protein thioester | TRUE | TRUE | NMDA - control | supernatant | fdrtool | 0,002537 | 0,060899 | FALSE | candidate |
| PRDX6 | decreased | Q6GT24 | Peroxiredoxin 6 OS=Mus musculus GN=Prdx6 PE=1 SV=1 | TRUE | FALSE | NMDA - control | supernatant | fdrtool | 0,059922 | 0,51761 | FALSE | trend |
| PRKAR2A | decreased | A0A0A6YX73\|P12367\|Q8K1M3 | cAMP-dependent protein kinase type II-alpha regulatory subunit OS=Mus musculus GN=Prkar2a PE= | TRUE | TRUE | NMDA - control | pellet | fdrtool | 0,037076 | 0,476014 | FALSE | trend |
| PRODH | decreased | Q9WU79 | Proline dehydrogenase 1, mitochondrial OS=Mus musculus GN=Prodh PE=1 SV=2 | TRUE | TRUE | NMDA - control | supernatant | fdrtool | 0,002779 | 0,064515 | FALSE | candidate |
| PSAT1 | decreased | Q543K5\|Q99K85 | Phosphoserine aminotransferase OS=Mus musculus GN=Psat1 PE=1 SV=1\|Phosphoserine aminotran | TRUE | FALSE | NMDA - control | supernatant | fdrtool | 0,031449 | 0,360264 | FALSE | trend |
| PSMA5 | decreased | Q3UPK6\|Q9Z2U1 | Proteasome subunit alpha type OS=Mus musculus GN=Psma5 PE=1 SV=1\|Proteasome subunit alpha | TRUE | FALSE | NMDA - control | supernatant | fdrtool | 0,07571 | 0,575499 | FALSE | trend |
| PSMD3 | decreased | P14685 | 26S proteasome non-ATPase regulatory subunit 3 OS=Mus musculus GN=Psmd3 PE=1 SV=3 | TRUE | TRUE | NMDA - control | pellet | fdrtool | 0,067082 | 0,569663 | FALSE | trend |
| PVALB | decreased | P32848\|Q545M7 | Parvalbumin alpha OS=Mus musculus GN=Pvalb PE=1 SV=3\|Parvalbumin alpha OS=Mus musculus GI | TRUE | FALSE | NMDA - control | supernatant | fdrtool | 1,55E-05 | 0,001217 | TRUE | hit |
| PYGB | decreased | Q8CI94 | Glycogen phosphorylase, brain form OS=Mus musculus GN=Pygb PE=1 SV=3 | TRUE | FALSE | NMDA - control | supernatant | fdrtool | 0,069105 | 0,553062 | FALSE | trend |
| Q3UNZ8 | decreased | Q3UNZ8 | Quinone oxidoreductase-like protein 2 OS=Mus musculus PE=1 SV=1 | TRUE | FALSE | NMDA - control | supernatant | fdrtool | 0,048157 | 0,463041 | FALSE | trend |
| Q8C3W1 | decreased | Q8C3W1 | Uncharacterized protein C1orf198 homolog OS=Mus musculus PE=1 SV=1 | TRUE | FALSE | NMDA - control | supernatant | fdrtool | 0,037799 | 0,403645 | FALSE | trend |
| RALGAPA1 | decreased | Q6GYP7\|Q6GYP7-2 | Ral GTPase-activating protein subunit alpha-1 OS=Mus musculus GN=Ralgapa1 PE=1 SV=1\|Isoform 2 | FALSE | TRUE | NMDA - control | pellet | fdrtool | 0,000127 | 0,033938 | TRUE | hit |
| RAP1GAP | decreased | A2ALS4\|A2ALS5\|A2ALS5-2\|A2ALS5-3\|A2ALS5-4 | Isoform 2 of Rap1 GTPase-activating protein 1 OS=Mus musculus GN=Rap1gap\|Isoform 3 of Rap1 GT | TRUE | TRUE | NMDA - control | pellet | fdrtool | 0,001958 | 0,11153 | FALSE | candidate |
| REEP5 | decreased | G3X8R0 | Receptor expression-enhancing protein OS=Mus musculus GN=Reep5 PE=1 SV=1 | FALSE | TRUE | NMDA - control | pellet | fdrtool | 0,041919 | 0,497123 | FALSE | trend |
| ROCK2 | decreased | F8VPK5\|P70336\|P70336-2 | Rho-associated protein kinase OS=Mus musculus GN=Rock2 PE=1 SV=1\|Rho-associated protein kina: | TRUE | TRUE | NMDA - control | pellet | fdrtool | 0,009236 | 0,231467 | FALSE | trend |
| RTCB | decreased | Q99LF4 | tRNA-splicing ligase RtcB homolog OS=Mus musculus GN=Rtcb PE=1 SV=1 | TRUE | TRUE | NMDA - control | pellet | fdrtool | 0,081234 | 0,594855 | FALSE | trend |
| RTN3 | decreased | Q9ES97\|Q9ES97-2\|Q9ES97-3 | Isoform 2 of Reticulon-3 OS=Mus musculus GN=Rtn3\|Isoform 3 of Reticulon-3 OS=Mus musculus GN | TRUE | TRUE | NMDA - control | pellet | fdrtool | 0,039005 | 0,48483 | FALSE | trend |
| RTN4 | decreased | Q99P72\|Q99P72-1\|Q99P72-3 | Isoform 2 of Reticulon-4 OS=Mus musculus GN=Rtn4\|Isoform 3 of Reticulon-4 OS=Mus musculus GN | TRUE | TRUE | NMDA - control | pellet | fdrtool | 0,05919 | 0,551779 | FALSE | trend |
| SARDH | decreased | Q3TWI2\|Q99LB7 | Sarcosine dehydrogenase OS=Mus musculus GN=Sardh PE=1 SV=1\|Sarcosine dehydrogenase, mitocl | TRUE | FALSE | NMDA - control | supernatant | fdrtool | 0,042154 | 0,430148 | FALSE | trend |
| SCP2 | decreased | P32020\|P32020-2 | Non-specific lipid-transfer protein OS=Mus musculus GN=Scp2 PE=1 SV=3\|Isoform SCP2 of Non-spec | TRUE | FALSE | NMDA - control | supernatant | fdrtool | 0,028258 | 0,335992 | FALSE | trend |
| SCRN1 | decreased | Q9CZC8 | Secernin-1 OS=Mus musculus GN=Scrn1 PE=1 SV=1 | TRUE | FALSE | NMDA - control | supernatant | fdrtool | 0,060811 | 0,521283 | FALSE | trend |
| SEPT3 | decreased | Q9Z1S5\|Q9Z1S5-2 | Neuronal-specific septin-3 OS=Mus musculus GN=Sept3 PE=1 SV=2\|Isoform 2 of Neuronal-specific se | TRUE | TRUE | NMDA - control | pellet | fdrtool | 0,050154 | 0,52647 | FALSE | trend |
| SH3GL2 | decreased | A2ALV3\|Q62420 | Endophilin-A1 OS=Mus musculus GN=Sh3gl2 PE=1 SV=1\|Endophilin-A1 OS=Mus musculus GN=Sh3gl | TRUE | TRUE | NMDA - control | pellet | fdrtool | 0,068188 | 0,571918 | FALSE | trend |
| SLC14A1 | decreased | Q8VHL0\|Q8VHL0-2 | Urea transporter 1 OS=Mus musculus GN=Slc14a1 PE=1 SV=2\|Isoform 2 of Urea transporter 1 OS=Mu | FALSE | TRUE | NMDA - control | pellet | fdrtool | 0,068874 | 0,573289 | FALSE | trend |
| SLC25A18 | decreased | Q9DB41\|Q9DB41-2 | Mitochondrial glutamate carrier 2 OS=Mus musculus GN=Slc25a18 PE=1 SV=4\|Isoform 2 of Mitochor | FALSE | TRUE | NMDA - control | pellet | fdrtool | 0,074659 | 0,584072 | FALSE | trend |
| SLC35A4 | decreased | A0A087WQH8 | Probable UDP-sugar transporter protein SLC35A4 OS=Mus musculus GN=Slc35a4 PE=1 SV=1 | FALSE | TRUE | NMDA - control | pellet | fdrtool | 0,008341 | 0,223981 | FALSE | trend |
| SLC39A12 | decreased | Q5FWH7\|Q5FWH7-2 | Zinc transporter ZIP12 OS=Mus musculus GN=Slc39a12 PE=1 SV=1\|Isoform 2 of Zinc transporter ZIP1 | FALSE | TRUE | NMDA - control | pellet | fdrtool | 0,015968 | 0,32037 | FALSE | trend |
| SNCB | decreased | Q91ZZ3 | Beta-synuclein OS=Mus musculus GN=Sncb PE=1 SV=1 | TRUE | TRUE | NMDA - control | supernatant | fdrtool | 0,053194 | 0,487846 | FALSE | trend |
| SNCB | decreased | Q91ZZ3 | Beta-synuclein OS=Mus musculus GN=Sncb PE=1 SV=1 | TRUE | TRUE | NMDA - control | pellet | fdrtool | 0,060866 | 0,555865 | FALSE | trend |
| SPECC1 | decreased | A0A0J9YTU3\|A0A0J9YUF8\|A0A0J9YUR2\|A0A0J9YV86\|Q5SXY1\|Q5SXY1-2\|Q5SXY1-3 | Cytospin-B OS=Mus musculus GN=Specc1 PE=1 SV=1\|Cytospin-B OS=Mus musculus GN=Specc1 PE=: | FALSE | TRUE | NMDA - control | pellet | fdrtool | 0,075317 | 0,585218 | FALSE | trend |
| SPTBN1 | decreased | Q62261 | Spectrin beta chain, non-erythrocytic 1 OS=Mus musculus GN=Sptbn1 PE=1 SV=2 | TRUE | TRUE | NMDA - control | pellet | fdrtool | 0,026932 | 0,418133 | FALSE | trend |
| SPTBN2 | decreased | Q68FG2 | Protein Sptbn2 OS=Mus musculus GN=Sptbn2 PE=1 SV=1 | TRUE | TRUE | NMDA - control | pellet | fdrtool | 0,002961 | 0,142298 | FALSE | candidate |
| STMN1 | decreased | P54227\|Q545B6 | Stathmin OS=Mus musculus GN=Stmn1 PE=1 SV=2\|MCG12955, isoform CRA a OS=Mus musculus GN | TRUE | FALSE | NMDA - control | supernatant | fdrtool | 0,017301 | 0,236723 | FALSE | trend |
| STRN | decreased | O55106 | Striatin OS=Mus musculus GN=Strn PE=1 SV=2 | FALSE | TRUE | NMDA - control | pellet | fdrtool | 0,031544 | 0,447193 | FALSE | trend |
| SUCLG1 | decreased | Q9WUM5 | Succinyl-CoA ligase [ADP/GDP-forming] subunit alpha, mitochondrial OS=Mus musculus GN=Suclg1 F | TRUE | TRUE | NMDA - control | pellet | fdrtool | 0,027898 | 0,424669 | FALSE | trend |
| SUCLG2 | decreased | Q9Z2I8\|Q9Z2I8-2 | Succinyl-CoA ligase [GDP-forming] subunit beta, mitochondrial OS=Mus musculus GN=Suclg2 PE=1 S' | TRUE | FALSE | NMDA - control | supernatant | fdrtool | 0,003233 | 0,070721 | FALSE | candidate |
| SYN1 | decreased | O88935 | Synapsin-1 OS=Mus musculus GN=Syn1 PE=1 SV=2 | TRUE | TRUE | NMDA - control | pellet | fdrtool | 0,057822 | 0,548317 | FALSE | trend |
| SYNJ1 | decreased | D3Z656\|E9Q7S0\|F7BQW7\|Q8CHC4 | Synaptojanin-1 OS=Mus musculus GN=Synj1 PE=1 SV=1\|Synaptojanin-1 OS=Mus musculus GN=Synj1 | TRUE | TRUE | NMDA - control | pellet | fdrtool | 0,00309 | 0,145588 | FALSE | candidate |
| TOMM5 | decreased | B1AXP6 | Mitochondrial import receptor subunit TOM5 homolog OS=Mus musculus GN=Tomm5 PE=3 SV=1 | FALSE | TRUE | NMDA - control | pellet | fdrtool | 0,042222 | 0,498339 | FALSE | trend |
| TPI1 | decreased | P17751 | Triosephosphate isomerase OS=Mus musculus GN=Tpi1 PE=1 SV=4 | TRUE | TRUE | NMDA - control | pellet | fdrtool | 0,059118 | 0,551599 | FALSE | trend |
| TPP1 | decreased | O89023 | Tripeptidyl-peptidase 1 OS=Mus musculus GN=Tpp1 PE=1 SV=2 | TRUE | FALSE | NMDA - control | supernatant | fdrtool | 7,83E-05 | 0,004841 | FALSE | candidate |
| TST | decreased | P52196\|Q545S0 | Thiosulfate sulfurtransferase OS=Mus musculus GN=Tst PE=1 SV=3\|Sulfurtransferase OS=Mus muscu | TRUE | TRUE | NMDA - control | supernatant | fdrtool | 0,077038 | 0,579744 | FALSE | trend |
| TST | decreased | P52196\|Q545S0 | Thiosulfate sulfurtransferase OS=Mus musculus GN=Tst PE=1 SV=3\|Sulfurtransferase OS=Mus muscu | TRUE | TRUE | NMDA - control | pellet | fdrtool | 0,048468 | 0,521022 | FALSE | trend |
| TWF1 | decreased | Q91YR1 | Twinfilin-1 OS=Mus musculus GN=Twf1 PE=1 SV=2 | FALSE | TRUE | NMDA - control | pellet | fdrtool | 0,048678 | 0,521716 | FALSE | trend |
| UBE2M | decreased | F6WMC0\|F7CDT0\|P61082 | NEDD8-conjugating enzyme Ubc12 (Fragment) OS=Mus musculus GN=Ube2m PE=1 SV=1\|NEDD8-cor | TRUE | TRUE | NMDA - control | pellet | fdrtool | 0,061142 | 0,55652 | FALSE | trend |
| UBE2O | decreased | Q6ZPJ3 | (E3-independent) E2 ubiquitin-conjugating enzyme UBE2O OS=Mus musculus GN=Ube2o PE=1 SV=3 | FALSE | TRUE | NMDA - control | pellet | fdrtool | 0,067952 | 0,571441 | FALSE | trend |
| VCP | decreased | Q01853 | Transitional endoplasmic reticulum ATPase OS=Mus musculus GN=Vcp PE=1 SV=4 | TRUE | TRUE | NMDA - control | pellet | fdrtool | 0,062273 | 0,559166 | FALSE | trend |
| WIPF3 | decreased | D3Z5R4\|E9Q2D0\|P0C7L0\|P0C7L0-2 | WAS/WASL-interacting protein family member 3 OS=Mus musculus GN=Wipf3 PE=1 SV=1\|WAS/WAS | FALSE | TRUE | NMDA - control | pellet | fdrtool | 0,066227 | 0,567882 | FALSE | trend |
