## Supplementary material for "Long-term synaptic depression triggers local biogenesis of autophagic vesicles in dendrites and requires autophagic degradation": table_S2

**Table_2**

| protein | hit annotation | UP/DOWN | ASD SCORE |
|---|---|---|---|
| TRIO | candidate | UP | 2 |
| SYT1 | hit | UP | S |
| SYNGAP1 | candidate | UP | 1S |
| SYN2 | trend | UP | 4 |
| SYN1 | candidate | UP | 4 |
| STXBP1 | candidate | UP | 3S |
| STX1A | trend | UP | 4 |
| SLC30A3 | trend | UP | 5 |
| SLC12A5 | trend | UP | 3 |
| SCN2A1 | trend | UP | 1 |
| RPS6KA2 | candidate | UP | 4 |
| RHEB | trend | UP | S |
| RGS7 | candidate | UP | 4 |
| RAB2A | candidate | UP | 3 |
| PSD3 | hit | UP | 4 |
| PRKCB | candidate | UP | 3 |
| PRKCA | trend | UP | 4 |
| PLCB1 | trend | UP | 3 |
| PACS1 | candidate | UP | S |
| NEFL | trend | UP | 5 |
| NDUFA5 | trend | UP | 4 |
| NCKAP1 | candidate | UP | 2 |
| MYO5A | hit | UP | 3 |
| MYH10 | trend | UP | 3 |
| MARK1 | hit | UP | 4 |
| MADD | hit | UP | 3 |
| KIF5C | trend | UP | 4S |
| KCNQ2 | trend | UP | 3 |
| IL1RAPL1 | trend | UP | 4 |
| ICA1 | trend | UP | 3 |
| GRIN1 | trend | UP | 3 |
| GAS7 | trend | UP | 1S |
| EXOC3 | trend | UP | 4 |
| DMXL2 | trend | UP | 4 |
| DLG4 | candidate | UP | 5 |
| DLG1 | candidate | UP | 4 |
| CTNNA2 | trend | UP | S |
| CAMK2B | candidate | UP | S |
| CAMK2A | hit | UP | 4S |
| CADPS2 | trend | UP | 4 |
| CACNA2D1 | trend | UP | 4 |
| CACNA1E | trend | UP | 3 |
| BRAF | candidate | UP | S |
| ATP8A1 | trend | UP | 5 |
| ATP1A3 | hit | UP | 4S |
| ATP1A1 | trend | UP | 4S |
| ANKS1B | trend | UP | 4 |
| ANK2 | candidate | UP | 1 |
| ADCY5 | trend | UP | 4 |
| SYNJ1 | candidate | DOWN | 4 |
| SUCLG2 | candidate | DOWN | 6 |
| SCP2 | trend | DOWN | 4 |
| PVALB | hit | DOWN | 5 |
| PRODH | candidate | DOWN | 3S |
| PPP2R5D | trend | DOWN | 4S |
| PCCB | trend | DOWN | S |
| NLGN3 | trend | DOWN | 2 |
| NF1 | trend | DOWN | S |
| MAP2 | hit | DOWN | 5 |
| HUWE1 | trend | DOWN | S |
| HSD11B1 | trend | DOWN | 4 |
| HBA | trend | DOWN | 1 |
| GPHN | candidate | DOWN | 3 |
| GLUD1 | candidate | DOWN | 3 |
| GLO1 | trend | DOWN | 4 |
| GAD1 | candidate | DOWN | 5 |
| FABP7 | hit | DOWN | 6 |
| FABP5 | candidate | DOWN | 4 |
| ETFB | candidate | DOWN | 3 |
| DPYSL3 | candidate | DOWN | 4 |
| DPYSL2 | hit | DOWN | 3 |
| DLG2 | trend | DOWN | NO SCORE |
| CTNNB1 | trend | DOWN | 3 |
| CLASP1 | trend | DOWN | 3 |
| CDC42BPB | trend | DOWN | 3 |
| CBS | trend | DOWN | 6 |
| CBLN1 | hit | DOWN | 5 |
| CAMSAP2 | trend | DOWN | 5 |
| BIN1 | candidate | DOWN | NO SCORE |
| ARHGAP32 | trend | DOWN | 4 |

S=Syndromic
1>2>3>4>5
